# The Alzheimer’s disease risk gene *ABCA7* regulates microglial lysosomal and lipid homeostasis and promotes phagocytosis to maintain neuronal health

**DOI:** 10.64898/2026.10.16.759681

**Authors:** Sarah E. Heuer, Fasih M. Ahsan, Gizem Terzioglu, Arnab Misra, Duc Duong, Chaitanya Katkar, Martin Giera, Rik van der Kant, Nicholas T. Seyfried, Tracy L. Young-Pearse

## Abstract

**Background:** Loss of function mutations in the lipid transporter ABCA7 have been linked to increased risk for late onset Alzheimer’s disease (AD). Although rare and common variants in *ABCA7* associated with AD alter the coding sequence and are predicted to impair protein function, the precise cell-type consequences of ABCA7 deficiency remain incompletely defined.

**Methods:** We used CRISPR/Cas9 gene editing to generate induced pluripotent stem cells (iPSCs) that were deficient for ABCA7 (ABCA7-KO) and their wild-type (ABCA7-WT) isogenic counterparts and subsequently performed unbiased proteomics on differentiated astrocytes (iAs), neurons (iNs) and microglia (iMGs). ABCA7 deficiency in iMGs produced the largest proteomic changes, so we therefore conducted additional functional analyses to define its effects on microglial lysosomal activity, phagocytosis, and lipid metabolism. In addition, we leveraged CRISPR/Cas9 to correct the ABCA7 rs3752246 AD risk variant in patient-derived iPSCs and compared proteomic and functional phenotypes between corrected iMGs and isogenic mutant carriers. Finally, we examined the functional consequence of ABCA7-deficient microglia on neurons.

**Results:** ABCA7-KO iMGs were deficient for proteins involved in lysosome function, measured by down-regulated lysosomal proteomic signatures, as well as reduced DQ-BSA activity and reduced LAMP1+ area. Additionally, ABCA7-KO iMGs displayed impaired phagocytic capacity and increased inflammatory markers, including MHC proteins CD1A and CD1E, and IL-18,. Multi-omic integration of lipidomic and proteomic profiles nominated a dysregulated metabolic network characterized by altered phospholipid steady-state metabolism and increased lipid peroxidation in ABCA7-KO iMGs. This was complemented by an accumulation of polyunsaturated triacylglycerols and lipid droplets, suggesting that ABCA7 deficiency disrupts intracellular lipid dynamics. This was supported by the alleviation of lysosomal and lipid droplet phenotypes through the inhibiting the synthesis of triacylglycerols. Concordantly, proteomic and functional analyses revealed lysosomal dysfunction in rs3752246-mutant iMGs, suggesting a shared ABCA7-dependent mechanism underlying AD risk. Tri-cultures of iNs, iAs and iMGs revealed that ABCA7 deficiency in iMGs worsened AD-relevant neuronal measures such as tau phosphorylation and synapse number.

**Conclusions:** This work supplies evidence for the important role of ABCA7 in microglia and provides unique insights into the molecular and functional consequences of reduced ABCA7 function in lysosomal and lipid regulation, paving a path forward to a more comprehensive understanding of AD molecular subtypes and personalized therapeutics.

## BACKGROUND

Alzheimer’s disease (AD) is a progressive neurodegenerative disorder and the leading cause of dementia worldwide. Individuals affected by AD often present in the clinic with reduced cognitive and memory function, with the accumulation of hallmark brain pathologies, such as extracellular beta amyloid (Aβ) plaques and intracellular tangles of hyperphosphorylated tau, confirming diagnosis. However, recent studies have shown that the age at onset, rate of progression, and severity by which neuropathological AD hallmarks arise is variable across affected individuals (1,2). While many factors contribute to this heterogeneity, genetic variation has been implicated as a critical factor due to the significant contribution of rare autosomal-dominant mutations in *APP*, *PSEN1* and *PSEN2* to familial AD (fAD) (3), and twin and family studies demonstrating a predicted heritability of 58-79% in the onset of the more common sporadic late-onset AD (LOAD) (4). Furthermore, recent human genome wide association studies (GWAS) have nominated over 70 loci with variants associated with LOAD risk (5–7). Many of these variants are common and have relatively low effect sizes, but are often localized to or around genomic regions encoding genes that are associated with biological processes altered in AD, including inflammation, proteostasis, and lipid metabolism (8). Elucidating the functions of genes associated with AD, and how their genetic variants impact their associated biological domains is key to understanding the disease and unlocking new and personalized therapies.

Both genetic and molecular studies have nominated the importance of lipid biology in contributing to AD risk. Along with Aβ plaques and tau tangles, Dr. Alois Alzheimer observed “lipoid granules” and lipid droplets as a key feature of the pathology of a brain from a patient with AD (9). Over a century after the original 1907 findings, lipid droplets are still a burgeoning research interest, as microglia laden with these pathological hallmarks are associated with increased parenchymal toxicity during aging and AD (10,11). In fact, lipids are intrinsically linked to the immunoregulatory activity of microglia, being linked to microglial effector functions such as phagocytosis and inflammation (12,13). Studies measuring lipid abundance have revealed that lipids change based on the progression of AD pathology, particularly phospholipids, sphingolipids and cholesterol (14–18). Genetic studies have also emphasized lipid biology as a crucial feature to AD risk since the E4 isoform of the lipid-binding protein APOE was discovered as the strongest genetic risk factor for developing LOAD (19–21). Compellingly, several genes bearing associated LOAD risk variants are also involved in lipid metabolism and transport, including the gene encoding the lipid transporter *ABCA7*.

The ATP-binding cassette, subfamily A, member 7 (*ABCA7*) gene is located on human chromosome 19 and encodes a transmembrane protein with a molecular weight of 220 kDa (22). The ABCA7 protein is a multi-pass transmembrane protein that forms a pore in the plasma membrane and utilizes both of its ATP binding domains to export lipids to the extracellular space in exchange for ATP hydrolysis (23). Prior biochemical and structural studies have implicated ABCA7 in phospholipid transport and efflux, including phosphatidylcholine, to extracellular lipid acceptors such as APOA-I and APOE (24–26). Single nuclear RNA sequencing (snRNAseq) datasets of the post-mortem human brain parenchyma show that *ABCA7* is expressed primarily in excitatory neurons, but it also has moderate nuclear RNA expression in microglia and astrocytes (27). While the functional role of ABCA7 in neurons has been more clearly elucidated to protect from metabolic stress and synaptic dysfunction (27–29), there are few studies interrogating the protein’s function in other cells of the brain parenchyma (30).

Several variants localized to *ABCA7* have been implicated with increased risk for developing AD, including mutations that cause changes to the protein-coding sequence (**Figure 1A**) (5,7,31,32). The most common variant, rs3752246, is a C>G single nucleotide polymorphism, altering the amino acid sequence at residue 1,527 from alanine to glycine (A1527G) (33). While the complete functional implications of this variant are unknown, recent work using computational modeling of protein conformational changes suggest that A1527G may drive a closed conformation of the protein, limiting the ability for lipids to be exported from the intracellular space (27). This variant is more predominant in individuals of European ancestry, and elicits a relatively low effect size on its own (5,7). Interestingly, a 44-base pair deletion in ABCA7 was recently discovered in individuals of primarily African descent, and heterozygous carriers of this mutation are at risk for developing AD almost equal to that of an individual carrying a E4 allele of APOE (31). Since most AD risk mutations localized to ABCA7 are thought to impair the function of the protein, it is crucial to gain a comprehensive understanding of the functional outcomes of ABCA7 deficiency using cell-intrinsic AD-relevant model systems. Furthermore, ABCA7 shares 79% of its protein-coding sequence between human and mouse, which is lower compared to conservation of similar ABC transporters such as ABCA1 (95%) and ABCA4 (88%) (34), suggesting murine studies of this important transporter may be confounded by interspecies discrepancies. Therefore, studying the functional genetic and proteomic implication of AD genetic risk factors, like *ABCA7*, requires the employment of proper human-relevant cellular models.

**Figure 1:**
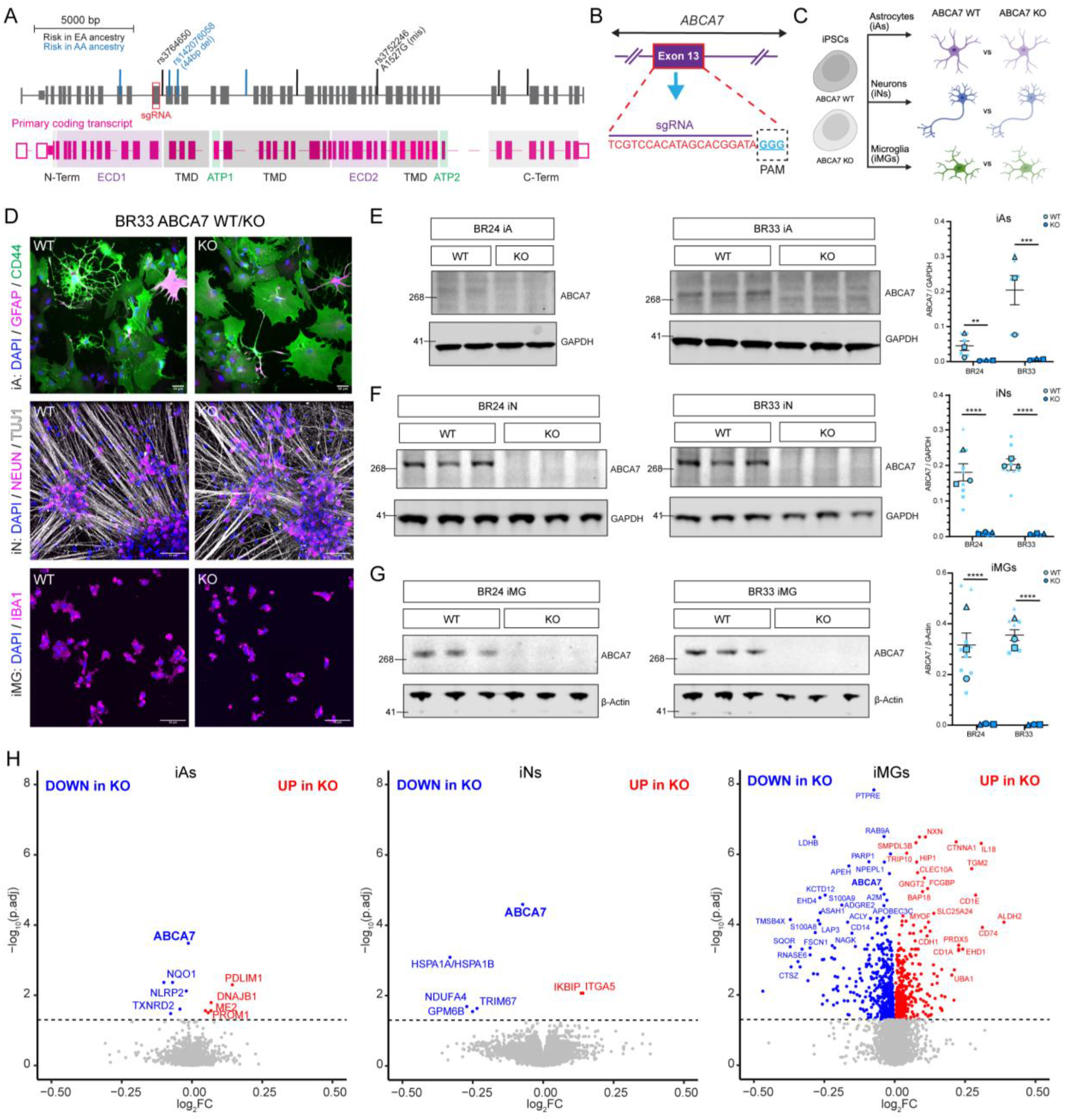
Microglia are sensitive to ABCA7-dependent proteomic alterations. (A) *ABCA7* gene structure and known AD risk variant locations. Exons are denoted as thick, grey rectangles; vertical black lines mark the location of risk variants predominant in European ancestries; vertical blue lines mark the location of risk variants predominant in African ancestries; red box relays the location of exon targeted for CRISPR/Cas9 to generate ABCA7 mutant iPSCs. (B) Schematic of sgRNA targeting ABCA7 exon 13 to generate mutant (ABCA7-KO) iPSCs. Red letters are the complete sgRNA sequence, and blue letters with dashed box denote the adjacent Cas9 PAM sequence. (C) Experimental design to generate iPSC-derived neurons (iNs), astrocytes (iAs) and microglia (iAs) from ABCA7-KO and isogenic WT counterparts for downstream analyses. (D) Representative immunocytochemistry of differentiated iAs (top), iNs (middle) and iMGs (bottom) from BR33 ABCA7-WT (left) and ABCA7-KO (right) iPSCs. Scale bars = 50µm. See also Figure S1 for ICC of BR24 differentiated cell-types. (E) Representative western blots from BR24 (left) and BR33 (middle) WT and KO iAs probing for ABCA7 and GAPDH. Quantification (right) for n=3 differentiations (large points with outline) and n=3 wells per differentiation (small points). Left numbers denote relative molecular weights of protein standard (kDa). (F) Representative western blots from BR24 (left) and BR33 (middle) WT and KO iNs probing for ABCA7 and GAPDH. Quantification (right) for n=3 differentiations (large points with outline) and n=3 wells per differentiation (small points). Left numbers denote relative molecular weights of protein standard (kDa). (G) Representative western blots from BR24 (left) and BR33 (middle) WT and KO iMGs probing for ABCA7 and β-actin. Quantification (right) for n=3 differentiations (large points with outline) and n=3 wells per differentiation (small points). Left numbers denote relative molecular weights of protein standard (kDa). (E-G) Western blot statistical analyses are results of mixed-effects ANOVA on raw non-WT-normalized data to capture within- and between-differentiation variance. Significant effect of genotype was determined: p<0.05 = *, p<0.01 = **, p<0.001 = ***, p<0.0001 = ****. Error bars = mean +/- SEM. (H) Volcano plots depicting differences in proteomic abundance between ABCA7-WT and -KO iAs (left), iNs (middle) and iMGs (right). Proteins were analyzed by their log_2_fold-change expression between WT and KO samples (x-axis) and statistical significance of change (-log_10_ adjusted p- value) determined using LIMMA analysis. Horizontal dashed line denotes log_10_ adjusted p-value < 0.05 significance threshold. Blue and red points are proteins significantly downregulated and upregulated in KO compared to WT samples, respectively.

We leveraged donor-derived human induced pluripotent stem cells (iPSCs) to identify the molecular and cellular consequences of ABCA7 deficiency in differentiated cells of the brain parenchyma: neurons (iNs), astrocytes (iAs) and microglia (iMGs). Using unbiased proteomics, we show that ABCA7 deficiency strongly impacts human iPSC-derived microglia, increasing immunological signaling proteins such as IL-18 and CD1A and negatively affecting lysosomal protein abundance. We validate that ABCA7 deficiency impairs microglial lysosomal homeostasis using orthogonal functional approaches, and negatively impacts phagocytosis with a reciprocal accumulation of lipid droplets. Further, perturbation of the phagocytic receptor TREM2 and inhibition of triacylglycerol formation via pharmacological co-manipulation of diglyceride acyltransferase 1 and 2 rescued deficits in lysosomal function. With the observed ABCA7-dependent impacts on lipid droplet accumulation, we integrated feature-rich proteomics data with unbiased lipidomics analysis to find that loss of ABCA7 in microglia alters phospholipid metabolism possibly by altering phosphatidic acid signaling and inducing associated oxidative stress responses such as intracellular lipid peroxidation. To test the AD-relevance of microglial ABCA7 deficiency, we show that the common rs3752246 AD risk variant in ABCA7 similarly impairs lysosomal homeostasis in human iPSC-derived microglia, proposing a mechanism by which ABCA7 loss promotes microglia dysfunction in AD. Finally, we provide evidence that microglial ABCA7 deficiency negatively impacts AD-related neuronal phenotypes such as Tau phosphorylation and synaptic colocalization in a parenchymal tri-culture microphysiological environment. These changes are coupled to a reduction in the extracellular lipid transporting proteins APOE and CLU. Together, we demonstrate that loss of ABCA7 impacts several crucial cell-autonomous functions for microglia, possibly stemming from a dysregulation of intracellular lipid dynamics. Microglial ABCA7 deficiency also has non-cell autonomous impacts on cells of the parenchymal microenvironment, nominating therapeutic avenues for correcting lipid-related microglia dysfunction in AD.

## METHODS

### Cell Culture

#### Human induced pluripotent stem cell (iPSC) lines

All work with human iPSC lines was performed following IRB review and approval through MGB/BWH IRB (#2015P001676). iPSCs were generated from cryopreserved peripheral blood mononuclear cell (PBMC) samples from autopsied participants from the ROS and MAP cohorts. iPSCs were generated using Sendai reprogramming method (35). iPSCs undergo a rigorous quality procedure that includes a sterility check, mycoplasma testing, karyotyping, and pluripotency assays performed by the New York Stem Cell Foundation (NYSCF). iPSCs were maintained using StemFlex Medium (ThermoFisher, A3349401). All cell lines were routinely tested for mycoplasma using PCR kit (MP0035-1KT) and short-tandem repeat (STR) profiling to prevent potential contamination or alteration to the cell lines.

#### CRISPR targeting to generate ABCA7 knockout iPSCs

We previously generated a cohort of iPSC lines from individuals in the ROSMAP cohorts, and these lines were confirmed to have normal karyotype, appropriately express pluripotent markers, and generate all germ layers upon differentiation (36). Here, we generated ABCA7 knockout (KO) and isogenic wild-type (WT) iPSC lines using methods as previously described (37,38) using a male and female ROSMAP line derived from healthy control individuals (BR24 and BR33), both homozygous for *APOE3/3* alleles. Briefly, guide RNAs (sgRNA) were designed using the Broad Institute CRISPick sgRNA design tool (39,40). The guide RNAs were ligated into plasmid backbone (pXPR_003, Addgene) and the sgRNA plasmid along with a plasmid containing Cas9 (pLX_311-Cas9, Addgene) were transfected into iPSCs using the Lonza P3 Primary Cell 4D-Nucleofector. After transfection, cells were selected using puromycin and blasticidin for 24 hours. Surviving cells were isolated as monoclones, and Sanger sequencing was performed to identify *ABCA7* mutations. Within these targeted genetic backgrounds (BR24 and BR33), one unedited/WT clone and one KO clone were chosen for downstream analysis. Successful KO clones were selected if a mutation was induced that would cause a coding frame shift that resulted in nonsense-mediated decay of the resulting transcripts (**Fig S1**). Off-target effects were evaluated by identifying coding regions that required 3-4 nucleotide mismatches to become identical to the gRNA sequence. The tables below outline all primer sequences for sgRNAs and sequencing for mutation confirmation and analysis of CRISPR off-target effects.

#### sgRNA Primers used for CRISPR

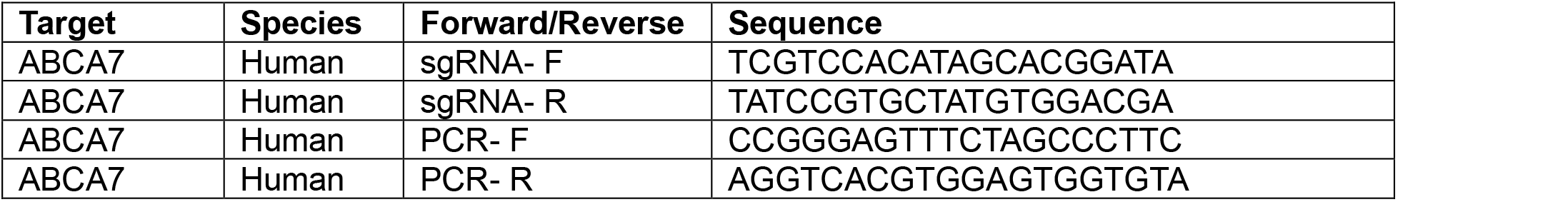

#### Primers used to evaluate off-target CRISPR effects

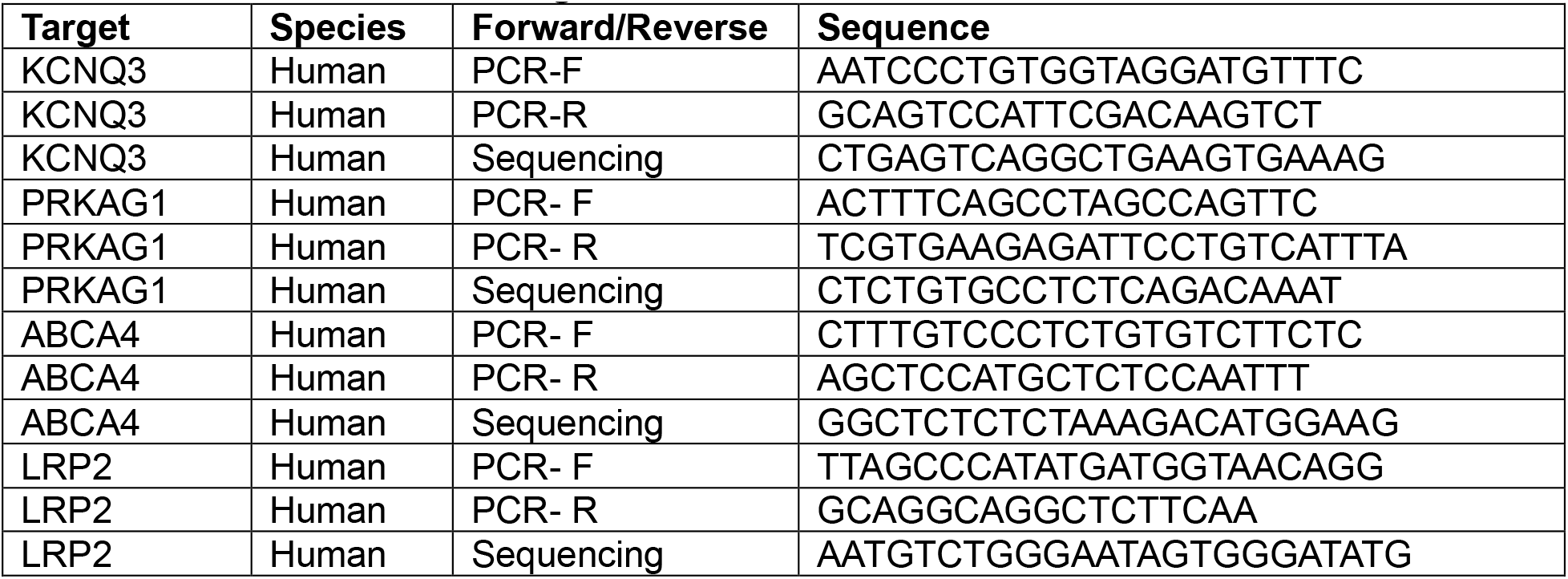

#### CRISPR targeting to correct the LOAD-associated rs372246 ABCA7 variant in iPSCs

We identified a ROSMAP participant that was homozygous for the rs3752246 Ala1527Gly coding variant associated with AD risk (BR103). This donor received a clinical and neuropathological diagnosis with AD and was homozygous for the neutral *APOE3/3* alleles. We previously had generated iPSCs from PBMCs derived from this donor (36), and here performed CRISPR targeting leveraging homology-directed repair by providing template DNA to alter the rs3752246 variant major allele. We designed sgRNAs and the template DNA sequencing using the Alt-R CRISPR HDR Design Tool provided by Integrated DNA Technologies (IDT). The sgRNAs were ligated into the eSpCas9(1.1) plasmid backbone (Addgene #71814), and transfected into iPSCs with 4uM of the corrected donor template DNA using the Lonza P3 Primary Cell 4D-Nucleofecto kit (Lonza #V4XP-3024) and Lonza 4D-Nucleofector X (Lonza #AAF-1003X) and Core Unit (Lonza #AAF-1003B). One day after transfection cells were selected using puromycin, and 24 hours after puromycin was added surviving cells were isolated for monoclonal selection. Sanger sequencing was performed using primers that amplified the rs3752246 region in ABCA7, and one edited minor allele (C/C genotype) and isogenic unedited major allele (G/G genotype) were chosen for downstream analyses. The table below outlines all sequences for sgRNAs, PCR primers, and HDR template DNA.

#### sgRNA Primers used for BR103 HDR-CRISPR

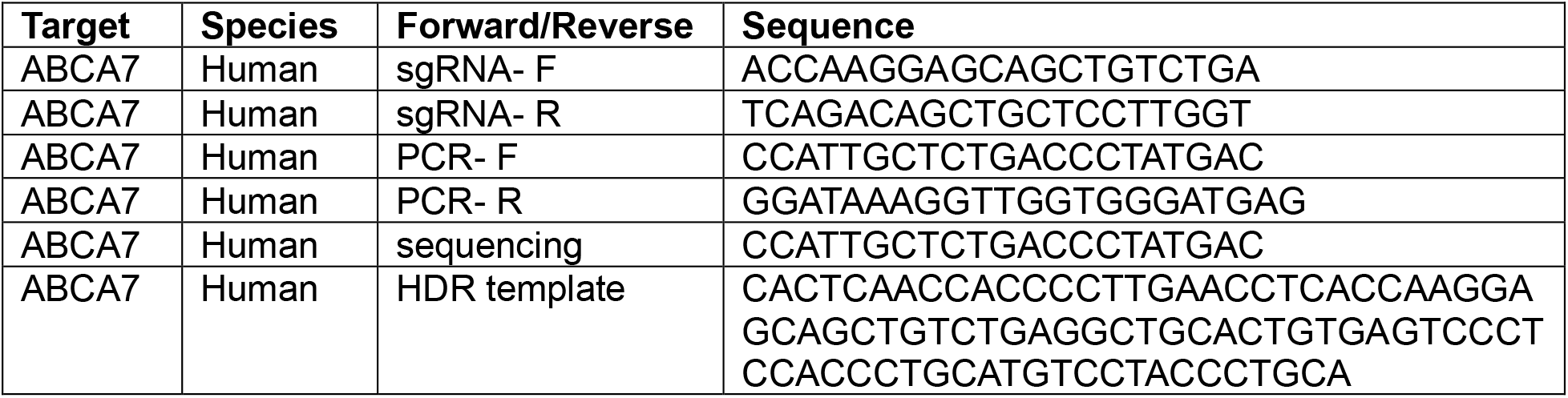

#### Differentiation of iPSCs into induced astrocytes (iAs)

iPSC-derived astrocytes (iAs) were generated following a published protocol (41), following minor modifications that have been previously described (37,38,42). iPSCs were plated at 95,000 cells/cm^2^ on growth factor reduced Matrigel (Corning #354230) coated plates prior to virus transduction. iPSCs were transduced 24 hours later with three lentiviruses: Tet-O-SOX9-puro (Addgene plasmid #117269), Tet-O-NFIB-hygro (Addgene plasmid #117271), and FUdeltaGW-rTa (Addgene plasmid #19780). The cells were then replated at 200,000 cells/cm^2^ using StemFlex medium and 10mM ROCK inhibitor (D0). The media was changed daily with Expansion Media (EM) from D1 to D3, then gradually switched from EM to FGF media from D4 to D7. On D8, cells were dissociated using 1:1 diluted Accutase and plated at 84,000 cells/cm^2^ using FGF media. Doxycycline (2.5ug/mL, Gibco) was added from D1 to the end of the differentiation, puromyclin (1.25mg/mL, Gibco) was added on D3 and D4 of the differentiation, and hygromycin (100mg/mL, Invivogen #ant-hg-1) was added from D4-D6 of the differentiation. From D8 to the end of the differentiation D21-22, cells were cultured with maturation media and fed every 2-3 days. Media formulations are listed below:

- Expansion Media (EM): DMEM/F12 (ThermoFisher), 10% Fetal Bovine Serum (FBS), 1X N2 Supplement (ThermoFisher), 1X GlutaMAX (Life Technologies)
- FGF Media: Neurobasal media (ThermoFisher), 1X B27 (ThermoFisher), 1X NEAA (Life Technologies), 1X GlutaMAX, 1% FBS, 8ng/mL FGF2 (Peprotech #100-18B), 5ng/mL CNTF (Peprotech #450-13), 10ng/mL BMP4 (Peprotech #120-05et)
- Maturation Media: 1:1 DMEM/F12 and Neurobasal media, 1X N2, 1X GlutaMAX, 1X Sodium Pyruvate (Life Technologies), 5ug/mL N-N-acetyl cysteine (Sigma A7250), 5ng/mL heparin-binding EGF-like GF (Sigma E4643), 10ng/mL CNTF, 10ng/mL BMP4, 500ug/mL dbcAMP (Sigma D0627)

#### Differentiation of iPSCs into induced neurons (iNs)

iPSC-derived neurons (iNs) were differentiated following a previously published protocol (43) with minor modifications we have described previously (36–38). iPSCs were plated at 95K cells/cm^2^ on plates coated with growth factor reduced Matrigel one day prior to virus transduction. 24 hours later, iPSCs were transduced with three lentiviruses: pTet-O-NGN2-puro (Addgene plasmid #52047, a gift from Marius Wernig), Tet-O-FUW-EGFP (Addgene plasmid #30130, a gift from Marius Wernig), and FUdeltaGW-rtTA (Addgene plasmid #197080, a gift from Konrad Hochedlinger). The cells were then replated at 200,000 cells/cm^2^ using StemFlex Medium and 10mM ROCK inhibitor (D0). The media was changed to KSR media (D1), 1:1 of KSR and N2B media (D2) and N2B media (D3). On D4, cells were dissociated using 1:3 diluted Accutase, and plated at 50,000 cells/cm^2^ using iN D4 media (NBM media + 1X B27 + 10ng/mL BDNF, GDNF, CNTF (Peprotech)) on non-growth factor reduced Matrigel (Corning #354277). Doxycycline (2ug/mL, Sigma) was added from D1 to the end of the differentiation, and puromycin (5mg/mL, Gibco) was added from D2 to the end of the differentiation. On D3, 0.5X B27 supplement was added. From D4 to the end of the differentiation on D21, cells were cultured in iN D4 media and fed every 2-3 days. Media formulations are listed below:

- KSR media: Knockout DMEM, 15% KOSR, 1X NEAA, 55µL beta-mercaptoethanol, 1X GlutaMAX.
- N2B media: DMEM/F12, 1X GlutaMAX, 1X N2 supplement B (StemCell Technologies), 0.3% dextrose (D-(+)-glucose, Sigma)
- NBM media: Neurobasal medium, 0.5X NEAA, 1X GlutaMAX, 0.3% dextrose (D-(+)-glucose, Sigma)

#### Differentiation of iPSCs into induced microglia (iMGs)

iPSC-derived microglia (iMGs) were differentiated following a previously published protocol (44) with minor modifications (45–47). iPSCs were plated on growth factor reduced Matrigel (Corning #354230) using StemFlex Medium (ThermoFisher) and 10mM ROCK inhibitor. From D0 to D12 of differentiation, StemDiff Hematopoietic Kit (StemCell Technologies) was used to generate hematopoietic progenitor cells (HPCs). On D12, cells were re-plated at 10,000 cells/cm^2^ onto phenol red-free growth factor reduced Matrigel (Corning) coated plates in iMG media supplemented with 3 growth factor cytokines: 100ng/mL IL34 (PeproTech #200-34), 50ng/mL TGFβ1 (Militenyi Biotech #130-108-969), and 25ng/mL M-CSF (ThermoFisher #PHC9501). From D12-D20 iMG media with freshly added cytokines was added to the culture every other day. On D20 cells were frozen in BamBanker and later thawed in fresh iMG medium at 100,000 cells/well in a 24 well plate coated with phenol red-free growth factor reduced Matrigel. From D20 to D37 iMG media with freshly added IL34, M-CSF and TGFβ1 was added to the culture every other day through ½ media changes. On D37 cells were dissociated with PBS at room temperature for 5 minutes, counted, and re-plated onto final experimental plates. Starting at D37, the iMG media was supplemented with two additional cytokines: 100ng/mL CD200 (Bon Opus, BP004) and 100ng/mL CX3CL1 (PeproTech #300-31). From D37 onwards iMG media with freshly added IL34, M-CSF, TGFβ1, CD200 and CX3CL1 was added to the culture every other day through ½ media changes until cells were harvested for experiments. Formulations of medias are listed below:

- iMG media: phenol red-free DMEM/F12 (ThermoFisher), 2X insulin-transferring-selenite (ThermoFisher), 2X B27, 0.5X N2, 1X GlutaMAX, 1X NEAA, 400mM monothioglycerol (Sigma), 5mg/mL insulin (Sigma), 1X PenStrep (ThermoFisher)

#### iPSC-derived microglia-astrocyte-neuron tri-cultures

iPSC-derived iMG-iA-iN tri-cultures were assembled as previously described (38,47,48). iNs and iAs were differentiated until D20 and iMGs were differentiated to D40 as described above. Initially, iNs, iAs and iMGs were seeded at 53,000 cells/cm^2^. On D20 of iN differentiation, iAs were dissociated using 1:3 diluted accutase and re-plated on top of iNs cultures at a density of 47,000 cells/cm^2^ in iA replating media. On iN-D21, iMGs were dissociated and re-plated on top if iN/iA co-cultures at a density of 42,000 cell/cm^2^ in tri-culture media. Cultures were replenished through a ½ media change on tri-culture D3 and harvested on tri-culture D6. Media formulations are listed below:

- iA replating media: iA Maturation media (see formulation above), 1X B27, 10ng/mL BDNF, 10ng/mL GDNF, 2.5µg/mL Doxycycline, 10mM ROCK inhibitor
- Tri-culture media: BrainPhys Neuronal Medium (Stem Cell Technology #5792), NeuroCult SM1 Neuronal Supplement (StemCell Technology #5711), 100ng/mL IL34, 50ng/mL TGFβ1, 25ng/mL M-CSF, 100ng/mL CD200, 100ng/mL CX3CL1

### Data-Independent Acquisition Mass Spectrometry (DIA-MS) Proteomics

#### Sample Preparation and Processing

Paired BR24 and BR33 iA, iN and iMG monocultures were differentiated as described above. iAs were plated at 120,000 cells/well onto a 12-well plate, and 3 wells were pooled together for an experimental replicate. iNs were plated at 200,000 cells/well onto a 24-well plate, with each well considered an experimental replicate. iMGs were plated at 100,000 cells/well on a 24-well plate on D37, with each well considered an experimental replicate. Cell pellets were collected and frozen at -80°C. Samples were sent to the Emory Integrated Proteomics Core (Emory University) for peptide sequencing and identification. Cell Pellets were lysed with 100µL EasyPep lysis buffer (ThermoFisher #A45735), and a Bicinchoninic Acid (BCA) assay was performed to determine protein concentration. Sample volumes equivalent to 15µg were obtained and reduced with 10mM Tris(2-carboxyethyl)phosphine (TCEP) and 40mM 2-chloroacetamide (CAA) at 90C for 10 min in the dark. Samples were digested with 1µg Lysl endopeptidase (Wako #125-05061) and 1µg trypsin (ThermoFisher #90057) overnight at room temperature. Digestion mixtures were cleaned with Water’s 10mg MCX plates (Fisher #50-780-153) according to manufacturer’s protocol, and eluents dried in a speedvac.

#### Liquid Chromatography and Mass Spectrometry

Each sample was resuspended in 20µL of loading buffer (0.1% formic acid) and 1µL was analyzed by liquid chromatography coupled to tandem mass spectrometry. Peptide eluents were separated by a Vanquish Neo (ThermoFisher) on a 15cm x 150µM internal diameter (ID) silica column packed with Dr. Maisch 1.5µm resin. Buffer A was water with 0.1% (vol/vol) formic acid, and buffer B was 100% (vol/vol) acetonitrile in water with 0.1% (vol/vol) formic acid. Elution was performed over a 14-minute gradient ranging from 3% to 50% solvent B. Peptides were monitored on an Orbitrap Astral mass spectrometer (ThermoFisher) fitted with a high-field asymmetric waveform ion mobility spectrometry (FAIMS Pro) ion mobility source (ThermoFisher). One compensation voltage (CV) of -35 was chosen for the FAIMS. Each cycle consisted of one full scan (MS1) with an m/z range of 380-980 at 240,000 resolution, 500% Automatic Gain Control (AGC) and 3 millisecond injection time. The higher energy collision-induced dissociation (HCD) data-independent acquisition (DIA) scans (MS2) were collected with a 2 m/z isolation window over the entire precursor range (380-980 m/z) with a time of 0.6 seconds and 3 millisecond injection time. Collision energy was set to 25% and scan range set to 150-2000 m/z.

#### Data Processing

Quantitative peptide measurements were acquired from the fragment ion MS2 spectra and aligned to known proteomic sequences. DIA-MS proteomics was analyzed separately for each cell-type. Proteomics data were first filtered for low-abundance proteins that were not present in more than 25% of the samples. Data were then batch regressed using ComBat (49), and voom normalized followed by calculation of differential expression using the LIMMA package in R (50). Proteins were considered significantly differentially expressed if the adjusted p-value < 0.05. Raw data are available at www.synapse.org under the following IDs: syn64759236 (ABCA7-WT/KO iA, iN, iMG), syn71758938 (BR103 iMG).

#### Gene Set Enrichment Preranked Analysis of iMGs

Ranking scores were calculated within the iMG LIMMA results by multiplying the log_2_FC of the protein by the adjusted p-value (adj.P) for BR24 and BR33 iMG, or nominal p-value for BR103 iMG. Data were then ranked in descending order, with the starting with proteins with the highest/most positive ranking score descending to proteins with the lowest/most negative ranking score. The pre-ranked gene/protein list was imported into GSEA (v4.3.3), and GSEA preranked analysis was run using GO Biological Pathways, GO Molecular Functions, GO Cell Compartments, and KEGG pathway reference datasets, with 1000 permutations set for p-value computation. Gene lists from the Human Proteostasis Network (HPN) were obtained and custom gene sets were generated using the Class designation for each listed gene/protein (51,52), and preranked analyses performed exactly as stated above. Pathways were considered significantly enriched if they had an enrichment p-value < 0.05.

### Western blotting

Cells were lysed with RIPA lysis buffer (ThermoFisher #89900) with protease inhibitors (Complete mini protease inhibitor, Roche) and phosphatase inhibitors (phosphoSTOP, Roche) added fresh before the cell lysis. Cells were lysed for 30 minutes on ice before transferring lysates to microcentrifuge tubes. Cell debris was pelleted by centrifugation at 13,000 x for 10 minutes at 4°C. Cell lysate supernatant was collected and stored at -20°C until use. Cell lysates were prepared with 4X LI-COR loading buffer (Fisher Scientific #NC9779096) and 2.5% β-mercaptoethanol, centrifuged and incubated at 95°C for 10 minutes. One exception was western blots on lysates probed for ABCA7 were not boiled. Samples were resolved using Novex NuPAGE 4-12% Bis-Tris gels (ThermoFisher #NP0336BOX) and NuPAGE 1X MOPS-SDS or MES-SDS running buffer (ThermoFisher #NP0001). Gel electrophoresis was run at 145V for 10 minutes followed by 200V for 30-45 minutes. SeeBlue Plus 2 (ThermoFisher #LC5925) or HiMark (ThermoFisher #LC5699) pre-stained protein standard was used for evaluation of relative molecular weight. The gel was extracted and transferred to a nitrocellulose membrane by incubation with 20% methanol tris-glycine transfer buffer at 400mA for 2 hours. The transferred blot was blocked with either Odyssey blocking buffer (LI-COR #927-50100) or 5% milk dissolved in TBST (1XTBS + 1% Tween-20) for 1 hour at room temperature with gentle agitation, then incubated in primary antibody (diluted in blocking buffer) overnight at 4°C with agitation. Blots were washed 3X in TBST, then incubated with LI-COR (1:10,000 dilution) or HRP-conjugated (1:1000) secondary diluted in TBST for 1 hour at room temperature with agitation. After secondary, blots were washed twice with TBST, once with TBS, and stored in TBS until imaging. Blots with LI-COR secondaries were imaged on a LI-COR Odyssey machine. Blots with HRP-conjugated antibodies (primary or secondary) were eluted using SuperSignal West Femto Maximum Sensitivity Substrate (ThermoFisher #34095) and imaged on an Invitrogen iBright CL1500 imaging system. Blots were quantified using ImageJ software. A table of antibodies used for western blotting is detailed below:

| Target | Host Species | Company | Catalog # | Concentration |
| --- | --- | --- | --- | --- |
| ABCA7 | Mouse | Santa Cruz | sc525408 | 1:300 |
| GAPDH | Mouse | Proteintech | 60004 | 1:10,000 |
| β-Actin | Mouse | Proteintech | 66009-1-Ig | 1:10,000 |
| β-Actin | Rabbit | Abcam | ab8227 | 1:1000 |
| IL18 | Rabbit | Proteintech | 10663-1-AP | 1:10,000 |
| LC3B | Rabbit | Novus | NB1002220 | 1:1000 |
| TREM2 | Rabbit | Abcam | AB209814 | 1:1000 |
| CTSD | Rabbit | Proteintech | 21327-1-AP | 1:10,000 |
| GM2A | Rabbit | Proteintech | 10864-2-AP | 1:1000 |
| AT8 / phospho-Tau (Ser202, Thr205) | Mouse | ThermoFisher | MN1020 | 1:250 |
| Tau | Rabbit | Cell Signaling | 46687S | 1:2000 |

### Immunocytochemistry

4% paraformaldehyde (PFA, Sigma) was added directly to the media at a 1:10 dilution, and cells incubated at 37°C for 15 minutes. Media was then removed from the cells and 4% PFA was added and incubated at room temperature for 15 minutes. For tri-cultures, ½ of the media was removed from each well and a mixture of 8% PFA and 8% Sucrose was added at a 1:1 ratio directly to the cells and incubated at room temperature for 15 minutes. Media/PFA was then removed from the cells and wells were washed once with PBS. Cells were blocked in 2% donkey serum (Jackson Immunoresearch) and 0.1% Triton-X-100 (Sigma) in PBS for 1 hour at room temperature with agitation. Primary antibodies were diluted in fresh blocking buffer and cells were incubated with primary antibody cocktails overnight at 4°C. Cells were then washed three times with PBS and incubated with secondary antibodies diluted in PBS for 1 hour at room temperature with agitation, then washed again once with PBS. Cells were treated with DAPI counterstain (1:1000 dilution in PBS, Invitrogen #D1306) for 10 minutes at room temperature, followed by a final PBS wash. To visualize lipid droplets, BODIPY (ThermoFisher, #D3922) was added to the counterstain mixture with DAPI at a 1:1000 dilution. Images of iNs, iMGs and tri-cultures were taken on the Andor Dragonfly 600 Spinning Disk Confocal at 100x or 40x magnification with oil immersion with 1µm (40x) or 0.5µm (100x) z-stacks. Images of iAs were taken on the GE INCELL Analyzer 2200 Widefield High-Content Imager at 20x magnification with air immersion, and 9 images taken per well. Images for TREM2 and DGAT intervention experiments were taken on the Operetta CLS high-content imaging system with at 63x magnification with water immersion, with 10 z-steps acquired at 1µm distance, and 25 fields of view acquired per well. All imaging and analyses were performed blinded by genotype and/or treatment. Images were pre-processed in ImageJ with brightness and contrast adjustments applied consistently across all representative images. A table of antibodies used for immunocytochemistry is detailed below:

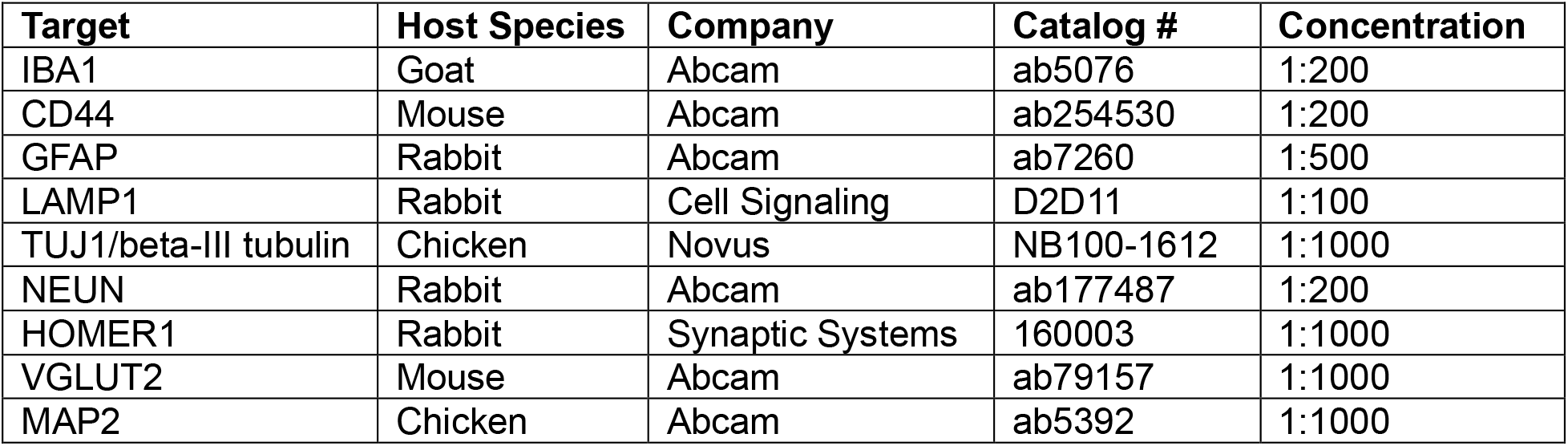

### Image analysis

#### General Image Analysis

ImageJ was used to generate maximum-intensity projections of individual channels from each image with 0.5/1 µm z-stack intervals. CellProfiler (53) was used to establish automated pipelines to measure signal area and intensity in confocal images. IdentifyPrimaryObjects was used to segment proteins of interest, IBA1+ microglia, CD44+ astrocytes, and NeuN+ cell bodies with DAPI+ nuclei. RelateObjects was used to segment whole cells containing nuclei and exclude debris. MaskObjects was used to remove any signal outside of cell bounds. MeasureImageAreaOccupied was used to measure area occupied by the proteins of interest and the cells, and MeasureObjectIntensity was used to quantify signal intensity.

#### Synapse colocalization

Six images per well of a 96-well plate were collected using a 40X (oil immersion) objective, and subsequently analyzed in CellProfiler (53) as previously described (38). Using the software’s primary object identification module, individual puncta corresponding to HOMER1 and VGLUT2 were detected. Co-localization analysis was performed by measuring the overlap between HOMER1 and VGLUT2 puncta, with an overall threshold of 80% as a proxy for synaptic sites. Additionally, puncta were required to be localized to TUJ1-marked neurites. For objects meeting both the co-localization and neurite criteria, the area of the puncta was quantified and normalized to the total neurite area.

### Enzyme-Linked Immunosorbent Assay (ELISA) kits for secreted protein measurements

All ELISAs were performed according to the manufacturer’s instructions:

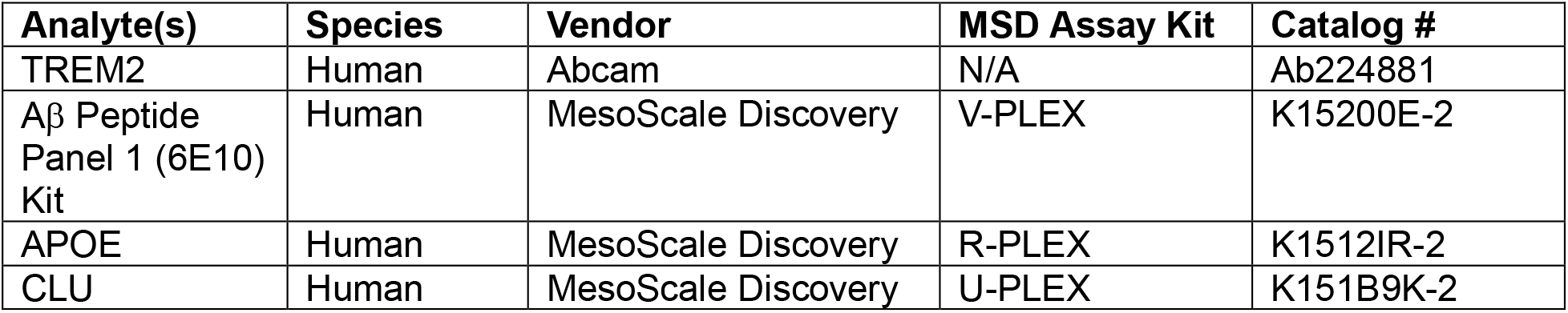

### Lysosomal measurements

#### DQ-Red BSA Assay

iMGs were incubated in 1 µg/mL DQ-Red BSA (Invitrogen, #d12051) for 1 hour at 37°C, after which the media was replaced with fresh iMG media for the chase period. Images were acquired hourly beginning at the start of the chase period, with 9 images per well acquired at 20x using the IncuCyte S3 Live-Cell Imaging System (Sartorius). Average DQ-BSA intensity was calculated by dividing the integrated DQ-Red BSA intensity by microglia area for each acquired image, measured by the IncuCyte S3 software (Sartorius). The average DQ-BSA intensity divided by microglia area was calculated as a normalized signal for each image and averaged across each well for a technical replicate. Normalized DQ-BSA intensities were plotted hourly post-chase to evaluate signal accumulation and stabilization over the time course experiment, with 2 hours identified as the peak stabilized time-point for cross-sectional analyses.

#### BSA endocytosis assay

To measure endocytosis, iMGs were incubated with 1 µg/mL BSA-Alexa-488 (Invitrogen #A13100) for 1 hour at 37°C. Negative control wells were pre-treated with 10 µM cytochalasin D (Sigma #C8273) for 30 minutes prior to the addition of BSA-Alexa 488. Cells were dissociated with PBS at room temperature for 5 minutes, then incubated with Zombie Violet (Biolegend #423113) at 1:1000 dilution for 10 minutes on ice. After washing and centrifugation, cells were stained with PE-Cy7-conjugated anti-CD11b antibodies in FACs buffer (HBSS + 20% FBS + 25mM HEPES + 2mM EDTA) and incubated on ice for 20 minutes. After centrifugation cells were fixed with 4% PFA for 10 minutes on ice, centrifuged, and resuspended in FACs buffer. Samples were analyzed for BSA-Alexa 488 fluorescence on a Fortessa flow cytometer equipped with 355nm, 488nm, 561nm and 640nm lasers. The mean fluorescence intensity of BSA-488 was quantified on live, CD11b+ gated populations. Data acquisition and analysis were performed using FlowJo v10.10.0 software, with compensation applied for multicolor fluorescence detection.

#### LysoTracker

Cells were incubated with 50nM LysoTracker Green (Invitrogen #L7526) for 1 hour at 37°C. Cells were dissociated at room temperature with PBS for 5 minutes, centrifuged at 500 x 5 for 5 minutes, then incubated in 1:1000 Zombie Violet for 10 minutes on ice. The cells were centrifuged again and resuspended in FACs buffer for flow cytometry measurements of Lysotracker fluorescence. The mean and median fluorescence intensity of Lysotracker green was quantified on live cell gated populations. Data acquisition and analysis were performed using FlowJo v10.10.0 software.

### Preparation of pHrodo-conjugated substrates for phagocytosis assays

#### Synaptosomes

Mice were anesthetized using isoflurane and euthanized following approved methods under BWH/MGB IACUC #2016N000464. Synaptosomes were isolated from mouse brains using Syn-PER (ThermoFisher #87793), following the manufacturer’s protocol as previously described(54,55) with minor modifications. Briefly, phosphatase and protease inhibitors were added to Syn-PER at a concentration of 1 tablet per 10mL. Mouse brain tissue was weighed, and 10mL of Syn-PER was added per gram of tissue. The tissue was Dounce homogenized on ice with 10-12 slow strokes, and the homogenate centrifuged at 1200 x g for 10 min at 4°C. The supernatant was collected and centrifuged at 15,000 x g for 20 minutes at 4°C to pellet the synaptosomes. The final pellet was resuspended in 0.1M sodium bicarbonate buffer (pH 8.3-8.4) at a concentration of 50 mg/mL. For pHrodo conjugation, the synaptosome suspension was incubated with 3µM of amine-reactive pHrodo Green STP dye (ThermoFisher #P35373) for 45 minutes at room temperature in the dark. The mixture was diluted 1:10 in PBS and centrifuged at 2500 x g for 5 minutes at room temperature. The pellet was washed twice with PBS resuspension followed by centrifugation, and the final pellet resuspended in phenol-free DMEM/F12 at a concentration of 50 mg/mL. Synaptosomes were provided to cells at a final concentration of 1 mg/mL.

#### Apoptotic Neurons

Apoptotic neurons (ANs) were generated from D14 iNs as previously described (54). Briefly, D14 iNs were subjected to ultraviolet (UV) radiation using an UV cross-linker at 500 Jm^-2^, and collected in PBS 24 hours later. ANs were centrifuged at 300 x g for 5 minutes, and the pellet weighed and resuspended in 0.1M sodium bicarbonate (pH 8.3-8.4) at 50 mg/mL. The cell suspension was conjugated with pHrodo using the methods described above. Following incubation period, the pellet was washed 3 times with PBS and centrifugation at 300 x g for 5 minutes. After the final wash cells were resuspended in 1mL PBS, counted, and provided to cells at 35,000 ANs/cm^2^.

#### Crude Myelin

Crude and purified myelin were prepared following previously published protocols (54,56) with minor modifications. Briefly, mice were anesthetized using isoflurane, euthanized following approved methods under BWH/MGB IACUC #2016N000464, and brains harvested. Brains were homogenized in 5mL 0.32M sucrose (Sigma) dissolved in PBS, and an additional 10mL 0.32M sucrose with protease and phosphatase inhibitors was added to the homogenate. A sucrose step gradient was performed by carefully overlaying the homogenate mixture on top of 15mL 0.85M sucrose in an ultracentrifuge tube. The gradient was centrifuged at 75,000 x g for 40 minutes at 4°C in a Beckman Coulter swinging bucket ultracentrifuge (SW41 Ti rotor). The gradient interface was collected using a Pasteur pipette, washed with MiliQ water, and centrifuged again at 75,000 x 6 for 15 minutes at 4°C. The supernatant was removed, and two osmotic shocks were delivered to the pellet by resuspending in sterile water and centrifuging at 12,000 x g for 15 minutes at 4°C. The final crude myelin pellet was weighed and resuspended in 0.1M sodium bicarbonate (pH 8.3-8.4) at 10mg/mL. Crude myelin aliquots could be stored at -80C for 6-12 months. For pHrodo conjugation, crude myelin was thawed on ice, and 10mM pHrodo was added at 1:1000 dilution and incubated for 45 minutes at room temperature covered. The mixture was centrifuged at 15,000 x g for 5 min at 4°C, and the pellet washed twice with PBS followed by centrifugation at the same speed and temperature. The final pellet was resuspended at 5mg/mL and applied to cells at 50ug/mL.

#### Purified Myelin

The preparation of purified myelin was performed identically to crude myelin until the osmotic shocks. After two osmotic shocks were provided to the crude myelin, the pellet was resuspended in 0.32M sucrose, and an additional step gradient was performed by layering on top of 0.85M sucrose in an ultracentrifuge tube. The gradient was centrifuged in a Beckman Coulter swinging bucket ultracentrifuge (SW41 Ti rotor) at 75,000 x g for 40 minutes at 4°C. The purified myelin gradient interface was collected and resuspended at 10mg/mL in 0.1M sodium bicarbonate (pH 8.3-8.4). Purified myelin aliquots could be stored at -80C for 6-12 months. The pHrodo conjugation was performed identically to the crude myelin conjugation described above and applied to cells at 50µg/mL concentration.

### Phagocytosis assay

D37 iMGs were plated at 30,000 cells/well onto 48 wells plates. 30 minutes prior to the addition of the pHrodo-conjugated substrate, iMGs were treated with 10µM cytochalasin D (Sigma #C8273) or equal volume DMSO. Cells were then exposed to pHrodo conjugated substrates overnight (12-18 hours). iMGs were gently dissociated with PBS for 5 minutes and centrifuged at 500 x g for 5 minutes at 4°C. Cells were stained with Zombie Violet (BioLegend) for viability discrimination and incubated on ice for 20 minutes, protected from light. After washing and centrifugation, cells were stained blocked with 7.5% FBS in FBS for 15 minutes, then stained with PE-Cy7-conjugated anti-CD11b antibodies in FACs buffer (HBSS + 20% FBS + 25mM HEPES + 2mM EDTA), and incubated on ice for 45 minutes. After centrifugation cells were fixed with 4% PFA for 10 minutes on ice, centrifuged, and resuspended in FACs buffer. Samples were analyzed on a Fortessa flow cytometer equipped with 355nm, 488nm, 561nm and 640nm lasers. Phagocytosis was quantified as the percentage of pHrodo-positive cells within live, CD11b+ gated populations. Data acquisition and analysis were performed using FlowJo v10.10.0 software, with compensation applied for multicolor fluorescence detection.

### Flow cytometric measurements of surface protein expression

On D37 iMGs were plated onto a 48wp at 30,000 cells per well. On D40, iMGs were gently dissociated and labeled for surface markers using the protocol above. Samples were analyzed on a Fortessa flow cytometer equipped with 355nm, 488nm, 561nm and 640nm lasers. The mean fluorescence intensity of identified surface markers was quantified on live, CD11b+ gated populations. Data acquisition and analysis were performed using FlowJo v10.10.0 software, with compensation applied for multicolor fluorescence detection. A detailed list of antibodies and concentrations are listed below:

| Antigen | Host | Dilution | Vendor | Catalog # |
| --- | --- | --- | --- | --- |
| CD11b | Rat | 1:200 | BioLegend | 101215 |
| TREM2 | Rat | 1:100 | R&D Systems | FAB17291P |
| CD1A | Mouse | 1:50 | BioLegend | 300106 |

### Treatments of cultures

To perturb lysosomal, phagocytic and/or lipid droplet phenotypes cells were provided with one of the following treatments before DQ-BSA, phagocytosis, or lipid droplet assays: Goat anti-Human TREM2 IgG (R&D Systems, #AF18281) or Normal Goat IgG control (R&D Systems, #AB-108-C) at 10µg/mL for 24 hours; AL002 (MedChemExpress, #HY-P991220) or Human IgG1 kappa isotype control (MedChemExpress, #HY-P99001) at 5µg/mL for 48 hours; combined DGAT1 PF-04620110 (MedChemExpress, #1109276-89-2) and DGAT2 PF-06424439 (MedChemExpress, #1469284-78-3) inhibitors or DMSO vehicle control both at 5µM for 18 hours.

### Unbiased Lipidomic Profiling

#### Sample Preparation

On D37, 5-6 wells per line of iMG were plated at 100,000 cells/well on a 24 well plate to acquire 500,000-600,000 iMG per ABCA7 genotype. On D40, microglia were dissociated at room temperature with PBS incubation for 5 minutes, followed by centrifugation at 500 x g for 5 minutes at 4°C. Cell pellets were frozen on dry ice and stored at -80°C until processing and analysis.

#### Flow-injection based Lipidomics

Lipidomics analysis was performed following standardized, quantitative protocols (57). Briefly, 25 µL of an internal standard mix containing more than 60 deuterated lipid standards (i.e. Avanti Ultimate SPLASH ONE mix with the addition of BMP, FA, CER, HexCER, LacCER and PA deuterated standards) was added to iMG pellets and lipids were extracted using a methyl tert-butyl ether–based protocol. Samples were dried under a gentle stream of nitrogen gas and were dissolved in running buffer (1:1 methanol:dichloromethane, containing 10 mM ammonium acetate) before injecting into the flow-injection platform, consisting of a SCIEX QTRAP 6500+ mass spectrometer equipped with SelexION DMS interface and a Shimadzu LC30 UHPLC-system. Shotgun Lipidomics Assistant (SLA) software was used to process data files and quantify lipid class, species concentration and composition values (58). Lipidomics data analysis was further accomplished using SODAlight as a built-in data browser for the Neurolipid atlas repository (18). Lipid species concentration data obtained from SLA output underwent preprocessing with blank filtering as follows: lipid species were retained for analysis if their signal intensity was at least twofold higher than blank in ≥80% of the samples; species absent or below threshold in >20% of samples were excluded. An exception was made for lipid species uniquely present in one experimental group, which were retained if detected above blank in ≥60% of samples within that group. No missing value imputation was performed. All further information on SLA control software can be found on (https://github.com/syjgino/SLA), and the code for SODAlight is available at (https://github.com/CPM-Metabolomics-Lipidomics/soda-light/). Raw data used for comparisons (in nmol/g) have been added to the Neurolipid Atlas repository (https://neurolipidatlas.com/).

#### Quantitative Lipidomics Analysis

Analysis of lipidomics data was performed using custom R scripts adapted from the LipidSigR (59) v1.0.3.0002 and Lipidr (60) v2.24.0 packages on the R v4.5.2 Bioconductor (61) v3.22 platform. Briefly, total sum and standard normalized lipid abundance values for three independent differentiations of ABCA7 WT/KO BR33 iMGs in technical duplicates (12 samples total) were tested for differential expression via an adjusted limma() function (50), treating the independent differentiation number of a given sample as a dependent covariate. Differentially abundant lipids (DALs) were calculated with a Benjamini-Hochberg (62) corrected P value < 0.05, and an absolute log_2_ fold change (log_2_FC) threshold > 1. Biophysical characteristics (bilayer thickness, headgroup charge, intrinsic curvature, lateral diffusion, and transition temperatures) for each lipid were extracted from the LIPID MAPS Ontology (LION) (63) v. 2023.04.14 database, with relative group enrichment tested between ABCA7 WT/KO via Lipid Set Enrichment Analysis (LSEA) using a t statistic generated ranked list, and subsequent visualization of the NES and log_10_ adjusted P value using ggplot v2.4.0.1. Principal components analysis (PCA) was performed using the prcomp() function. Lollipop plots displaying the log_2_FC between ABCA7 WT/KO was generated using DALs as described above, with stratification based on major lipid class and cellular compartment categorization via the LION database, as described above. Lipids were similarly assessed for triglyceride (TG) and phospholipid (PL) unsaturation, binning lipids based on TG and PL membership and visualizing the number of double bonds for a given lipid and their given log_2_FC.

Predicted ABCA7 perturbation lipid flux analysis was computed using a statistical strategy as previously described (64,65). Briefly, BioPAN (66) (Bioinformatics Methodology for Pathway Analysis) on the LIPID MAPS Lipidomics Gateway (67) was leveraged to generate an interaction network, binning quantitatively assessed lipids into major lipid class nodes and their respective edge interactions. For each BioPAN-defined individual reaction, we calculated the weighted ratio of product over reactant across genotypes 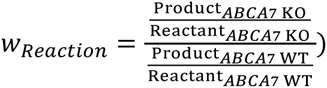 as a measure of predicted perturbation-mediated reaction activity. We then classified overall reaction activity as a measure of the t test between ABCA7 KO/WT for each weighted edge of the pathway, and generated a reaction activity Z score by converting the reaction P value through a cumulative distribution function [*Z_Reaction_* = *CDF*^−1^(1 − *P*)]. Z_Reaction_ scores were then aggregated into BioPAN-defined pathway level activity Z_Pathway_ scores via summation of all k number of reaction edges per pathway 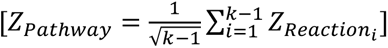 Pathways were subsequently defined as significantly active or suppressed with an absolute Z_Pathway_ > 1.645 (corresponding to a P value < 0.05). Network node and edge weights were visualized using Cytoscape11 v3.10.4 (68).

#### Integrated Proteomic/Lipidomic Network Analysis

Generation of integrated proteomic and lipidomic ABCA7 perturbation networks was performed using custom R scripts adapted from the LipidSigR (59) v1.0.3.0002, DEP2 (69) v0.5.28.2, and gatom (70,71) v1.8.4 packages on the R v4.5.2 Bioconductor (61) v3.22 platform. Briefly, DIA-MS MS2 proteomic abundance levels for 3 independent differentiations of ABCA7 KO/WT BR33 iMGs in technical duplicate (12 samples total) were extracted from our prior proteomic analyses (as described above in the methods section *Data-Independent Acquisition Mass Spectrometry (DIA-MS) Proteomics*), log2 transformed, and filtered for empty values in greater than 25% of all samples. Proteomics data were then variance stabilizing normalized (VSN), imputed using Quantile Regression Imputation of Left-Censored Data (QRILC) (72), and assessed for normality using the DEP2 impute() function. Differentially expressed proteins (DEPs) were identified using the DEP2 test_diff() function, leveraging the design formula “formula(∼ 0 + genotype + differentiation)”, and adjusting for multiple comparisons testing using the Benjamini-Hochberg (62) method. DEPs were identified with an adjusted P value < 0.05 and absolute log_2_FC > 1. The contribution of technical duplicates were assessed in the differential expression model using the *limma::duplicateCorrelation()* function and calculated highly-conserved adjusted p-values with both methods (lipidomics: Pearson = 0.9988; proteomics: Pearson = 0.9998). DALs and DEPs were subsequently integrated into a single protein-lipid reactant/product interaction network using a modified LipidSigR *nw_gatom()* function, with a connected subgraph of maximal weight identified using a signal GMWCS (generalized maximum weight connected subgraph) solver to generate a concordantly perturbed proteome-lipidome submodule (73). Protein-lipid interactions were connected in a metabolic network graph using the Rhea reaction database (74) where each node is a ChEBI labeled metabolite/lipid entity, and each edge is a shared enzymatic reaction between two entities sourced from Rhea, leveraging 11,886 unique reaction participants and evidenced by 15,500 unique PMID literature references. Node weight (size) is scaled by the −log_10_(p-value) of the connected feature’s differential abundance, while edge weight (width) is held as a fixed constant across the network and is not weighted by reaction confidence, flux magnitude, or any other measure. For GMWCS integration, we utilized the Rhea human reference network with node and edge inclusion capped at 3,000 genes and 3,000 lipids, a significance basis of BH-adjusted p-value at 0.05, a fold-change cutoff of 2-fold on the linear scale, the *mwcsr::rnc_solver()* for the underlying optimization, and a fixed random seed of 123 to ensure reproducibility. We further tested module stability across a small parameter grid, observing that the final enriched subgraph module is largely unchanged at a relaxed p-cutoff of 0.10 (Jaccard = 0.65 vs. baseline), to establish confidence in its stability across the chosen parameters. The network was subsequently visualized using Cytoscape v3.10.4, with heatmap visualization of DEPs/DALs of interest generated using the CRAN package pheatmap v1.0.13. Graph labels and aesthetics were lightly edited using Adobe Photoshop and Illustrator 2026 on the Adobe Creative Cloud Suite.

#### Reanalysis of FinnGen METSIM Genome-Wide Association Studies

Association signals for circulating plasma metabolites (1,391 compounds) linked to ∼26 million genetic variants identified in the FinnGen METSIM study of middle- and older- aged men (6,136 participants) recruited from northeast Finland were extracted from the original METSIM study at https://pheweb.org/metsim-metab (75,76). Genome-wide phenotype/association signals for C100015620: lactosyl-N-nervonoyl-sphingosine (d18:1/24:1) and C100009030: lactosyl-N-palmitoyl-sphingosine (d18:1/16:0) were extracted and visualized as a Manhattan plot using the R Bioconductor package topr (77) v2.0.2, with genome-wide significance P value thresholds set at 1e-10. Association signals for SNPs located in the ABCA7 genetic locus was similarly extracted and visualized using the topr function regionplot(). Population structure and systematic genotyping bias was ruled out in the METSIM dataset via Quantile-Quantile plot (QQ) visualization and subsequent calculation of λGC to quantify genomic inflation. (λ = 0.998 for association of both metabolite signals).

### Exogenous Lipid Release Assays

Exogenous lipid release assays were performed as previously described (24), with minor adaptations for cultured microglia. D40 microglia were loaded with 2µM Red-C12 (ThermoFisher, #D3835) or 2µM NBD cholesterol (ThermoFisher, #N1148) for 16 hours in complete media. After loading cells were placed in the IncuCyte S3 Live-Cell Imaging System, and 9 images per well were acquired every 2 hours at 20x magnification, collecting phase contrast, red (Red-C12) and green (NBD cholesterol) fluorescent channels. Media was replaced with warm HBSS (ThermoFisher, #14025134) which was then incubated for 4 hours at 37°C. HBSS was collected and analyzed for Red-C12 (excitation/emission 561/600 nm) or NBD cholesterol (excitation/emission 465/535) fluorescence using a Tecan Infinite M Plex plate reader. Intracellular lipid data was analyzed using integrated intensity of red (Red-C12) or green (NBD cholesterol) fluorescent signals restricted to cell masks identified using phase contrast. Extracellular lipid release was captured by measuring fluorescence intensity with background signal or signal from un-treated wells subtracted.

### Lipid Peroxidation Measurements

At D37 iMGs were plated on a glass-bottom 96 well plate at 15,000 cells/well. On D40 cells were treated with 2µM BODIPY 581/591 C11 (ThermoFisher #D3861) for 1 hour, then fixed in 4% PFA. ICC for IBA1 was performed on fixed cells using methods described above, and 6 images/well were acquired at 40x with 1µm z-steps on the Andor Dragonfly Spinning Disk confocal microscope. ImageJ was used to generate maximum-intensity projections of each channel. CellProfiler was used to establish an automated pipeline to measure signal area and intensity in confocal images. Lipid peroxidation was measured as the ratio of oxidized lipids with excitation/emission at 488/510 nm, and reduced lipids with excitation/emission at 581/591 nm. The signal was restricted to IBA1+ area to avoid any channel background noise.

### Statistical Analysis

Specific information regarding statistical analysis can be found in the figure legends. All statistical tests were performed using GraphPad Prism 10 on raw, non-WT-normalized data. All data is visualized as mean ± SEM and normalized to WT and/or vehicle treated conditions within each differentiation, unless otherwise stated. Statistical analysis for comparing two groups across multiple differentiations were performed using a two-way mixed-effects analysis. Data were arranged in a grouped format using individual technical observations across experiments. Culture type (e.g. WT vs KO or Vehicle vs Treated) was treated as the column factor and differentiation as the time/row factor. A mixed-effects model was selected, and a main effects model was fit (column/culture type effect, row/differentiation effect). The asterisks presented in bar graphs represent the statistical significance of the column/culture type effect. Unless otherwise notes, significant effects were determined as: p<0.05 = *, p<0.01 = **, p<0.001 = ***, p<0.0001 = ****.

### Data visualization

Schematics were generated using BioRender.com. Graphs were generated using ggplot2 in R Studio (R v4.1.1) or GraphPad Prism 10. The smaller data points on the graphs represent technical replicates (wells within a differentiation), the larger data points represent experimental/differentiation averages. Independent differentiations are distinguished by different shapes.

## RESULTS

### Robust dysregulation of proteomic signatures with ABCA7 loss-of-function in human iPSC-derived microglia

We employed CRISPR/Cas9 genetic editing to generate human induced pluripotent stem cells (iPSCs) that were deficient for ABCA7 (ABCA7-KO) by targeting exon 13 of the *ABCA7* locus in two independent genetic backgrounds (**Figure 1B, S1A-C**). In parallel, wild-type isogenic counterparts that underwent the same process were monoclonally isolated and used as the controls (ABCA7-WT) for each genetic background. Parental iPSC lines were derived from donors in the Religious Orders Study Memory and Aging Project (ROSMAP): one male (BR33) and one female (BR24) (36). Both donors were over 85 years old at the time of death, non-cognitively impaired, had no neurological or neuropathological diagnosis of AD, and were homozygous carriers of the neutral *APOE3* allele. Potential sgRNA off-target sites were identified using CRISPick. No predicted off-target sites with 1-2 nucleotide mismatches were identified, consistent with low off-target risk. Four predicted off-target sites in coding regions, each containing 3-4 mismatches, were therefore selected for PCR amplification and sequencing in all generated cell lines. No off-target editing was detected between isogenic pairs at any of these loci (**Figure S1D-G**). Since *ABCA7* has been shown to be expressed in excitatory neurons, microglia and astrocytes (27), ABCA7-WT and -KO iPSCs from both genetic backgrounds were differentiated from iPSCs into astrocytes (iAs) (41), neurons (iNs) (43) and microglia (iMGs) (78) (**Figure 1C**). ABCA7-WT and -KO iPSCs from both genetic backgrounds properly differentiated into iAs, iNs, and iMGs, as measured by immunostaining and protein quantification of cell type markers (**Figure 1D, S1H**), and that our CRISPR editing successfully eliminated full-length ABCA7 expression in all three cell-types, using a monoclonal antibody specific to the C-terminal domain (**Figure 1E-G, S1I-L**). Of note, ABCA7 was highly expressed at the protein level in iNs and iMGs, and was less abundant in iAs, consistent with single nucleus RNAseq data from human brain (27).

We next leveraged unsupervised methods to evaluate the cell-type-specific molecular changes induced by ABCA7 deficiency by performing Data-Independent Acquisition Mass Spectrometry (DIA-MS) proteomic profiling of ABCA7-WT and -KO iAs, iNs, and iMGs (**Table S1-S3**). We generated two replicates of each cell type for both genetic backgrounds (BR24/BR33) and genotypes (WT/KO) across three independent differentiations. Principal Component Analysis (PCA) of the proteomics data using PC1 and PC2 demonstrated that the samples separated by cell-type (**Figure S1M-N**). Samples also expressed proper cell-type specific protein markers (**Figure S1N**). Analysis of ABCA7 peptides confirmed complete deficiency of all detectable ABCA7 fragments (**Figure S1O-P**). We employed Linear Models for Microarray and RNA-seq Data (LIMMA) (50) analysis to identify differentially expressed proteins (DEPs) for each cell-type across both genetic backgrounds (**Figure 1H, Table S4**) and found that while iAs and iNs exhibited relatively few ABCA7-dependent changes in protein abundance (iAs = 11 DEPs, iNs = 7 DEPs), iMGs showed robust proteomic changes (DEPs = 870). This was intriguing as previous studies have focused on the function of ABCA7 in excitatory neurons, as single nuclear transcriptomic data from the human brain suggest they express the highest levels of *ABCA7* (27). These data suggest that while ABCA7 has a functional role in excitatory neurons and astrocytes, microglia rely heavily on ABCA7. We next sought to further interrogate how these molecular changes could perturb microglia on the functional level.

### Microglia alter immune response and lysosomal proteins in response to ABCA7 deficiency

To gain a better understanding of the microglial molecular pathways affected by ABCA7 deficiency, we performed Gene Set Enrichment Analysis (GSEA) on iMG proteins ranked by multiplying the degree (log_2_FC) and significance (-log(p-value)) of fold-change (**Figure 2A, Table S5**). We identified a significant upregulation of enriched pathways involved in immune response and RNA processing (transcription/splicing), and down-regulated pathways involved in the endolysosomal system and lipid metabolism.

**Figure 2:**
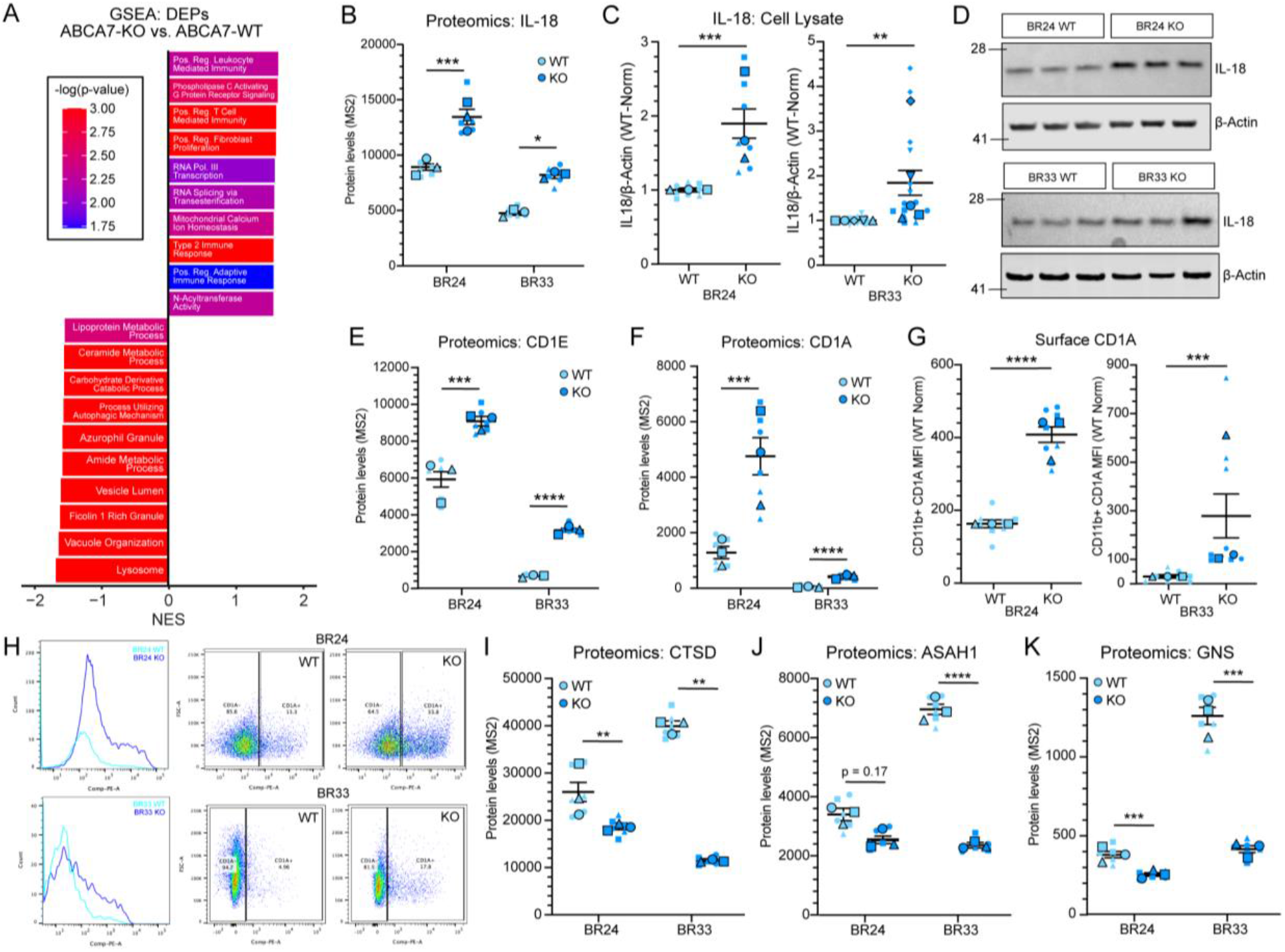
Human iPSC-derived microglia respond to ABCA7 deficiency through immunological priming and reducing proteins involved in lysosome-mediated degradation. (A) Top 10 positively and negatively significantly enriched pathways identified through preranked Gene Set Enrichment Analysis (GSEA). NES = normalized enrichment score. Full details of significantly enriched GSEA pathways, ranked gene lists, and input list of differentially expressed proteins can be found in Table S5. (B) IL-18 abundance (MS2) in BR24 and BR33 ABCA7-WT and -KO microglia from DIA-MS proteomics data. (C) Western blot analysis of IL-18 expression in BR24 (left) and BR33 (right) cell lysates. Results normalized to WT = 1 for each differentiation. (D) Example western blot of IL-18 and beta-actin in BR24 (top) and BR33 (bottom) iMG cell lysates. Left numbers denote relative molecular weights of protein standard (kDa). (E-F) CD1E (E) and CD1A (F) abundance (MS2) in BR24 and BR33 ABCA7-WT and -KO microglia from DIA-MS proteomics data. (G) Surface expression of CD1A in BR24 (left) and BR33 (right) iMGs, determined by flow cytometry. Results normalized to WT average across all differentiations within each genetic background. (H) Example flow cytometry traces of CD1A fluorescence intensity distribution (left) and dot plot (right) for BR24 (top) and BR33 (bottom) genetic backgrounds. (I-K) CTSD (I), ASAH1 (J) and GNS (K) abundance (MS2) in BR24 and BR33 ABCA7-WT and - KO microglia from DIA-MS proteomics data. For all scatter plots, large, outlined points are representative of differentiation average and small points represent well replicates within each differentiation. Different shapes denote separate differentiation. Statistical analyses display results of mixed-effects ANOVA raw data to capture within- and between-differentiation variance. Significant effect of genotype was determined: p<0.05 = *, p<0.01 = **, p<0.001 = ***, p<0.0001 = ****. Error bars = mean +/- SEM.

One of the strongest upregulated proteins found in the proteomic profiling of microglia was IL-18 (**Figure 1H, 2B**) which is a downstream effector of activation of the NLRP3 inflammasome, a disease-relevant outcome of inflammatory priming of innate immune cells. We validated the upregulation of IL-18 using western blot analysis (**Figure 2C-D; S2A**). Another set of important immune response molecules that were upregulated by ABCA7 deficiency across both genetic backgrounds are lipid antigen presentation proteins, including CD1A, and the intracellular lipid chaperone CD1E (79,80) (**Figure 2E-F**). We validated the upregulation of CD1A in both genetic backgrounds using flow cytometric analysis (**Figure 2G-H**).

The most significantly enriched down-regulated pathway in ABCA7-KO iMGs was the lysosomal protein degradation system (**Figure 2A**). These pathways encompassed a variety of down-regulated proteins measured across both genetic backgrounds including lysosomal peptidase Cathepsin D (CTSD) (**Figure 2I**), and proteins involved in lysosomal lipid metabolism such as Acid Ceramidase (ASAH1) (**Figure 2J**), and lysosomal catabolism such as N-acetylglucoasmine-6-sulfatase (GNS) (**Figure 2K**). These molecular responses of microglial ABCA7 deficiency support a crucial role for this protein in regulating a diverse array of functions, culminating with enhanced inflammatory priming and reduction of lysosomal proteolytic processing. Next, we sought to determine if these protein-level changes were consequential to microglial function, which could suggest a mechanism by which ABCA7 reduction in function causes increased AD risk.

### ABCA7 deficiency negatively impacts intracellular microglial lysosomal homeostasis

The enriched molecular pathways identified through the proteomic analyses nominated a novel role for ABCA7 in regulating the endolysosomal system. We next measured lysosomal degradation capacity using a DQ-Red Bovine Serum Albumin (DQ-BSA) assay in ABCA7-WT and -KO iMGs (81). For this assay, DQ-BSA is provided to cells, where it is taken up and processed through the endolysosomal system. DQ-BSA is quenched at a neutral pH, and it only fluoresces when degraded at the acidic pH present in the lysosomes, proving a direct measure of lysosomal cleavage activity (**Figure 3A**). In both BR24 and BR33 iMGs, lysosomal degradative capacity was reduced (**Figure 3B-C**). Since the DQ-BSA assay is reliant on endocytosis, we next tested for this process by measuring the uptake of constitutively fluorescent Alexa488-conjugated BSA using flow cytometric and live-cell imaging (**Figure S3A**). ABCA7 deficiency negatively impacted endocytosis in the BR33 but not the BR24 genetic background, suggesting that the DQ-BSA substrate was not able to get into the cell to be processed effectively in this cell line (**Figure S3B-E**). To control for this experimental caveat, we obtained an engulfment metric from the BSA488 endocytosis data by calculating the average raw (non-WT normalized) BSA488 fluorescence intensity of CD11b+ microglia across 3-4 differentiations for BR33 and BR24 ABCA7-WT and -KO iMG obtained from flow experiments (**Figure S3F**). We then divided the DQ-BSA intensity measures (**Figure 3B**) by this engulfment metric to obtain a statistic for DQ-BSA reagent that was cleaved normalized to the approximate quantity that was endocytosed. By controlling for endocytic capacity using BSA488 engulfment, we found that both BR33 and BR24 ABCA7-KO iMG had impaired DQ-BSA lysosomal processing compared to ABCA7-WT counterparts (**Figure S3G**).

**Figure 3:**
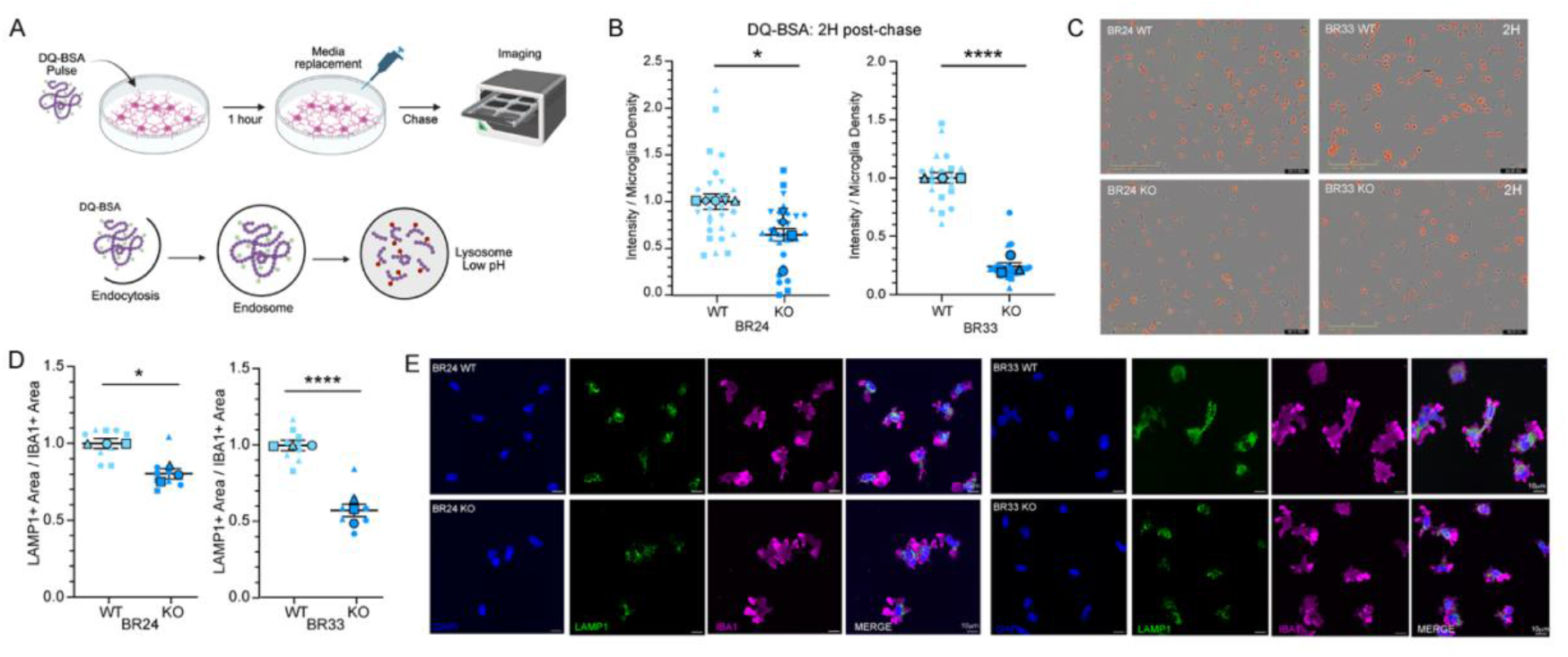
ABCA7 deficiency disrupts measures of lysosomal area and function in human iPSC-derived microglia. (A) Schematic of DQ-BSA assay. Cells area treated with DQ-BSA for 1 hour to allow for uptake. A full media change is performed to remove DQ-BSA, and images acquired two hours post-chase to measure DQ-BSA signal intensity. (B) Analysis of DQ-BSA integrated intensity normalized to microglia density for BR24 (left) and BR33 (right) ABCA7-WT and KO microglia 2-hours after DQ-BSA was removed from the media across differentiations and wells. Results are normalized to WT within each differentiation. (C) Representative images of BR24 (top) and BR33 (bottom) ABCA7-WT (left) and -KO (right) iMGs at the 2-hour post-chase time-point. Scale bar = 200µm. (D) Analysis of LAMP1+ area normalized to IBA1+ area for BR24 (left) and BR33 (right) ABCA7-WT and -KO microglia. Results were normalized to within each differentiation. (E) Representative ICC images of LAMP1 in IBA1+ microglia in BR24 (left) and BR33 (right) ABCA7-WT and -KO microglia. Scale bar = 10µm. For all scatter plots, large, outlined points are representative of differentiation average and small points represent well-level data within differentiation. Different shapes denote separate differentiation. Statistical analyses display results of mixed-effects ANOVA on raw data (not normalized to WT) to capture within- and between-differentiation variance. Significant effect of genotype was determined: p<0.05 = *, p<0.01 = **, p<0.001 = ***, p<0.0001 = ****. Error bars = mean +/- SEM.

We used additional assays to obtain measures of lysosomal content and function. Lysotracker flow cytometric analysis showed that ABCA7-KO in the BR33 genetic background negatively impacted lysosomal content (**Figure S3H-I**). Autophagic flux was also negatively impacted in BR33 ABCA7-KO iMGs, which was measured using treatments with the lysosomal V-ATPase inhibitor Bafilomycin (**Figure S3J-L**). Finally, we measured late endosome and lysosomal abundance using immunocytochemical (ICC) analysis of LAMP1, a lysosomal membrane protein, and found a significant reduction of LAMP1 area by ABCA7 deficiency in both genetic backgrounds when normalized to iMG area, providing evidence that there is overall less lysosomal content in ABCA7-KO iMG, and the lysosomes present are dysfunctional (**Figure 3D-E, S3M**). These findings support a role of ABCA7 in maintaining microglial endolysosomal homeostasis and proteolytic capacity.

### ABCA7 deficiency impairs microglial phagocytosis and alters TREM2-dependent lysosomal function

Lysosomal proteolytic processing is a key hub in the efferocytosis pathway (82), which is the mechanism by which microglia take up, digest, and excrete extracellular debris. Our current data support a role for ABCA7 in regulating lysosomal integrity, so we next investigated if this deficiency is also connected to additional aspects of efferocytosis, particularly phagocytosis. TREM2, a receptor known to regulate microglia phagocytosis, was down-regulated by ABCA7-KO in DIA-MS data, which we validated in cell lysates using western blot analysis (**Figure 4A-C, S4A**). Soluble TREM2 also was significantly reduced in the conditioned media of both BR24 and BR33 ABCA7-KO iMGs (**Figure 4D**).

**Figure 4:**
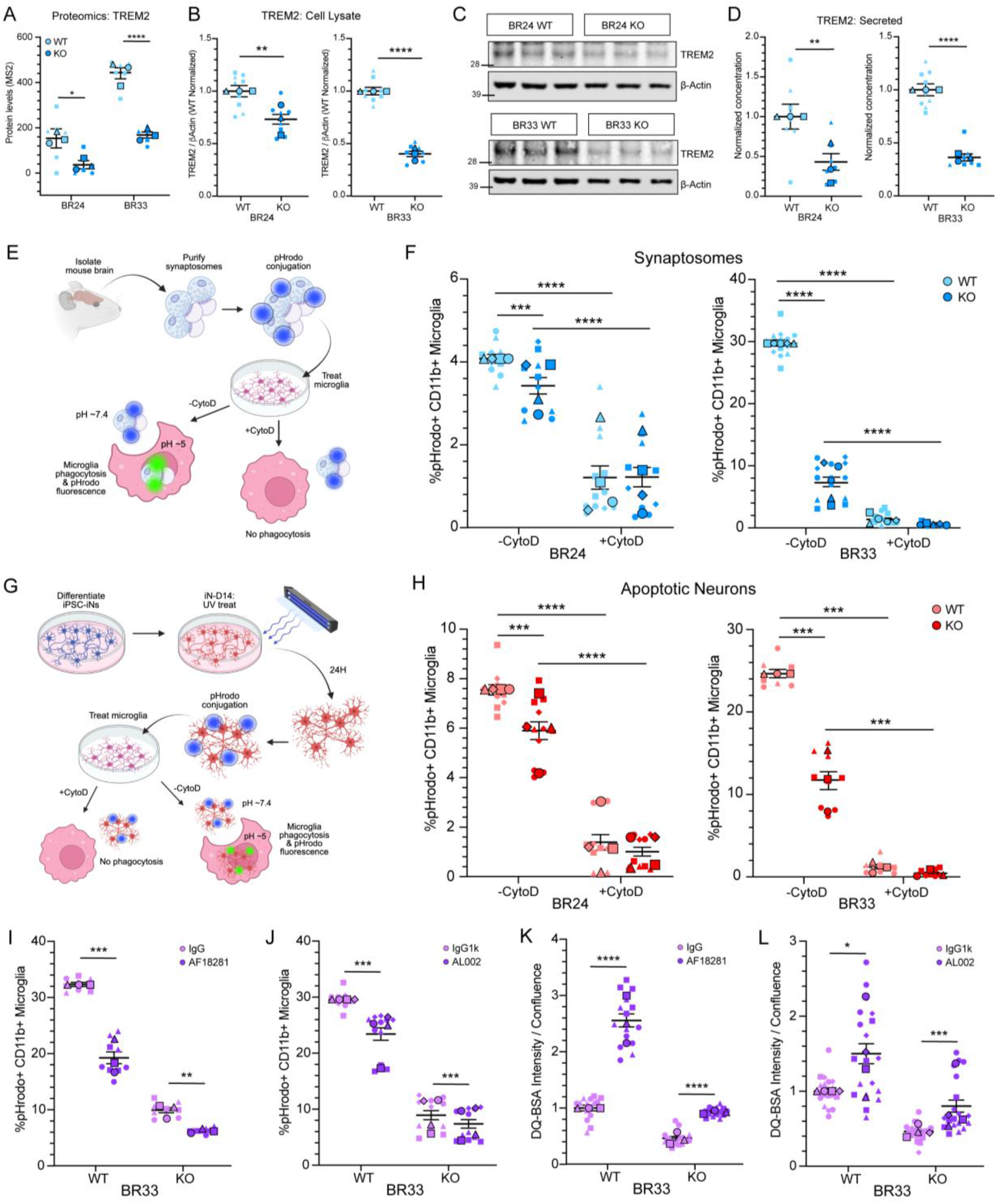
Microglia deficient in ABCA7 show reduced TREM2 protein levels and impaired phagocytosis. (A) TREM2 abundance (MS2) in BR24 and BR33 ABCA7-WT and -KO iMGs from DIA-MS proteomics data across wells and differentiations. (B) Western blot analysis of TREM expression in BR24 (left) and BR33 (right) cell lysates. Results normalized to WT for each differentiation. (C) Example western blot of TREM2 and β-actin in BR24 (top) and BR33 (bottom) iMGs cell lysates. Numbers on the left denote relative molecular weights of protein standard (kDa). (D) Secreted TREM2 protein levels measured in conditioned media of BR24 (left) and BR33 (right) iMG cultures. Results were normalized to β-actin expression measured by western blot of cell lysates, and to WT within each differentiation. (E) Schematic of preparation and pHrodo conjugation of mouse synaptosomes. pHrodo-labeled synaptosomes were provided to either untreated or CytoD pre-treated microglia for 12-18 hours followed by flow cytometric analysis. (F) Flow cytometric analysis of iMG synaptosome engulfment. Percent (%) CD11b+ pHrodo+ microglia analyzed in BR24 (left) and BR33 (right) ABCA7-WT and -KO microglia untreated (CytoD) or pre-treated (+CytoD) with cytochalasin D. Results from each differentiation were normalized to the WT average across all differentiations. (G) Schematic of preparation and pHrodo conjugation of human iPSC-derived apoptotic neurons (ANs). NGN2 iPSC neurons (iNs) were differentiated for 14 days (iN-D14), then irradiated with 50,000 joules of ultraviolet (UV) light. 24 hours post-irradiation, neurons were dissociated and conjugated with pHrodo, then provided to either CytoD pre-treated or untreated microglia for 12-18 hours. (H) Flow cytometric analysis of microglial AN engulfment. Percent (%) CD11b+ pHrodo+ microglia in BR24 (left) and BR33 (right) ABCA7-WT and -KO microglia untreated (-CytoD) or pre-treated (+CytoD) with cytochalasin D. Results from each differentiation were normalized the WT CytoD-average across all differentiations. (I-J) Flow cytometric analysis of microglia AN engulfment after TREM2 antibody treatment. Percent (%) CD11b+ pHrodo+ microglia in BR33 ABCA7-WT and -KO microglia treated with AF18281 (I) or AL002 (J) anti-TREM2 antibody. Results from each differentiation were normalized to the WT vehicle average across all differentiations for visualization. (K-L) Analysis of DQ-BSA integrated intensity normalized to microglia density for BR33 ABCA7-WT and -KO microglia treated with AF18281 (K) or AL002 (L) anti-TREM2 antibodies 2-hours after DQ-BSA was removed from the media across differentiations and wells. Results are normalized to WT vehicle condition within each differentiation for visualization. For all scatter plots, large, outlined points are representative of differentiation average and small points represent well-level data within each differentiation. Different shapes denote separate differentiation. Statistical analyses display results of mixed-effects ANOVA on raw non-WT-normalized data to capture within- and between-differentiation variance. Significant effect of genotype was determined: p<0.05 = *, p<0.01 = **, p<0.001 = ***, p<0.0001 = ****. Error bars = mean +/- SEM.

We next tested if ABCA7 deficiency impacted functional measures of cellular uptake using live cell flow cytometric based approaches. Since phagocytosis is reliant on receptor recognition of substrates and occurs in response to a variety of substrates, we leveraged multiple substrate exposures to test if this process had a more specific or general impairment in our ABCA7-KO iMGs. First, we tested the engulfment of pHrodo-conjugated adult mouse brain-derived synaptosomes (**Figure S4B**). The pHrodo dye is not fluorescently active until it is exposed to an acidic pH that is present intracellularly, allowing for the measurement of uptake of a specified substrate. We also pre-treated a subset of cells with CytoD as a negative control to ensure we were measuring engulfed fluorescent substrate (**Figure 4E**). Both BR33 and BR24 ABCA7-KO iMGs exhibited significantly reduced phagocytosis of synaptosomes (**Figure 4F, S4C**). Next, we tested phagocytosis of human iPSC-derived apoptotic neurons (ANs) (**Figure 4G**). Complementing the synaptosome results, we found that ABCA7-KO iMGs in both BR33 and BR24 genetic backgrounds exhibited reduced phagocytosis of this human-derived substrate (**Figure 4H, S4D**). Finally, we tested if ABCA7 deficiency affected the phagocytosis of smaller substrates with higher degrees of lipidation, and exposed microglia to crude and purified myelin from mouse brain (**Figure S4E**). ABCA7-KO iMGs exhibited reduced phagocytic capacity to these lipidated substrates in both genetic backgrounds (**Figure S4F-I**).

To interrogate whether the observed down-regulation of TREM2 contributes to lysosomal and phagocytic abnormalities observed in ABCA7-KO iMG, we perturbed TREM2 signaling using two different anti-TREM2 antibodies: a goat anti-human TREM2 (AF18281) and a humanized monoclonal anti-TREM2 (AL002) that was recently tested in human AD clinical trials (showing non-significant results) (83). While AF18281 and AL002 discordantly reduced and increased surface TREM2 levels, respectively, both antibodies reduced phagocytosis of human ANs in ABCA7-WT and -KO iMG (**Figure 4I-J, Figure S4J-K**). Interestingly, while reducing phagocytic capacity, both TREM2 antibodies significantly increased DQ-BSA lysosomal processing in both ABCA7-WT and -KO iMG (**Figure 4K-L, S4L-M**), forming a functional connection between TREM2 signaling, phagocytosis, and lysosomal activity. However, while ABCA7-KO reduces both phagocytosis and lysosomal homeostasis, these TREM2 perturbations rescued the lysosomal phenotype while exacerbating the phagocytosis phenotype. Therefore, negative impacts on phagocytosis may serve as an adaptive feedback mechanism in response to microglial ABCA7 deficiency that is separable from TREM2 signaling.

### ABCA7 deficiency disrupts intracellular microglial lipid balance

Previous studies have established ABCA7’s role in regulating lipid homeostasis in mice, human neurons, and cerebral organoids (24,27,28,30,84). Here, we aimed to complement this work to interrogate if loss of ABCA7 has similar impacts on our human microglial model. Therefore, we next interrogated if and how ABCA7 disruption affects intracellular lipid properties in human iPSC-derived microglia using comprehensive targeted lipidomics (18,85) on three differentiations of ABCA7-WT and -KO iMGs (**Table S6**). Using PCA of the complete lipidomic dataset, a striking separation of ABCA7-WT and -KO iMGs was observed in principal component 1 (PC1), supporting an impactful role for ABCA7 in regulating microglial lipid homeostasis (**Figure 5A**). To identify individual lipids contributing to this genotype-dependent lipid divergence, a lipidome-wide comparison was performed to identify differentially abundant lipids in ABCA7-KO versus ABCA7-WT iMGs (**Figure 5B**). Overall, we observed a robust upregulation of triacylglycerols (TG), and a down-regulation of several phosphatidylcholines (PC) and phosphatidylinositol (PI) species consistent with broad lipid remodeling characterized by increased neutral lipids and reduced abundance of several membrane phospholipid species. Lipid Set Enrichment Analysis (59) (LSEA) identified a dysregulation of several overall lipid classes, including negative enrichment of hexosylceramide (HexCer), lysophosphatidylcholine (LPC), ceramide (Cer), sphingomyelin (SM), bis(monoacylglycerol)phosphate (BMP), and phosphatidylethanolamine (PE) classes, and positive enrichment of phosphatidylethanolamine plasmalogens (PE P-) and TG classes (**Figure 5C**). Many down-regulated lipids have been reported to be localized to the Golgi, lysosome, and plasma membrane cellular compartments (**Figure 5D**), consistent with our proteomic and functional evidence of altered lysosomal biology following ABCA7 loss (**Figures 2, 3**). We also used these data to explore if and how differentially abundant lipids could affect membrane structural characteristics. Differentially abundant lipids in ABCA7-KO iMGs were enriched towards consequences predicted to increase bilayer thickness, negative intrinsic curvature, reduced lateral diffusion, and elevation transition temperature (**Figure S5A-E**).

**Figure 5:**
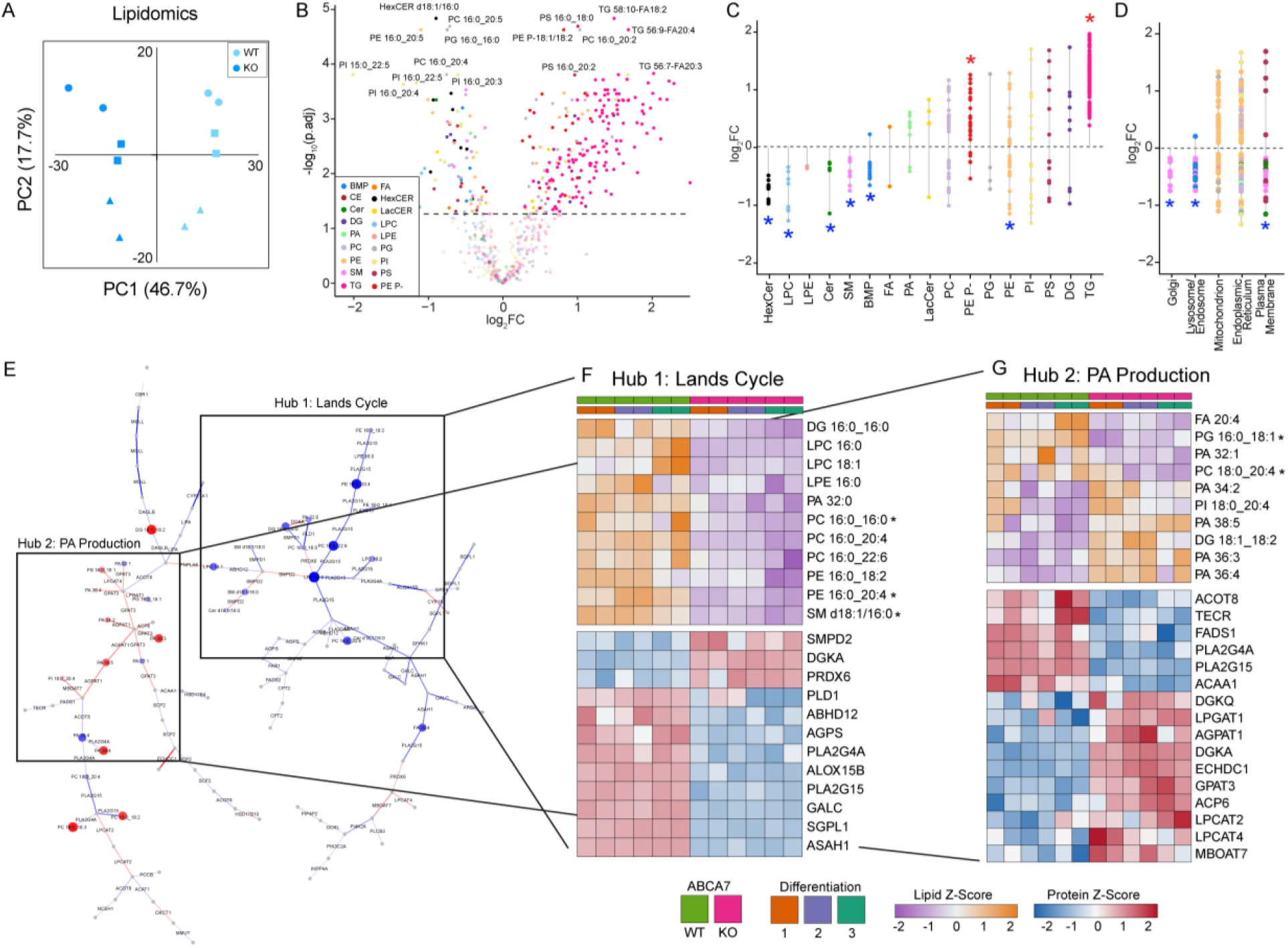
Microglial ABCA7 deficiency shapes intracellular metabolic pathways suggestive of cumulative oxidative stress. (A) Lipidomic profiling performed on BR33 ABCA7-WT and -KO iMGs across three differentiations, with two wells per differentiation. Analysis of the first two principal components (PC1, PC2) of the lipidomic profiles from BR33 ABCA7-WT and -KO iMGs are shown. Percentages reflect the amount of variance explained by each principal component. (B) Volcano plot depicting differential lipid abundance of ABCA7-WT versus -KO microglia. Horizontal dashed bar denotes significance threshold cutoff of adjusted p-value = 0.05. Dots are colored by their primary lipid class. BMP = Bis(monoacylglycerol)phosphate; CE = Cholesterol Ester; Cer = Ceramide; DG = Diacylglycerol; PA = Phosphatidic Acid; PC = Phosphatidylcholine; PE = Phosphatidylethanolamine; PE P-= Phosphatidylethanolamine plasmalogen; SM = Sphingomyelin; TG = Triglyceride; FA = Fatty Acid; HexCER = Hexosylceramide; LacCER = Lactosylceramide; LPC = Lysophosphatidylcholine; LPE = Lysophosphatidylethanolamine; PG = Phosphatidylglycerol; PI = Phosphatidylinositol; PS = Phosphatidylserine. (C) Lollipop plot of significantly differentially expressed lipids separated by class. log_2_FC = degree of abundance difference between WT and KO iMGs for each lipid. Asterisks denote lipid classes that are significantly positively (red asterisk) and negatively (blue asterisk) enriched, calculated using Lipid Set Enrichment Analysis (LSEA). (D) Lollipop plot of significantly differentially expressed lipids that could be assigned to a cellular compartment. Points are colored by lipid class assigned in 2B. Asterisks denote cellular compartments significantly negatively (blue asterisk) enriched in lipidomics data identified via LSEA. (E) Top interconnected lipid-protein network identified through GATOM network analysis. Lipid nodes are colored and sized by most negative (blue) to positive (red) log_2_ fold change between ABCA7-WT and -KO iMGs. Grey nodes represent metabolite intermediaries not measured in the current dataset. Edges represent proteins involved in metabolite or lipid conversion and colored by log_2_ fold change between ABCA7-WT and -KO iMGs. Boxed areas are two central, manually selected hubs of dysregulated protein-lipid interactions identified in the network. (F-G) Heatmaps of lipid and protein members of identified Lands cycle (F) and phosphatidic acid production (G) central network hubs. Lipid and protein abundance levels were z-transformed, with lipid relative abundance values depicted in orange/purple, and protein relative abundance in red/blue heatmaps. Asterisks denote lipid was found to be altered by ABCA7 deficiency in orthogonal study (28). In all analysis graphs, large, outlined points are representative of differentiation average and small points represent well-level replicates within differentiation. Different shapes denote separate differentiations. Statistical analyses display results of mixed-effects ANOVA on raw data (not normalized to WT) to capture within- and between-differentiation variance. Significant effect of genotype was determined: p<0.05 = *, p<0.01 = **, p<0.001 = ***, p<0.0001 = ****. Error bars = mean +/- SEM.

### Multi-omic integration identifies coordinated alterations in phospholipid removdeling pathways in ABCA7-deficient microglia

To identify coordinated lipid-protein pathways altered by ABCA7 deficiency, we performed a **G**ene **a**nd Me**t**ab**o**lite **M**etabolic (GATOM) (70) network analysis, integrating our unbiased lipidomics and proteomics data generated from BR33 ABCA7-WT and -KO microglia. Using a generalized maximum-weight connected subgraph approach, we identified a metabolic sub-network enriched for differentially abundant proteins and lipids in ABCA7-KO iMGs. Evaluation of this network highlighted two interconnected regions involving phospholipid remodeling and phosphatidic acid (PA)-associated metabolism (**Figure 5E, S5F**). Several lipid species involved in both of these candidate network hubs were dysregulated by ABCA7 deficiency in recent work using human cortical organoids (28) (**denoted by asterisk in Figure 5E**), suggesting aspects of ABCA7-associated lipid dysregulation are shared between neural cell types and microglia.

One prominent network region contained components of the Lands cycle, which participates in phospholipid remodeling (**Figure 5F**). The Lands cycle is involved in the intracellular processing of phospholipids, primarily phosphatidylcholine (PC) (86). Intriguingly, PC biosynthesis and transport is a known function of ABCA7 (23,84,87). Dysregulation of aspects of PC processing such as the Kennedy pathway, which provides phospholipid precursors that are remodeled in the Lands cycle (86), were reported to be affected with ABCA7 loss-of-function mutations in human neurons (27). Here, in human iMGs we observed reduced LPC and PE species, altered abundance of phospholipase proteins involved in phospholipid remodeling, and increased abundance of lysophosphatidylcholine acyltransferase (LPCAT) enzymes that catalyze the re-acylation of lysophospholipids (**Figure 5F**). PLA2G15, a lysosomal hydrolase involved in lipid catabolism, including BMP processing (88), was also reduced in ABCA7-KO iMGs. Consistent with this observation, BMP was a negatively enriched class in the lipidomic dataset (**Figure 5C**). Together, these coordinated protein and lipid changes identify phospholipid remodeling as a metabolic pathway altered by ABCA7 deficiency in microglia, although these steady-state measurements do not establish the direction or magnitude of flux through individual reactions.

A second region of the integrated network contained PA species and proteins involved in PA-associated metabolism (**Figure 5G**). Several PA species were increased in ABCA7-KO iMGs, together with increased abundance of enzymes including AGPAT1, GPAT3, DGKA, and DGKQ (**Figure 5G**). These findings further support broad remodeling of phospholipid metabolism following loss of ABCA7 in microglia, but do not by themselves distinguish changes in synthesis, degradation, retention or transport.

Increased intracellular PA can be deleterious to the cell as it has been reported to be vulnerable to lipid peroxidation which triggers downstream inflammatory and oxidative stress pathways (89,90). Since lipid peroxidation has been reported previously in ABCA7-deficient human neurons (24), we next asked whether a similar phenotype was present in ABCA7-deficient microglia. Our integrated analyses also identified alterations in proteins and lipids associated with peroxisomal β-oxidation and arachidonic acid metabolism (**Figure S5G-H**), pathways relevant to cellular lipid peroxidation. We therefore directly measured lipid peroxidation using the BODIPY-C11 sensor. ABCA7-KO iMGs exhibited a significantly increased oxidized-to-reduced BODIPY-C11 signal compared to their isogenic counterparts (**Figure S6I-J**), demonstrating increased intracellular lipid peroxidation. Together, these data identify coordinated alterations in phospholipid remodeling and increased lipid peroxidation as features of ABCA7 deficiency in human microglia, paralleling aspects of lipid dysregulation previously reported in ABCA7-deficient neuronal models (24,27,28).

### Lysosomal dysfunction and lipid droplet accumulation are alleviated by inhibition of triacylglycerol synthesis in ABCA7-deficient microglia

While the disruption of dynamics of lipid-protein metabolic networks were consistent with aspects of lipid dysregulation previously reported in ABCA7-deficient human neurons (24,27,28), ABCA7-deficient microglia exhibited a particularly striking increase in both individual triacylglycerol (TG) species (**Figure 5B**) and the TG lipid class (**Figure 5C**). We first asked whether altered fatty-acid uptake or efflux could contribute to this phenotype. ABCA7-KO iMGs exhibited reduced uptake of labeled fatty acids compared to their isogenic WT counterparts (**Figure 6A-B**, **S6A-C**). Conversely, there was no evidence of differences in fatty acid efflux between ABCA7-WT and - KO iMGs (**Figure 6C**, **S6D**). Therefore, TG accumulation was not explained by increased uptake or reduced efflux of exogenous fatty acids, suggesting that altered intracellular lipid metabolism and/or storage contributes to this phenotype.

**Figure 6:**
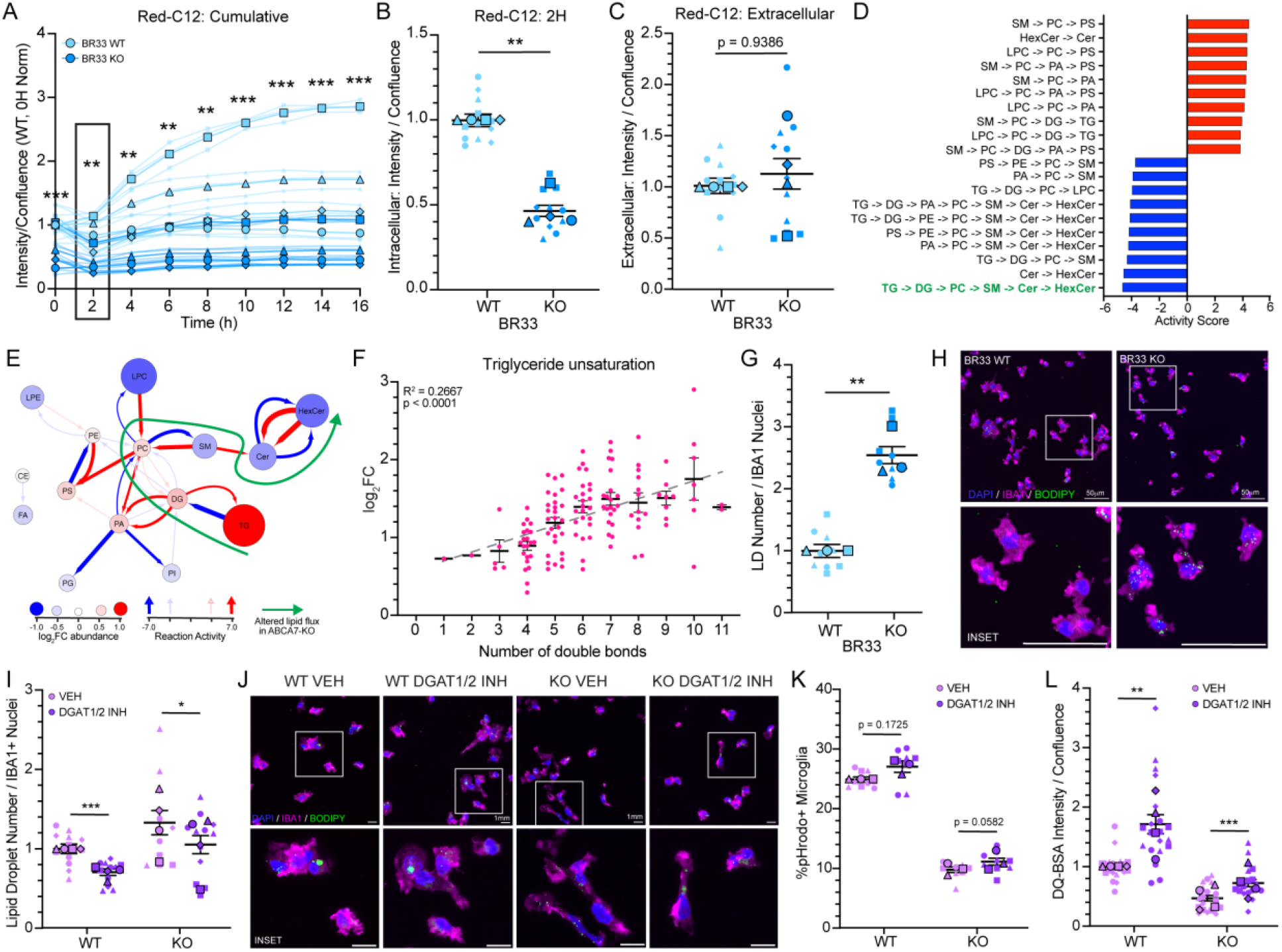
Inhibiting TG synthesis in ABCA7-deficient microglia rescues lipid droplet accumulation and lysosomal impairments. (A) Longitudinal measurement of intracellular signal of fatty acids labeled for 16 hours with BODIPY Red-C12 in BR33 ABCA7-WT and -KO iMGs. Solid, thick lines correspond to differentiation averages, with thin, faint lines corresponding to individual replicates within each experiment. Results were normalized to WT = 1 for 0H time-point for visualization. (B) Analysis of intracellular fatty acids 2 hours post-treatment with Red-C12 lipid sensor in BR33 ABCA7-WT and -KO iMGs. Results normalized to WT = 1 for each differentiation for visualization. (C) Analysis of extracellular Red-C12 fatty acids 4 hours post-media replacement in BR33 ABCA7-WT and -KO iMGs. Results normalized to WT = 1 for each differentiation for visualization. Large, outlined points are representative of differentiation average and small points represent technical replicates within differentiation. Different shapes denote separate differentiation. (D) Top ten positively (red) and negatively (blue) enriched lipid turnover pathways identified using BioPAN analysis through LipidMAPS. The pathway highlighted in green shows the strongest significant negative enrichment and encompasses the largest number of lipid class transitions. (E) Network integration of significantly enriched pathways identified in the BioPAN analysis through LipidMAPS. Node colors and sizes are based on degree of abundance of each lipid class. Edge colors and widths are based on degree and direction of predicted reaction activity. Green arrow is summarized prediction of the lipid flux pathway that is predicted to be the most altered by ABCA7 deficiency. (F) Scatter plot of TGs classified by degree of unsaturation (number of double bonds) and abundance difference between ABCA7-WT and -KO iMGs. Horizontal line and error bars are mean log_2_FC and SD for lipids with each level of unsaturation. Grey line denotes results from linear regression of double bond number to mean log_2_FC of TG abundance. R^2^ and p-value of linear regression are denoted in upper left-hand corner of graph. (G) Quantification of BODIPY+ lipid droplet number normalized to IBA1+ nuclei in BR33 ABCA7-WT and -KO microglia. Results were normalized to WT = 1 within each differentiation for visualization. (H) Representative ICC images of DAPI (blue), BODIPY (green) and IBA1 (magenta) in BR33 ABCA7-WT and -KO iMGs. Scale bar = 50µm. (I) Quantification of BODIPY+ lipid droplet number normalized to IBA1+ nuclei in BR33 ABCA7-WT and -KO microglia treated with DGAT1/2 inhibitors (DGAT1/2 INH) or DMSO vehicle (VEH). Results were normalized to WT VEH = 1 within each differentiation for visualization. (J) Representative ICC images of DAPI (blue), BODIPY (green) and IBA1 (magenta) in BR33 ABCA7-WT and -KO iMGs treated with DGAT1/2 inhibitors (DGAT1/2 INH) or DMSO vehicle (VEH). Scale bar = 1mm. (K) Percent (%) CD11b+ pHrodo+ microglia in BR33 ABCA7-WT and -KO microglia treated with DGAT1/2 inhibitor (DGAT1/2 INH) or DMSO vehicle (VEH). Results from each differentiation were normalized to the WT VEH average across all differentiations for visualization. (L) Analysis of DQ-BSA integrated intensity normalized to microglia density for BR33 ABCA7-WT and -KO microglia treated with DGAT1/2 inhibitor (DGAT1/2 INH) 2-hours after DQ-BSA was removed from the media across differentiations and wells. Results are normalized to WT VEH condition within each differentiation. In all analysis graphs, large, outlined points are representative of differentiation average and small points represent well-level replicates within differentiation. Different shapes denote separate differentiations. Statistical analyses display results of mixed-effects ANOVA on raw data (not normalized to WT) to capture within- and between-differentiation variance. Significant effect of genotype was determined: p<0.05 = *, p<0.01 = **, p<0.001 = ***, p<0.0001 = ****. Error bars = mean +/- SEM.

To further investigate intracellular lipid pathways associated with TG accumulation, we used BioPAN to integrate the lipidomic data with known lipid-conversion reactions curated in the LipidMAPs database (66,67). Pathway activity scores were calculated from differences in lipid abundance between ABCA7-KO and -WT iMGs (**Figure 6D**) and visualized as a network in which edges represent predicted lipid-conversion activities and nodes represent lipid classes (**Figure 6E**). This analysis identified TG-associated lipolytic pathways among those with the most negative predicted activity scores, consistent with reduced utilization of stored TGs (**Figure 6E**). The network also highlighted coordinated reductions in several sphingolipid classes, including sphingomyelin (SM), ceramide (Cer), and hexosylceramide (HexCer). Because BioPAN infers pathway activity from steady-state lipid abundance, these analyses nominate altered TG utilization and related lipid-conversion pathways but do not directly measure metabolic flux.

Closer inspection of the lipidomics dataset showed that TGs with higher amounts of unsaturation were more abundant in ABCA7-KO iMGs (**Figure 6F**). This pattern appeared specific to TGs, as phospholipid species did not exhibit a comparable shift in unsaturation (**Figure S6E**). Because TGs are major constituents of lipid droplets (LDs) (91), and accumulation of polyunsaturated TGs has previously been associated with LD accumulation in *APOE4/4* astrocytes (18,92), we directly measured LD abundance. ABCA7-KO iMGs from both BR24 and BR33 backgrounds exhibited significantly increased LD accumulation relative to their isogenic WT counterparts (**Figure 6G-H, S6F-G**).

We next asked whether reducing TG synthesis and LD accumulation could modify the functional phenotypes associated with ABCA7 deficiency. Treatment with inhibitors of diacylglycerol acid transferases 1 and 2 (DGAT1/2), which catalyze the final step in TG synthesis, reduced LD accumulation in ABCA7-KO iMGs and increased lysosomal degradative capacity, without significantly altering phagocytosis or TREM2 surface abundance (**Figure 6I-L, S6I-J**). These findings provide functional evidence linking abnormal TG metabolism and LD accumulation to impaired lysosomal processing in ABCA7-deficient microglia. Notably, restoration of lysosomal activity occurred without a corresponding change in phagocytic capacity, indicating that these two phenotypes can be functionally dissociated.

### A common AD risk variant in ABCA7 alters microglial lysosomal and inflammatory function

Our current work, leveraging loss-of-function cellular models was helpful for dissecting out the molecular and metabolic pathways controlled by ABCA7. However, while AD risk variants in *ABCA7* are associated with loss of protein function, they seldom cause complete protein deficiency. A common *ABCA7* variant associated with increased AD risk is the rs3752246 single nucleotide polymorphism (SNP), which causes an alanine to glycine amino acid change at position 1527 in the protein coding sequence (5). While the functional consequence of this variant is not completely known, recent work using computational modeling of protein folding has hypothesized it may affect the flexibility of the extracellular domain, impacting the ability for the channel to transport lipids through the plasma membrane (27). We leveraged an existing metabolomic dataset of 6,136 human plasma samples that are linked to genome-wide genetic data in a Finnish cohort (76) to interrogate whether any of the identified metabolites are associated with rs3752246 in the plasma. The abundance of two metabolites, lactosyl-N-nervonoyl-sphingosine (d18:1/24:1) and lactosyl-N-palmitoyl-sphingosine (d18:1/16:0) were significantly associated with rs3752246. We next performed GWAS analyses to interrogate which SNPs have the strongest association with levels of these two metabolites and found that SNPs at the ABCA7 locus have the strongest association with these metabolites relative to all other SNPs analyzed genome-wide (**Figure 7A-B**). In this cohort, the risk allele of rs3752246 was associated with accumulation of these metabolites, both of which are species of lactosylceramide (LacCer). These findings from human plasma are concordant with the lipid and proteomic profiles observed with loss-of-function of ABCA7 in iMGs, which also showed evidence for a reduction of enzymes involved in the LacCer to Cer conversion such as GBA, GBA2, GBA3 and GLB1 (**Figure 7C**). These data provided preliminary evidence that the rs3752246 common AD risk variant has similar functional consequences to our *in vitro* ABCA7 deficiency model; we thus next aimed to functionally interrogate this using additional approaches.

**Figure 7:**
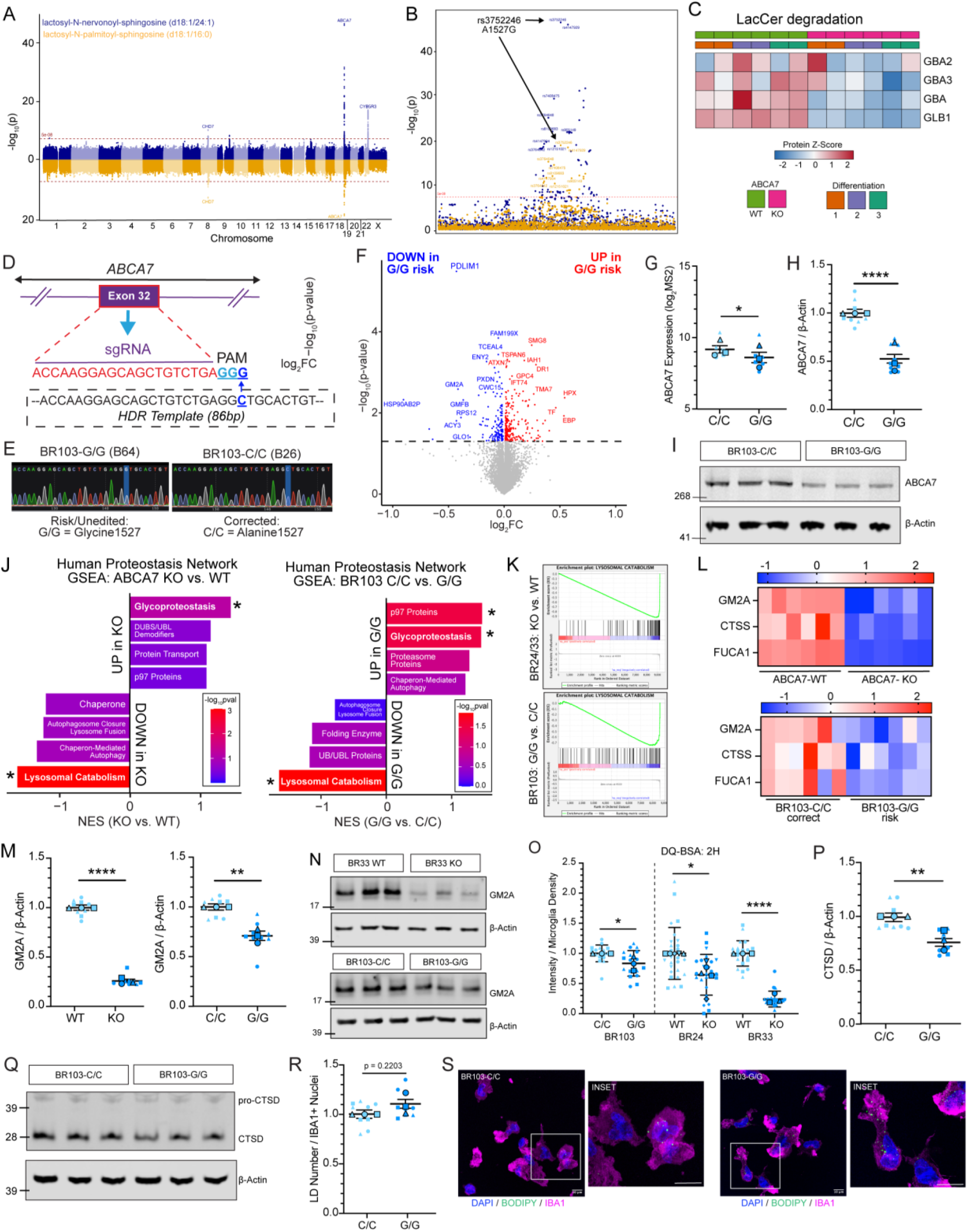
Correction of a common Alzheimer’s risk variant in ABCA7 improves measures of microglial lysosomal function. (A) Manhattan plot demonstrating genome-wide association of genetic variation in *ABCA7* with two LacCer species identified in human plasma. Genetic and metabolomic data were obtained from Yin et al, 2022 (76) and re-analyzed here to identify *ABCA7* variant-associated phenotypes. (B) Detailed Manhattan plot of the *ABCA7* locus demonstrating the G1527A common variant (rs3752246) is the most significantly associated with accumulated plasma LacCer-associated metabolites. (C) Heatmap of differentially expressed proteins associated with LacCer degradation in ABCA7-WT and -KO iMGs. Lipid and protein abundance levels were z-transformed, with protein relative abundance in red/blue heatmaps. (D) Schematic of CRISPR/Cas9 correction of rs3752246 in BR103 iPSCs. The sgRNA sequence is highlighted in red, with the adjacent PAM sequence in blue. Underneath in the dashed box is the template provided to cells to perform HDR. The underlined dark blue letter in the PAM site and HDR template is the base pair being altered corresponding to rs3752246. (E) Sanger sequencing traces of BR103 rs3752246-HDR iPSC clones, with the rs3752246 site highlighted in light blue. The left clone (BR103-B64) is an un-edited parental line, and the clone on the right (BR103-B26) has been edited to correct rs3752246 to the major allele C/C genotype. (F) Volcano plot depicting differences in proteomic abundance between BR103-G/G to corrected BR103-C/C isogenic counterparts. Proteins were analyzed by their log_2_fold-change expression between G/G and C/C iMGs (x-axis) and statistical significance of change (-log_10_ p-value) determined using LIMMA analysis. Horizontal dashed line denotes log_10_ p-value < 0.05 significance threshold. Blue and red points are proteins significantly down-regulated and upregulated in BR103-G/G compared to BR103-C/C samples, respectively. (G) ABCA7 abundance (log_2_MS2) in BR103 microglia from DIA-MS proteomics data. (H) Western blot analysis of ABCA7 expression in BR103 cell lysates. Results normalized to BR103-C/C for each differentiation for visualization. (I) Example western blot of ABCA7 and β-actin in BR103 iMG cell lysates. Left numbers denote relative molecular weights of protein standard (kDa). (J) Top four enriched pathways from GSEA preranked analysis using Human Proteostasis Network gene sets in ABCA7-WT versus -KO iMGs (left) and BR103 BR103-C/C versus BR103-G/G iMGs (right). Asterisks denote gene set with significant enrichment in the dataset. Bolded terms denote gene sets with shared enrichment between both comparisons. (K) Enrichment plot from GSEA of the “Lysosomal Catabolism” gene set from the Human Proteostasis Network showing results from preranked analysis of differentially expressed proteins between ABCA7-WT versus -KO iMGs (top) and BR103-C/C versus BR103-G/G (bottom) iMGs. (L) Heatmap of scaled protein abundance values of shared leading-edge proteins from the “Lysosomal Catabolism” GSEA results from BR33 ABCA7-WT versus -KO (left) and BR103-C/C versus BR103-G/G (right) iMGs. (M) Western blot analysis of GM2A and β-actin in BR33 (left) and BR103 (right) iMGs. Results were normalized to WT = 1 or BR103-C/C = 1 within each differentiation. (N) Example western blot of GM2A and β-actin in BR33 (top) and BR103 (bottom) iMGs. Numbers on the left denote relative molecular weights of protein standard (kDa). (O) Analysis of DQ-BSA integrated intensity (2-hours after DQ-BSA was removed from the media) normalized to microglia density for BR103 iMGs on same plot as BR24 and BR33 ABCA7-WT and -KO iMGs (data from Figure 3B) for efficient comparison. Results were normalized to C/C = 1 or WT = 1 within each differentiation for visualization. (P) Western blot analysis of mature CTSD and beta-actin in BR103 iMGs. Results were normalized to BR103-C/C for each differentiation. (Q) Example western blot of CTSD and β-actin in BR103 iMGs. Left numbers denote relative molecular weights of protein standard (kDa). (R) Quantification of BODIPY+ lipid droplet number normalized to IBA1+ nuclei in BR103 iMGs. Results were normalized to BR103-C/C within each differentiation. (S) Representative ICC images of BODIPY and IBA1 in BR103 iMGs. Scale bar = 10µm. For all scatter plots, large, outlined points are representative of differentiation average and small points represent well-level replicates within differentiation. Different shapes denote separate differentiation. Statistical analyses display results of mixed-effects ANOVA on raw data to capture within- and between-differentiation variance. Significant effect of genotype was determined: p<0.05 = *, p<0.01 = **, p<0.001 = ***, p<0.0001 = ****. Error bars = mean +/- SEM.

We had previously generated iPSCs from a ROSMAP participant (BR103) that was homozygous for the rs3752246 AD risk variant in *ABCA7*, and who had both a pathological and clinical diagnosis of AD at the time of death. Since this participant was also homozygous for the neutral *APOE3* allele, this provided an opportune APOE4-independent cellular model to understand the mechanistic consequences of the rs3752246 *ABCA7* AD risk variant in human iPSC-derived microglia. We performed CRISPR-Cas9 genetic editing on iPSCs from this donor using a homologous DNA template to generate iPSCs that were corrected for the rs3752246 major allele (BR103-C/C) that corresponded to their isogenic minor allele risk genotype (BR103-G/G) counterparts (**Figure 7D-E**). We employed unbiased DIA-MS proteomics on differentiated microglia from BR103-C/C corrected and BR103-G/G isogenic clones that underwent the same targeting procedures (**Table S7**). It is expected that a single SNP with an odds ratio (OR) of 1.3 (93) would have much less dramatic effects on the proteome compared to complete deletion of the associated gene. Thus, a relaxed p-value cutoff (p<0.05) was used to identify 349 proteins that were differentially expressed between BR103-C/C corrected and BR103-G/G isogenic iMGs (**Figure 7F**). 33 of the DEPs in the BR103 clones were identified to be significantly (adj.p<0.05) differentially expressed between BR24/BR33 ABCA7-WT and -KO iMGs (**Table S8**). Several immunoregulatory proteins were downregulated by ABCA7 deficiency and altered in expression with correction of the rs3752246 variant (**Figure S7A**, **Table S8**), including PDLIM1 which has been implicated in NF-kB signaling (94) and CLEC10A which is downstream of the TLR7/8 signaling pathway (95). These data support the hypothesis that a common AD risk variant in *ABCA7* disrupts immune-related processes similar to those observed in ABCA7-deficient human iPSC-derived microglia, and that correction of the variant reverses this effect.

Another DEP identified in the BR103 proteomics data was ABCA7 itself, although the magnitude of the change was modest (**Figure 7G**). This was intriguing because the rs3752246 variant is not hypothesized to directly regulate ABCA7 protein expression, although its effects in microglia remain poorly characterized. We therefore examined ABCA7 protein levels using western blot analysis in independent differentiations. These analyses revealed a significant reduction in ABCA7 abundance in iMGs with rs3752246 risk (G/G) alleles (**Figure 7H-I; S7B**). Because the magnitude of this effect was greater by western blot than by proteomics, we next examined the individual ABCA7 peptides detected by mass spectrometry to determine whether regional differences in protein abundance might account for this discrepancy (**Figure S7C-D, Table S7**). The monoclonal antibody used for western blotting recognizes the C-terminal region of ABCA7. Among the detected peptides, a single peptide (P5) mapping to the first nucleotide binding domain (ATP1) that was significantly downregulated in the BR103-G/G clones compared to their corrected isogenic counterparts (**Figure S7D**), with a trend towards a reduction in other peptides. However, the degree of change of this single peptide didn’t correspond to the robust increase observed in the western blot, likely reflecting the known issue with ratio compression inherent to isobaric labeling-based mass spectrometry (96).

We next sought to investigate proteomic molecular signatures that were conserved by ABCA7 deficiency and the rs3752246 ABCA7 risk variant. Since ABCA7 deficiency strongly impacted lysosomal protein networks in iMGs, we interrogated changes in BR103 and ABCA7-KO DEPs by performing GSEA on curated gene sets of human proteostasis networks (51,52) (**Figure S7E, Table S9**). Consistent with ABCA7-KO findings (**Figure 2**), “lysosomal catabolism” was the top negatively enriched pathway, and “glycoproteostasis” was positively enriched with rs3752246 G/G iMGs compared to C/C corrected isogenic iMGs, concordant with the molecular consequences of complete ABCA7 deficiency (**Figure 7J**). Further, the specific proteins that were concordantly altered between the KO and variant models were categorized into the top enriched pathway for both proteomics datasets, “lysosomal catabolism” (**Figure 7K-L**). Using western blot analysis, we validated with an independent method that one of these proteins, GM2A, was significantly downregulated in ABCA7-KO iMGs and in the rs3752246 G/G risk allele in BR103 iMGs (**Figure 7M-N, S7F**).

To evaluate if the rs352246 ABCA7 AD risk variant affected microglial lysosomal function, we performed the DQ-BSA assay and found that lysosomal degradative capacity was reduced in BR103-G/G iMGs compared to their BR103-C/C corrected isogenic counterparts (**Figure 7O, S7G**). This functional impact also corresponded to a reduction in protein levels of the lysosomal peptidase CTSD, measured via Western blot (**Figure 7P-Q, S7H**). However, correcting the BR103 rs3752246 AD risk variant did not alter lysosomal area measured using lysotracker and LAMP1+ area (**Figure S7I-K**). Additionally, the rs3752246 correction in BR103 iMGs did not induce detectable changes in endocytosis, phagocytosis, or TREM2 expression (**Figure S7L-R**). Finally, we interrogated lipid-related functional measures of BR103 rs3752246-G/G risk-carrying microglia and found that peroxidized lipids and lipid droplet accumulation were not significantly different by rs3752246 genotype in iMGs (**Figure 7R-S, Figure S7S-T**). These data suggest that the risk variant alone is insufficient to produce detectable changes in some of the phenotypes observed in ABCA7-KO iMGs. In contrast, the lysosomal phenotypes remain robust, suggesting that partial impairment of ABCA7 may have a particularly strong impact on the microglial endolysosomal system.

### Microglial ABCA7 deficiency impacts AD-relevant neuronal phenotypes through non-autonomous mechanisms

Microglia impact the parenchymal cellular environment through a variety of direct and indirect mechanisms, including communicating with surrounding astrocytes and neurons. Since ABCA7 deficiency and AD risk variants both caused microglial phenotypes that included a primed inflammatory state and impaired lysosomal homeostasis, we next evaluated if and how ABCA7-KO microglia impacted other brain cells using a tri-culture method we previously described (38,47,48). Here, WT iAs and iNs were cultured with either ABCA7-WT or -KO iMGs to identify non-autonomous consequences of ABCA7 microglial deficiency using a human relevant microphysiological co-culture system (**Figure 8A-B**). While there was no effect on levels of secreted Aβ40, Aβ42, and Aβ38 (**Figure S8A-D**), a significant increase of phosphorylated tau (pTau) was observed (**Figure 8C-D, S8E**). Additionally, neurons in tri-cultures with ABCA7-KO iMGs had reduced synapse number (as measured by area of colocalized pre- and post-synaptic markers) compared to tri-cultures with ABCA7-WT iMGs (**Figure 8E-F**). This reduction in synapses corresponded to an overall reduction in VGLUT2+ pre-synaptic and HOMER1+ post-synaptic content (**Figure S8E-F**). Finally, tri-cultures with ABCA7-KO iMGs displayed reduced levels of extracellular APOE and CLU, AD-associated proteins primarily expressed by astrocytes and known to have roles in binding and transporting exported extracellular lipids (**Figure 8G-H**). Together, these data suggest that there is non-autonomous impact of ABCA7-deficient microglia on AD-related phenotypes on neural cell-types.

**Figure 8:**
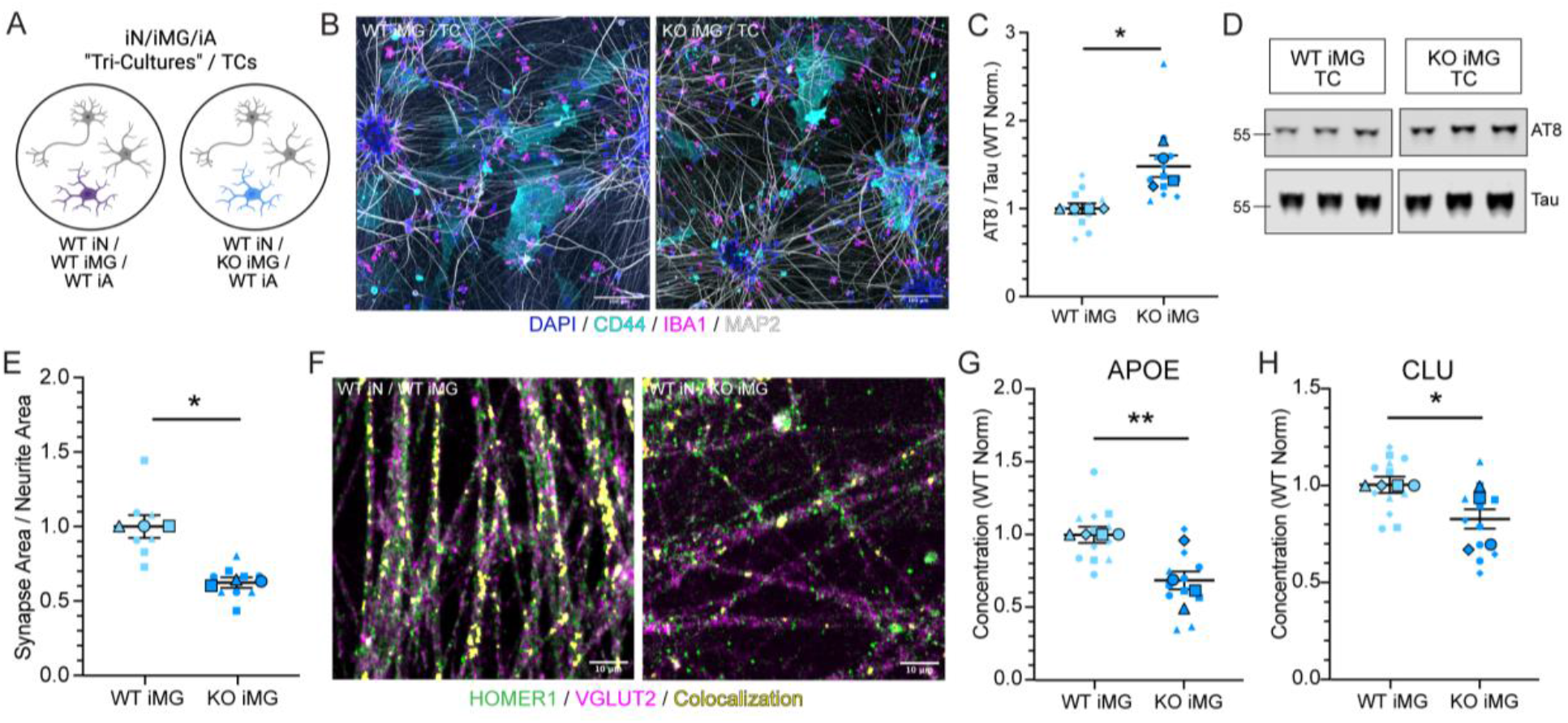
Microglial ABCA7 deficiency negatively impacts AD-relevant neuronal phenotypes in a microphysiological tri-culture system. (A) Schematic of tri-cultures with ABCA7-WT or -KO iMGs, WT iAs, and WT iNs. (B) Representative ICC images of DAPI, CD44, IBA1, and MAP2 in ABCA7-WT and -KO iMGs tri-cultures. Scale bar = 100µm. (C) Western blot analysis of Ser202/Thr205 phosphorylated Tau (AT8) and total Tau in ABCA7-WT and -KO iMG tri-cultures. Results were normalized to ABCA7-WT iMG tri-cultures within each differentiation. (D) Example western blot of AT8 and Tau in ABCA7-WT and -KO iMG tri-cultures. Numbers on the left denote relative molecular weights of protein standard (kDa). (E) Quantification of the area of colocalization of HOMER1+ and VGLUT2+ synapses divided by TUJ1+ neurite area for ABCA7-WT and -KO iMG tri-cultures. Results were normalized to ABCA7-WT iMG tri-cultures within each differentiation for visualization. (F) Representative ICC images of HOMER1, VGLUT2, and region of HOMER1/VGLUT2 colocalization identified in CellProfiler pipeline (see Methods). Images have been masked to only display TUJ1+ regions. Scale bar = 10µm. (G-H) Secreted APOE (G) and CLU (H) protein levels measured in conditioned media of tri-cultures with ABCA7-WT or -KO iMG. Results were normalized to WT = 1 within each differentiation for visualization. For all scatter plots, large, outlined points are representative of differentiation average and small points represent well-level replicates within differentiation. Different shapes denote separate differentiation. Statistical analyses display results of mixed-effects ANOVA on raw data (not normalized to WT) to capture within- and between-differentiation variance. Significant effect of genotype was determined: p<0.05 = *, p<0.01 = **, p<0.001 = ***, p<0.0001 = ****. Error bars = mean +/- SEM.

## DISCUSSION

Here, we employed unbiased proteomic profiling on differentiated human-derived iPSCs to identify molecular hallmarks that were altered by ABCA7 deficiency in three cell-types of the brain parenchyma: excitatory neurons, astrocytes and microglia. While the current body of literature supports a role for ABCA7 in regulating neuronal homeostasis (27–29), we demonstrate that of the three cell-types investigated, ABCA7 deficiency had the greatest impact on the microglia proteome. ABCA7 deficiency caused an impairment of microglial phagocytosis, lysosomal catabolism, accumulation of intracellular lipid droplets, consistent with the hypothesis that ABCA7 plays a key role in the efferocytosis pathway (82). Since the primary reported function of ABCA7 is in exporting lipids, including phospholipids (25,26,84,87) out of the cell, we interrogated the lipidome and proteome of ABCA7-deficient iMGs to identify the metabolic pathways altered by loss of this activity. First, we performed multi-omic analyses to replicate previous findings from human neuronal models that ABCA7 deficiency negatively impacts phospholipid remodeling (27) and promotes accumulation of intracellular oxidative stress through a possible accumulation of peroxidated lipids (24,28). This was complemented by novel findings that microglial ABCA7 deficiency promotes the accumulation of TGs and LDs. LD accumulation and functional lysosomal deficits induced by ABCA7-KO were found to be intertwined, as inhibiting LD accumulation using DGAT1/2 inhibitors rescued ABCA7-KO-induced lysosomal dysfunction. Concordantly, interrogation of the common *ABCA7* AD risk variant rs3752246 in variant-corrected isogenic iMGs further confirmed impaired lysosomal catabolism as a central hallmark of ABCA7 dysregulation. Taken together, these findings support a model in which ABCA7 dysfunction disrupts microglial lipid and lysosomal homeostasis, which may ultimately compromise the capacity of microglia to efficiently clear and process extracellular debris. This is consistent with mouse studies in AD models that show that ABCA7 deficiency leads to elevated plaque burden (97,98). Furthermore, tri-culture data suggest that microglial-derived ABCA7 may play a role in AD-relevant events such as suppressing the accumulation of phosphorylated tau and synapse loss.

An assumption of many functional genomics studies is that disease-related variants impact genes in the cell-types in which they exhibit the highest expression. While parenchymal *ABCA7* expression was initially thought to be highest in microglia and neurons (99), single cell/nuclear RNA sequencing studies have placed a greater emphasis on excitatory neurons due to the higher RNA expression of *ABCA7* in neurons (27,100). Investigation of a recently published snRNAseq dataset reveals that *ABCA7* is diffusely expressed across all excitatory neurons, as well as in subsets of microglia and astrocytes. Other subsets of microglia and astrocyte do not express detectable *ABCA7*, suggesting there could be regulated expression of cellular states in glial cell-types that affect *ABCA7* expression. Importantly, however, there are limitations to detecting glial transcriptomes using snRNAseq techniques, especially since much of the glial transcriptome is localized to the cytoplasm, particularly genes involved in microglia activation (101). Therefore, with previous evidence nominating microglia, astrocytes and excitatory neurons as primary expressors of ABCA7, we chose a data-driven unbiased approach to evaluate the molecular and functional impacts of ABCA7 deficiency on these cell-types using human iPSCs. We also chose to focus on the changes in proteomic compared to RNA abundance, as protein changes were more likely to reflect a cellular functional consequence.

The genetic architecture of AD has implicated microglial pathways involved in phagocytic clearance and efferocytosis, with numerous AD risk genes functioning in the recognition, uptake and intracellular processing of extracellular cargo (reviewed in 71). Our molecular and functional findings support an important role for ABCA7 in coordinating multiple steps within this process. ABCA7-deficient human microglia exhibited impaired phagocytosis of diverse extracellular substrates, disrupted lysosomal homeostasis, and accumulation of intracellular lipid droplets, suggesting defects in both cargo uptake and its subsequent intracellular processing. However, our TREM2 perturbation experiments argue against a simple model in which reduced phagocytosis directly drives the lysosomal phenotype: although both TREM2 antibodies reduced phagocytosis, they increased lysosomal activity. These findings instead suggest that ABCA7 deficiency may produce broader intracellular reprogramming of microglial lysosomal and lipid-handling pathways that secondarily influences phagocytic capacity, potentially through feedback mechanisms linking lipid metabolism, lysosomal function and cargo clearance.

Lipid droplets accumulate intracellularly when there is a disconnect of metabolic signals driving lipid production and lipid utilization (91). Although lipid droplets can serve a protective role by sequestering excess lipids, their chronic accumulation may become detrimental, particularly when they are enriched in polyunsaturated fatty acids (PUFAs) that are susceptible to ROS-mediated peroxidation (102). Consistent with this model, ABCA7-deficient iMGs exhibited increased PUFA-containing lipids and lipid droplets together with accumulation of peroxidized lipids and impaired endolysosomal function. Importantly, reducing lipid droplet accumulation by inhibiting the final step of triacylglycerol synthesis with DGAT1/2 inhibitors increased lysosomal activity in ABCA7-KO iMGs without altering phagocytosis. These findings suggest that excessive neutral lipid storage itself contributes to impaired lysosomal processing in microglia, and that reducing intracellular lipid burden can restore lysosomal function without requiring a reduction in the uptake of extracellular cargo.

This work along with a multitude of genetic and molecular evidence continue to highlight the importance of microglia lipid biology as a significant factor in AD pathogenesis (5,8,103). However, many genes with lipid-relevant function such as *ABCA7* remain under-investigated and functions within some cell-types unstudied. Lipids may be an amenable target for treating age-related neurodegeneration, and manipulating these pathways has shown promising initial results in related disorders such as Parkinson’s disease (104,105). Lipid droplet accumulation has been proposed as a central hallmark to microglia activation in AD (11,106), and data presented here suggest that alleviation of this pathology using DGAT1/2 inhibitors could assist with rescuing orthogonal deficits in lysosomal degradative phenotypes. However, complete genetic loss of DGAT1/2 in microglia in the PS19 mouse model exacerbated lysosomal lipid accumulation and hippocampal atrophy, indicating that caution is warranted when targeting this pathway while also recognizing that complete loss may have distinct consequence from partial pharmacological inhibition (107). Therefore, additional work is needed to understand this interplay between TG synthesis, lipid droplet accumulation, and lysosomal dysfunction in microglia under steady-state and AD-relevant pathological contexts.

Lipids and lipid-derived metabolites also serve as important plasma biomarkers for the impacts of AD risk variants on brain health. Evidence for this has been demonstrated through a recent metabolome wide association study (MWAS) that identified a relationship between LacCer plasma metabolites and the rs3752246 ABCA7 AD risk variant (108). These data corresponded to alterations in ceramide, HexCer, and LacCer species in *Abca7*-deficient mouse brains (108), suggesting that metabolic changes caused by loss-of-function mutations could be reflected in the periphery. We complemented this study and leveraged a large metabolite dataset derived from Finnish donors to identify two LacCer metabolites that accumulate in the plasma in individuals carrying the rs3752246 ABCA7 risk variant, which corresponded to proteomic deficits in LacCer degradation in rs3752246 risk allele-carrying microglia (76). These findings together support using plasma metabolomes for biomarker discovery of microglial metabolic activity, proposing a translational method for identifying the disease activity of an individual with a lipid-related AD risk variant, and possibly to monitor the progress of disease-modifying therapeutics.

The synthesis of findings identifying important molecular pathways and processes driven by complete ABCA7 deficiency, coupled to a targeted functional interrogation of a common *ABCA7* AD genetic risk factor has significant implications for the functional understanding of AD pathological progression. Previous human studies have associated the rs3752246 *ABCA7* AD risk variant with increased Aβ accumulation by leveraging data from longitudinal PET imaging of Aβ pathological accumulation. Earlier work found that the rs3752246 risk variant is associated with increased amyloid load in early-stage but not late-stage AD patients (109), proposing amyloid accumulation occurs earlier when the variant is present. Recent work has supported these findings, which identified the rs3752246 risk variant as significantly associated with an earlier age estimate of amyloid positivity, suggesting that a risk variant-carrying individual accumulates Aβ pathology earlier than their statistically predicted trajectory (110). These findings are supported by studies of the rare variable non-tandem repeat (VNTR) which is thought to cause a premature truncation of ABCA7. In a small cohort of individuals diagnosed with AD, VNTR length (associated with higher disease penetrance) was inversely correlated with Ab1-42, and positively correlated with pTau181 in cerebrospinal fluid (CSF) (111).

The data presented herein nominate microglia as a key cellular mediator of ABCA7-associated AD risk. Whereas ABCA7-KO iMGs exhibited clear deficits in phagocytosis, iMGs carrying the rs3752246 risk variant maintained phagocytic capacity under basal conditions. However, the rs3752246 risk allele-carrying iMGs exhibited measurable lysosomal deficits, suggesting that impaired intracellular processing may precede detectable defects in cargo uptake. We speculate that lysosomal dysfunction must exceed a critical threshold before phagocytic capacity becomes compromised. Further, we recognize that the rs3752246 iMGs leveraged in this current study were homozygous for both risk (G/G) or corrected neutral (C/C) alleles, which is less common in the population than heterozygosity, and this study has not determined if heterozygous iMGs would show an intermediate phenotype.

Consistent with a context-dependent effect, our tri-culture studies revealed no detectable changed in extracellular Aβ accumulation, although these experiments were performed under relatively homeostatic conditions in the absence of Aβ plaques or fibrils. Increasing Aβ burden or other age-associated cellular stresses may further challenge the lysosomal and lipid-handling defects produced by ABCA7 risk variants, eventually exceeding compensatory mechanisms that remain sufficient in microglia carrying this missense variant. This possibility is particularly relevant to AD as an age-related disorder, in which the functional consequences of variants affecting microglial activation and processing may emerge most prominently in the context of inflammatory, metabolic, or pathological stress. Similarly, non-cell-autonomous consequences of microglial ABCA7 impairment may require additional lipid, inflammatory or disease-associated challenge to become more apparent, although our data do suggest some subtle phenotypes associated with synapse remodeling and lipid transport proteins involved in regulating brain health.

## CONCLUSIONS

Taken together, this work identifies ABCA7 as a critical regulator of microglial lysosomal and lipid homeostasis, both important molecular events associated with AD. By leveraging multiple complementary human genetic models, we define convergent consequences of ABCA7 loss-of-function and identify potentially targetable metabolic pathways linking lipid metabolism, lysosomal function, and phagocytic activity. Manipulation of TREM2 signaling and triacylglycerol synthesis further demonstrates that these phenotypes can be functionally modified, highlighting metabolic pathways that may ultimately be leveraged therapeutically in the context of ABCA7-associated AD genetic risk. Furthermore, we provide initial evidence that microglial ABCA7 deficiency negatively impacts the parenchymal cellular environment and induces negative neuronal events that are relevant to AD, possibly through negative crosstalk with astrocyte-derived lipid transporting proteins APOE and CLU. Additional work to understand how ABCA7-deficient microglia are impacting their surrounding microenvironment, as well as how *ABCA7* risk variants interact with other common AD genetic risk factors such as *APOE*, *INPP5D* and *CLU* will further benefit the field in gaining a greater understanding of the benefits for therapeutically correcting lipid homeostasis mechanisms in AD.

## LIST OF ABBREVIATIONS

AD: Alzheimer’s disease
LOAD: late onset Alzheimer’s disease
fAD: familial Alzheimer’s disease
ABCA7: ATP Binding Cassette Subfamily A Member 7
iPSCs: induced pluripotent stem cells
WT: wild type
KO: knockout
iAs: induced astrocytes
iNs: induced neurons
iMGs: induced microglia
GWAS: genome wide association studies
Aβ: amyloid beta
SNP: single nucleotide polymorphism
PBMC: peripheral blood mononuclear cells
sgRNA: guide RNA
DIA-MS: data independent acquisition mass spectrometry
adj.P: adjusted p-value
HPN: human proteostasis network
TBS: tris buffered saline
TBST: tris buffered saline with tween
PBS: phosphate buffered saline
PFA: paraformaldehyde
BSA: bovine serum albumin
FBS: fetal bovine serum
HBSS: hanks balanced salt solution
ANs: apoptotic neurons
UV: ultraviolet
CM: crude myelin
PM: purified myelin
EDTA: ethylenediaminetetraacetic acid
HEPES: 4-(2-hydroxyethyl)-1-piperazineethanesulfonic acid
SLA: shotgun lipidomics assistant
LC-MS: liquid chromatography-mass spectrometry
DALS: differentially abundant lipids
log_2_FC: log2 fold change
LION: lipid MAPS Ontology
NES: normalized enrichment score
LSEA: lipid set enrichment analysis
PCA: principal component analysis
BioPAN: bioinformatics methodology for pathway analysis
BODIPY: boron-dipyrromethene
LD: lipid droplet
ROSMAP: religious orders study memory and aging project
DEP: differentially expressed protein
GSEA: gene set enrichment analysis
CTSD: cathepsin D
ASAH1: acid ceramidase
GNS: N-acetylglucoasmine-6-sulfatase
DQ-BSA: DQ-Red Bovine Serum Albumin
ICC: immunocytochemistry
Baf: bafilomycin
TREM2: triggering receptor expressed on myeloid cells 2
LAMP1: lysosome-associated membrane protein 1
CytoD: Cytochalasin D
HexCer: hexosylceramide
LPC: lysophosphatidyl choline
Cer: ceramide
SM: sphingomyelin
BMP: bis(monoacylglycerol)phosphate
PE: phosphatidylethanolamine
PE P: phosphatidylethanolamine plasmalogen
CE: cholesterol ester
TG: triglyceride
DG: Diacylglycerol
PC: phosphatidylcholine
PLD: phospholipase
LPCAT: lysophopsphatidylcholine acyltransferase
DGAT1/2: diacylglycerol O-acyltransferase 1 and 2
PA: phosphatidic acid
ROS: reactive oxygen species
LacCer: lactosylceramide
TC: tri-culture
snRNAseq: single nuclear RNA sequencing
scRNAseq: single cell RNA sequencing
PUFA: polyunsaturated fatty acid
MWAS: metabolome wide association study
PET: positron emission tomography
VNTR: variable non-tandem repeat
CSF: cerebrospinal fluid
pTau: phosphorylated tau

## DECLARATIONS

### Ethics approval and consent to participate

Humans: All work with human cell lines were approved by BWH/MBG Institutional Review Boards (IRB #2015P001676). All studies involving human materials were conducted in accordance with the ethical principles of the Declaration of Helsinki.

Animals: All animal procedures were approved by the BWH/MGB Animal Care and Use Committee (IACUC approval #2016N000464) and were performed in accordance with the NIH Guide for the Care and Use of Laboratory Animals and the ARRIVE guidelines.

### Consent for Publication

All authors have approved the contents of this manuscript and provided consent for publication.

## AVAILABILITY OF DATA AND MATERIALS

The datasets generated in the current study are available in Supplementary Material. Raw DIA-MS proteomics data are available at www.synapse.org under the following IDs: syn64759236 (ABCA7-WT/KO iAs, iNs, iMGs), syn71758938 (BR103 iMGs). Raw LC-MS Lipidomics data (in nmol/g) have been added to the Neurolipid Atlas repository (https://neurolipidatlas.com/). All cell lines are available upon request to Tracy Young-Pearse (corresponding author).

## AUTHOR CONTRIBUTIONS

Conceptualization, S.E.H. and T.L.Y-P.; methodology, S.E.H., T.L.Y-P., G.T., F.A., D.D., C.K., M.G., R.V.D.K., N.T.S.; validation, S.E.H., G.T., A.M.; formal analysis, S.E.H., F.A., G.T., C.K., investigation, S.E.H., F.A., G.T., A.M., C.K., M.G., R.V.D.K., D.D.; resources, T.L.Y-P.; data curation, S.E.H., F.A., T.L.Y-P., N.T.S., R.V.D.K.; writing-original draft, S.E.H. and T.L.Y-P.; writing-review and editing, S.E.H., F.A., G.T., A.M., D.D., C.K., M.G., R.V.D.K., N.T.S., T.L.Y-P.; visualization, S.E.H., F.A., T.L.Y-P.; supervision, T.L.Y-P.; project administration, T.L.Y-P.; funding acquisition T.L.Y-P. and S.E.H.

## ACKNOWLEDGEMENTS

We thank the NeuroTechnology Studio at Mass General Brigham for providing the Andor Dragonfly 600 Spinning Disk confocal instrument access and consultation on data acquisition and analysis; The Ann Romney Center for Neurologic Diseases Flow Cytometry Core Facility at Mass General Brigham for providing the BD Biosciences Fortessa Flow Cytometer instrument access and consultation on data acquisition and analysis; The iPSC NeuroHub at Mass General Brigham for technical assistance; the Proteomics Core at Emory University for proteomic data acquisition, analysis and consultation; and members of the Young-Pearse lab for their input. Scientific illustrations in this work incorporate elements adapted from BioRender.

