## Supplemental Information for "The Alzheimer’s disease risk gene *ABCA7* regulates microglial lysosomal and lipid homeostasis and promotes phagocytosis to maintain neuronal health"

**GRAPHICAL ABSTRACT**

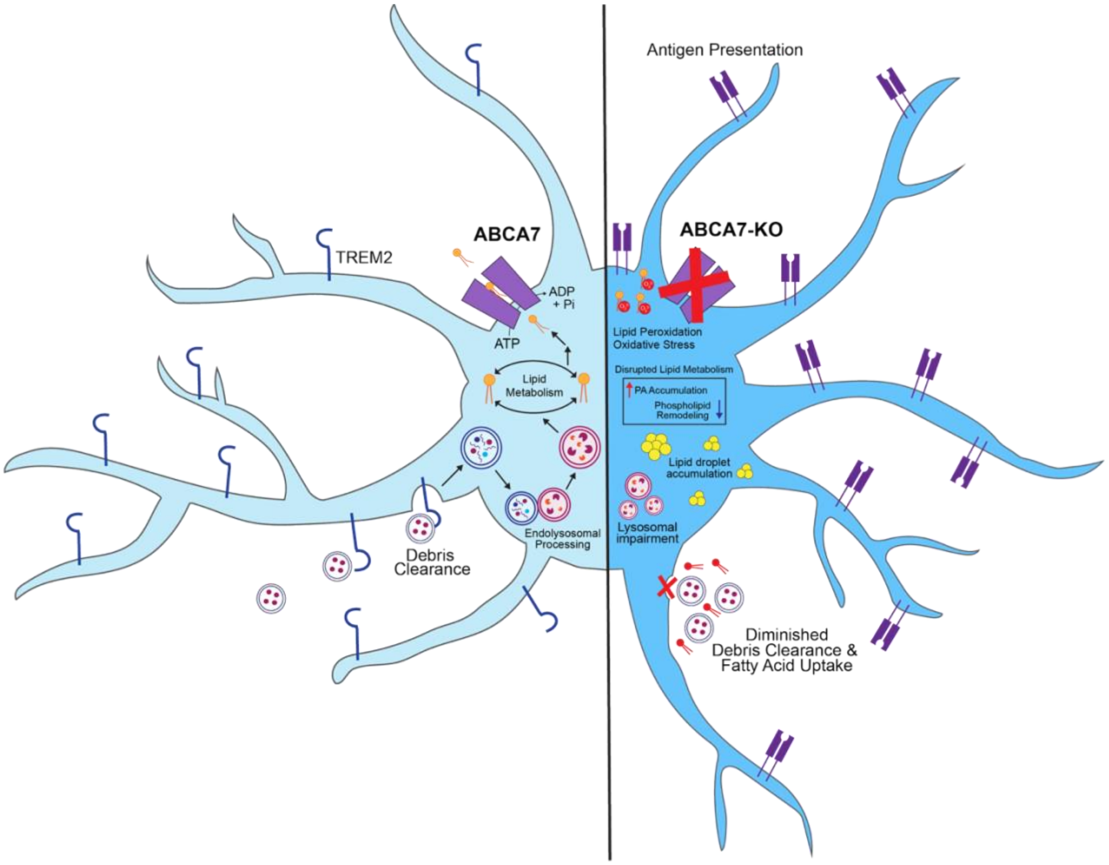


Under cellular homeostasis, microglial ABCA7 facilitates the removal of processed lipids through exporting via ATP hydrolysis. When ABCA7 is deficient, microglia accumulate peroxidized lipids and lipid droplets, shut down lysosomal function, and are unable to phagocytose extracellular debris. This is coupled with the loss of external TREM2 and upregulation of MHC antigen presentation proteins, suggesting a primed inflammatory state.

### SUPPLEMENTARY INFORMATION

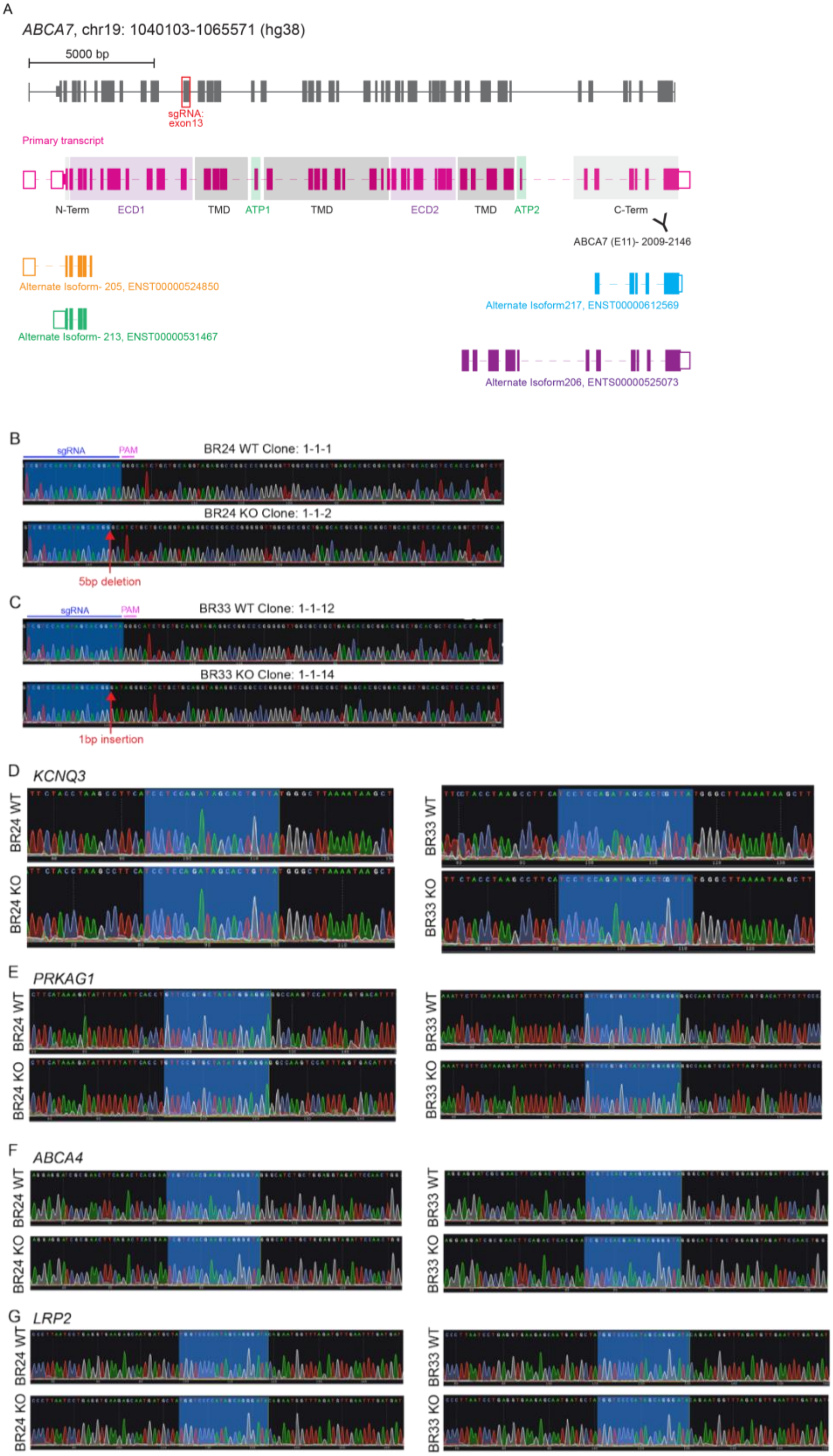

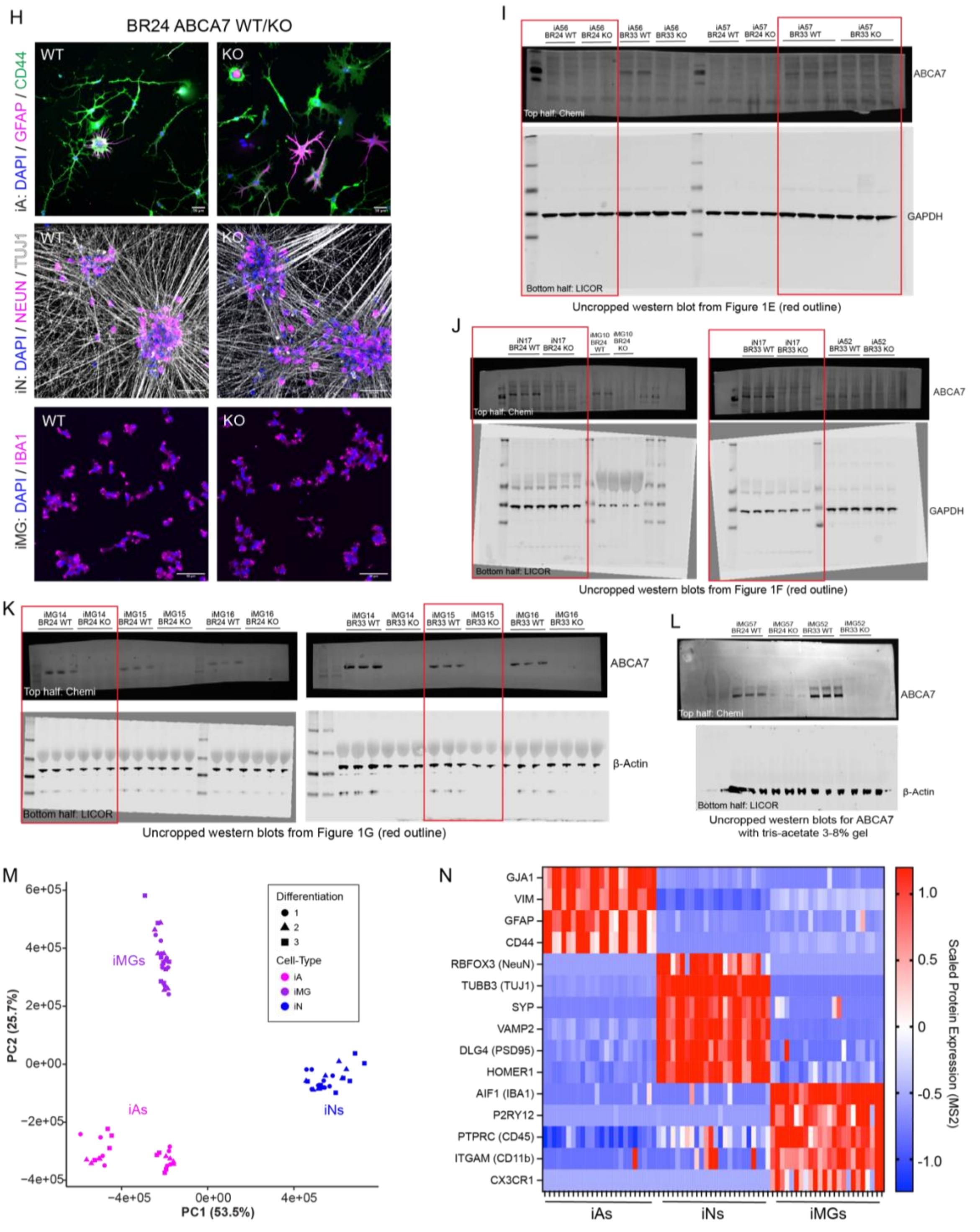
H
BR24 ABCA7 WT/KO
WT
KO
iA: DAPI / GFAP / CD44
iN: DAPI / NEUN / TUJ1
iMG: DAPI / IBA1
I
ABCA7
GAPDH
Top half: Chemi
Bottom half: LICOR
Uncropped western blot from Figure 1E (red outline)
J
ABCA7
GAPDH
Top half: Chemi
Bottom half: LICOR
Uncropped western blots from Figure 1F (red outline)
K
ABCA7
β-Actin
Top half: Chemi
Bottom half: LICOR
Uncropped western blots from Figure 1G (red outline)
L
ABCA7
β-Actin
Top half: Chemi
Bottom half: LICOR
Uncropped western blots for ABCA7 with tris-acetate 3-8% gel
M
iMGs
iAs
iNs
Differentiation
1
2
3
Cell-Type
iA
iMG
iN
PC1 (53.5%)
PC2 (25.7%)
N
GJA1
VIM
GFAP
CD44
RBFOX3 (NeuN)
TUBB3 (TUJ1)
SYP
VAMP2
DLG4 (PSD95)
HOMER1
AIF1 (IBA1)
P2RY12
PTPRC (CD45)
ITGAM (CD11b)
CX3CR1
iAs
iNs
iMGs
Scaled Protein Expression (MS2)

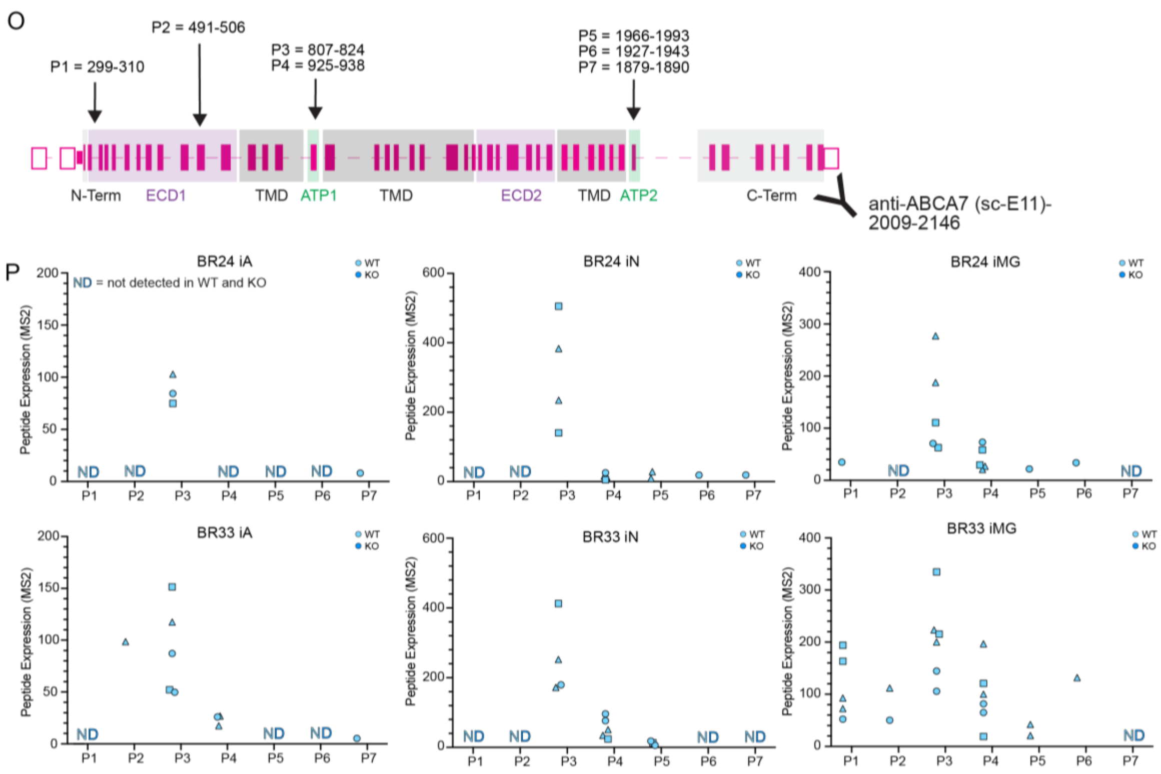


**Figure S1: Supplemental data corresponding to Figure 1**

(A) *ABCA7* gene structure with alternative isoforms and sgRNA location. Exons are denoted as thick, grey rectangles; red box relays the location of exon targeted for CRISPR/Cas9 to generate ABCA7 mutant iPSCs. Multi-colored structures below indicate known alternative isoforms of *ABCA7* and relative coding location on gene body.

(B-C) Sanger sequencing traces of BR24 (B) and BR33 (C) ABCA7-WT and ABCA7-KO iPSCs. Blue highlighted area denotes sgRNA region; magenta underline denotes PAM site. For ABCA7-KO clones the red area indicates region where insertion/deletion was made to generate a frame-shift mutation.

(D-G) Sanger sequencing traces of BR24 (left) and BR33 (right) ABCA7-WT (top) and ABCA7-KO (bottom) iPSCs for off-target effects of predicted genomic coding regions: KCNQ3 (D), PRKAG1 (E), ABCA4 (F) and LRP2 (G). Blue highlighted are denotes predicted sgRNA region if nucleotide mismatches occurred.

(H) Corresponding to Figure 1D. Representative ICC of differentiated iAs (top), iNs (middle) and iMGs (bottom) from BR24 ABCA7-WT (left) and ABCA7-KO (right) iPSCs. Scale bar = 50µm.

(I-K) Uncropped western blot corresponding to Figure 1E (I), Figure 1F (J), Figure 1G (K). Red outlined areas mark regions shown in the corresponding figure.

(L) Uncropped western blot of ABCA7 and β-actin for BR24 and BR33 ABCA7-WT and -KO iMGs run on a 3-8% tris-acetate gel to allow for separation of higher molecular weight bands.

(M) Principal component analysis of DIA-MS proteomics data of ABCA7-WT and -KO iMGs (purple), iAs (magenta) and iNs (blue). Symbol shapes correspond to differentiation.

(N) Heatmap of scaled protein abundance values for proteins with known high and/or exclusive expression within each cell-type. Rows correspond to proteins and columns correspond to individual samples. Samples are organized by cell-type (iAs, iNs, iMGs).

(O) *ABCA7* gene structure outlining exons corresponding to proteomic regions. Arrows indicate peptides captured by DIA-MS (P1-P7), and their corresponding ABCA7 amino acids. Y symbol

denotes the C-terminal binding region of ABCA7 antibody (Santa Cruz, sc-E11) used for the western blot analyses.
(P) Abundance of ABCA7 peptides denoted in S1N in BR24 (top) and BR33 (bottom) iAs (left), iNs (middle) and iMGs (right). Symbol shapes denote differentiation. ND = peptide was not detected in all samples. All ABCA7 peptides were not detected in ABCA7-KO samples.

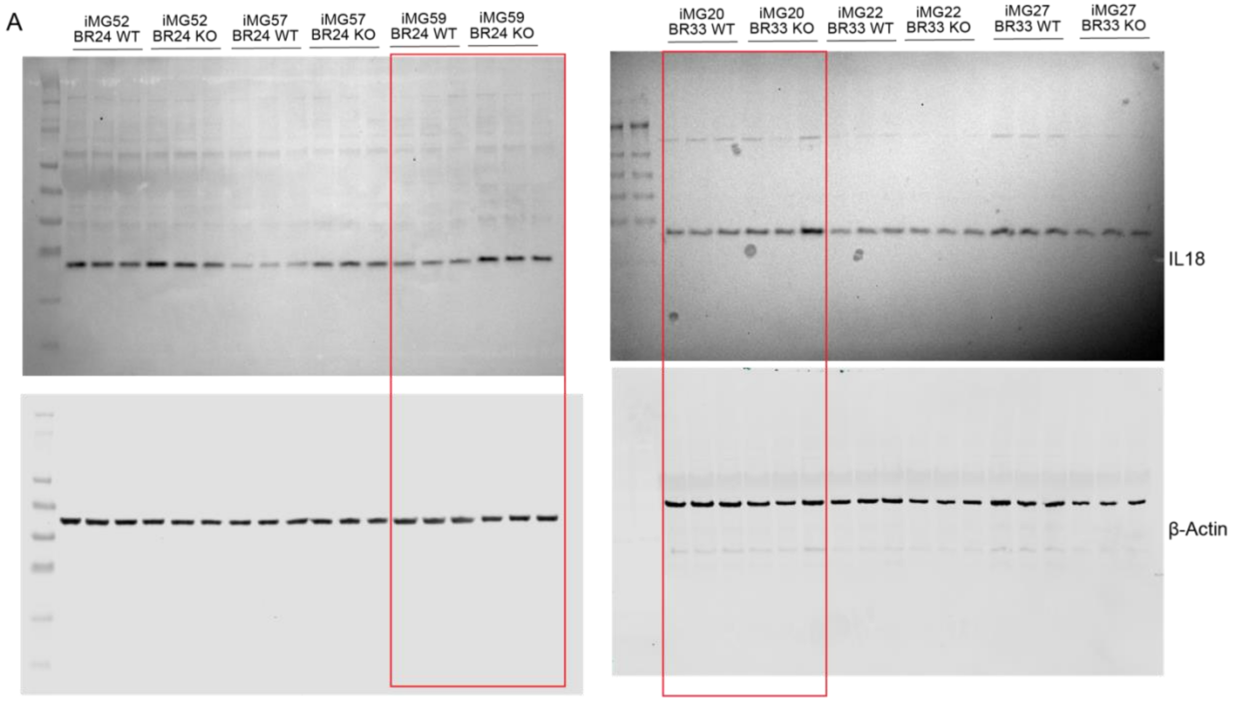

Uncropped western blots from Figure 2D (red outline)

**Figure S2: Supplemental data corresponding to Figure 2**.
(A) Uncropped western blot corresponding to Figure 2D. Red outlined areas mark regions shown in the corresponding figure.

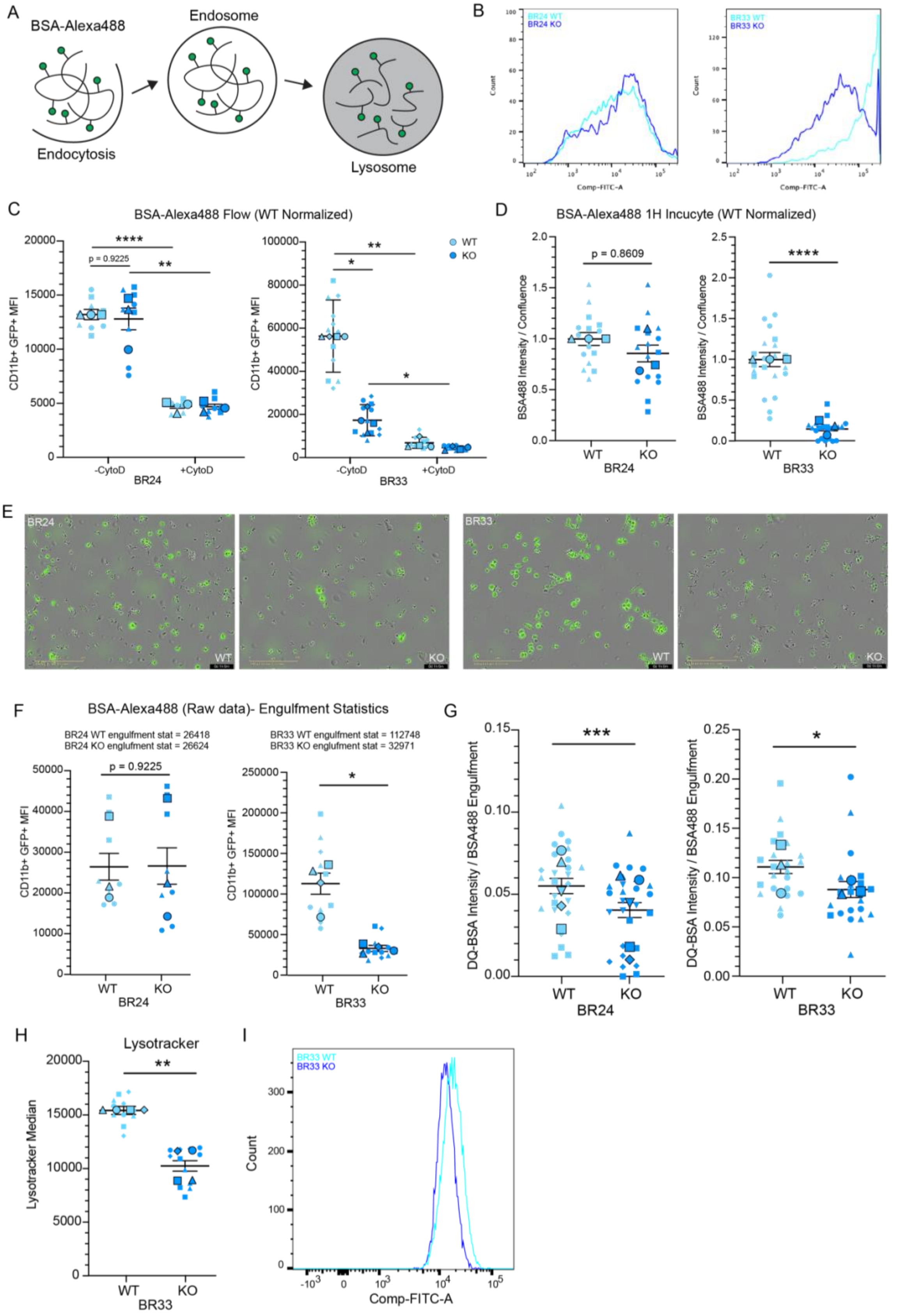
A
BSA-Alexa488
Endosome
Endocytosis
Lysosome
B
BR24 WT
BR24 KO
BR33 WT
BR33 KO
Count
Comp-FITC-A
C
BSA-Alexa488 Flow (WT Normalized)
CD11b+ GFP+ MFI
p = 0.9225
****
**
*
WT
KO
-CytoD
+CytoD
BR24
BR33
D
BSA-Alexa488 1H Incucyte (WT Normalized)
BSA488 Intensity / Confluence
p = 0.8609
****
WT
KO
BR24
BR33
E
BR24
BR33
WT
KO
F
BSA-Alexa488 (Raw data)- Engulfment Statistics
BR24 WT englufment stat = 26418
BR24 KO englufment stat = 26624
BR33 WT englufment stat = 112748
BR33 KO englufment stat = 32971
CD11b+ GFP+ MFI
p = 0.9225
*
WT
KO
BR24
BR33
G
DQ-BSA Intensity / BSA488 Engulfment
***
*
WT
KO
BR24
BR33
H
Lysotracker
Lysotracker Median
**
WT
KO
BR33
I
BR33 WT
BR33 KO
Count
Comp-FITC-A

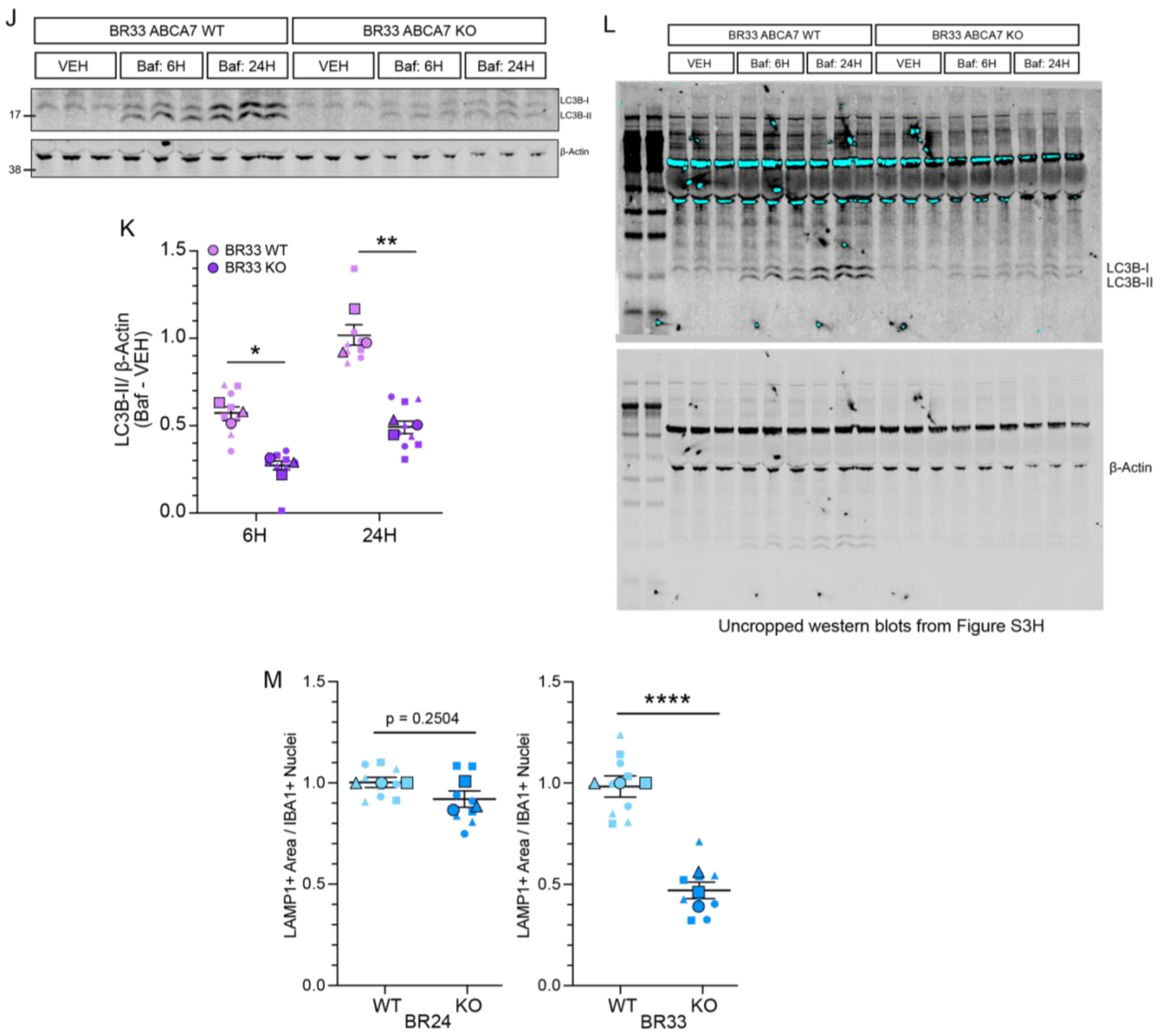


**Figure S3: Supplemental data corresponding to Figure 3.**

(A) Schematic of BSA488 treatment to measure endocytosis. Cells are treated or left untreated with the actin polymerization inhibitor cytochalasin D (CytoD) for 30 minutes, followed by treatment with BSA-Alexa488 substrate for 1 hour. Samples are then processed and analyzed for Alexa488/FITC fluorescence intensity to measure degree of engulfment.

(B) Example flow cytometry traces of BSA-488 (FITC) fluorescence intensity in gated CD11b+ BR24 (left) and BR33 (right) ABCA7-WT and -KO iMGs.

(C) Quantification of BSA488 mean fluorescence intensity (MFI) in CD11b+ BR24 (left) and BR33 (right) ABCA7-WT and -KO iMGs. Results were normalized to the WT average within each genetic background across all experiments for visualization.

(D) Quantification of BSA488 integrated intensity normalized to microglia confluence measured 1-hour post-treatment using Incucyte live cell imaging for BR24 (left) and BR33 (right) ABCA7-WT and -KO iMG. Results were normalized to the WT = 1 within each genetic background for each differentiation for visualization.

(E) Representative BSA488 images of BR24 (left) and BR33 (right) ABCA7-WT and -KO iMGs at the 1-hour post-treatment time-point. Scale bar = 200µm.

(F) Quantification of non-WT-normalized BSA488 MFI in CD11b+ BR24 (left) and BR33 (right) in ABCA7-WT and -KO iMGs. Mean MFI for each ABCA7-WT and -KO iMG cell line was calculated as a metric for endocytosis ("engulfment statistic") and listed above the graphs. These metrics were used for normalization of DQ-BSA intensity signal to quantify lysosomal activity based on amount of substrate engulfed by each cell line.

(G) DQ-BSA intensity normalized to engulfment statistics calculated in Figure S3F for ABCA7-WT and -KO iMG in BR24 (left) and BR33 (right) genetic backgrounds.

(H) Flow cytometric measurement of Lysotracker median fluorescence intensity in BR33 ABCA7-WT and -KO iMGs. Results were normalized to the WT average within each genetic background across all experiments for visualization.

(I) Example flow cytometry traces of Lysotracker (FITC) fluorescence intensity in BR33 ABCA7-WT and -KO iMGs.

(J) Representative western blot of LC3B-II accumulation as a measure of autophagic flux for BR33 ABCA7-WT and -KO iMGs treated with DMSO (VEH) or 100nM Bafilomycin for 6 or 24 hours. Numbers on the left denote relative molecular weights of protein standard (kDa).

(K) Analysis of autophagic flux LC3B-II accumulation in BR33 iMGs for 6 and 24 hours, subtracting baseline (VEH) LC3B-II levels from LC3B-II for each time-point. Results were normalized to the WT within each differentiation.

(L) Uncropped western blot corresponding to Figure S3J.

(M) Analysis of LAMP1+ area normalized to IBA1+ nuclei in BR24 (left) and BR33 (right) ABCA7-WT and -KO iMGs, corresponding to data depicted in Figure 3D. Results were normalized to WT = 1 within each differentiation.

For all scatter plots, large, outlined points are representative of differentiation average and small points represent well-level replicates within differentiation. Different shapes denote separate differentiation. Statistical analyses display results of mixed-effects ANOVA on raw data (not normalized to WT) to capture within- and between-differentiation variance. Significant effect of genotype was determined: $p<0.05$ = *, $p<0.01$ = **, $p<0.001$ = ***, $p<0.0001$ = ****. Error bars = mean +/- SEM.

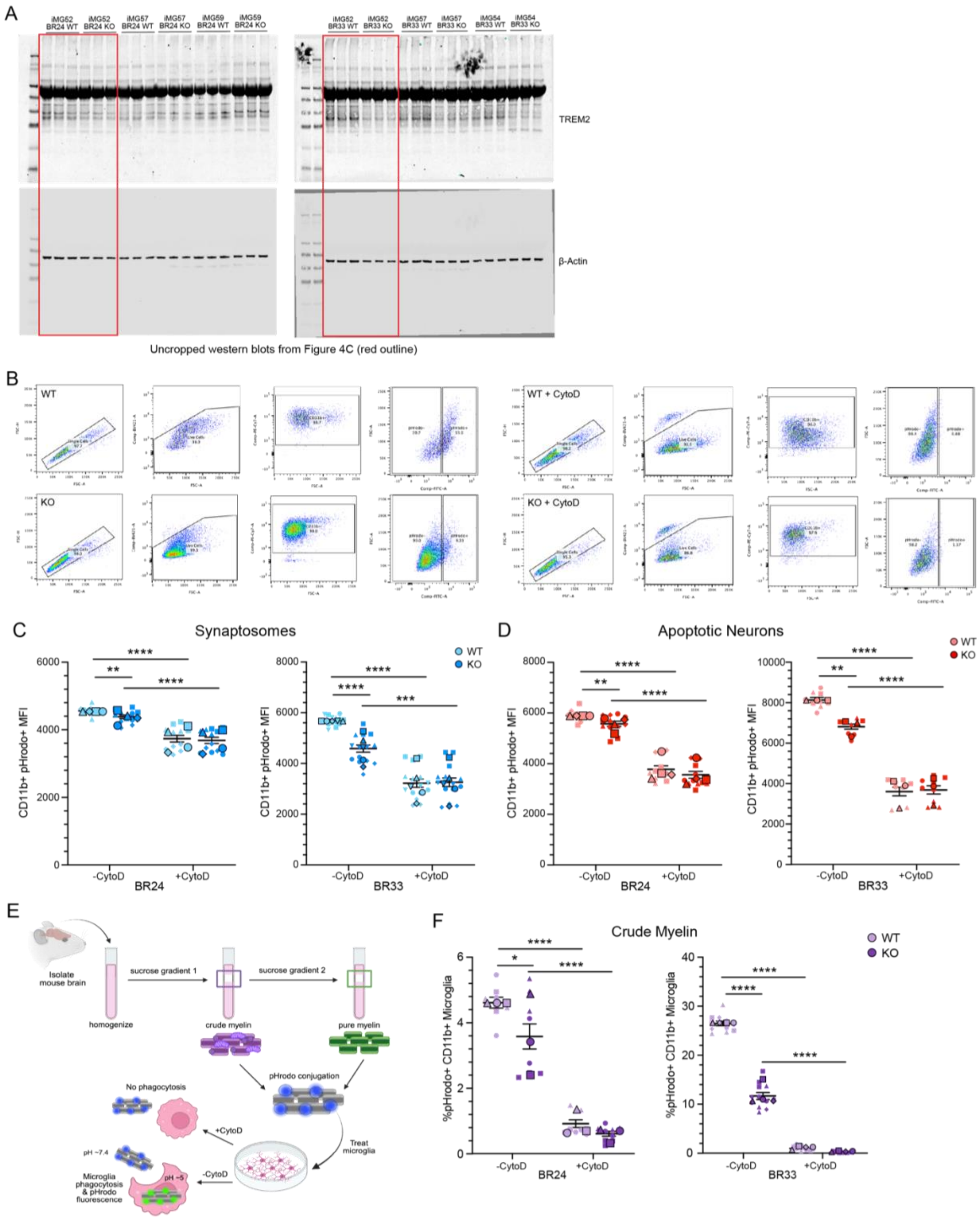
A
iMG52 BR24 WT
iMG52 BR24 KO
iMG57 BR24 WT
iMG57 BR24 KO
iMG59 BR24 WT
iMG59 BR24 KO
iMG52 BR33 WT
iMG52 BR33 KO
iMG57 BR33 WT
iMG57 BR33 KO
iMG54 BR33 WT
iMG54 BR33 KO
TREM2
β-Actin
Uncropped western blots from Figure 4C (red outline)
B
WT
WT + CytoD
KO
KO + CytoD
C
Synaptosomes
WT
KO
CD11b+ pHrodo+ MFI
-CytoD
+CytoD
BR24
BR33
D
Apoptotic Neurons
E
Isolate mouse brain
homogenize
sucrose gradient 1
crude myelin
sucrose gradient 2
pure myelin
pHrodo conjugation
Treat microglia
No phagocytosis
+CytoD
-CytoD
pH ~7.4
pH ~5
Microglia phagocytosis & pHrodo fluorescence
F
Crude Myelin
%pHrodo+ CD11b+ Microglia

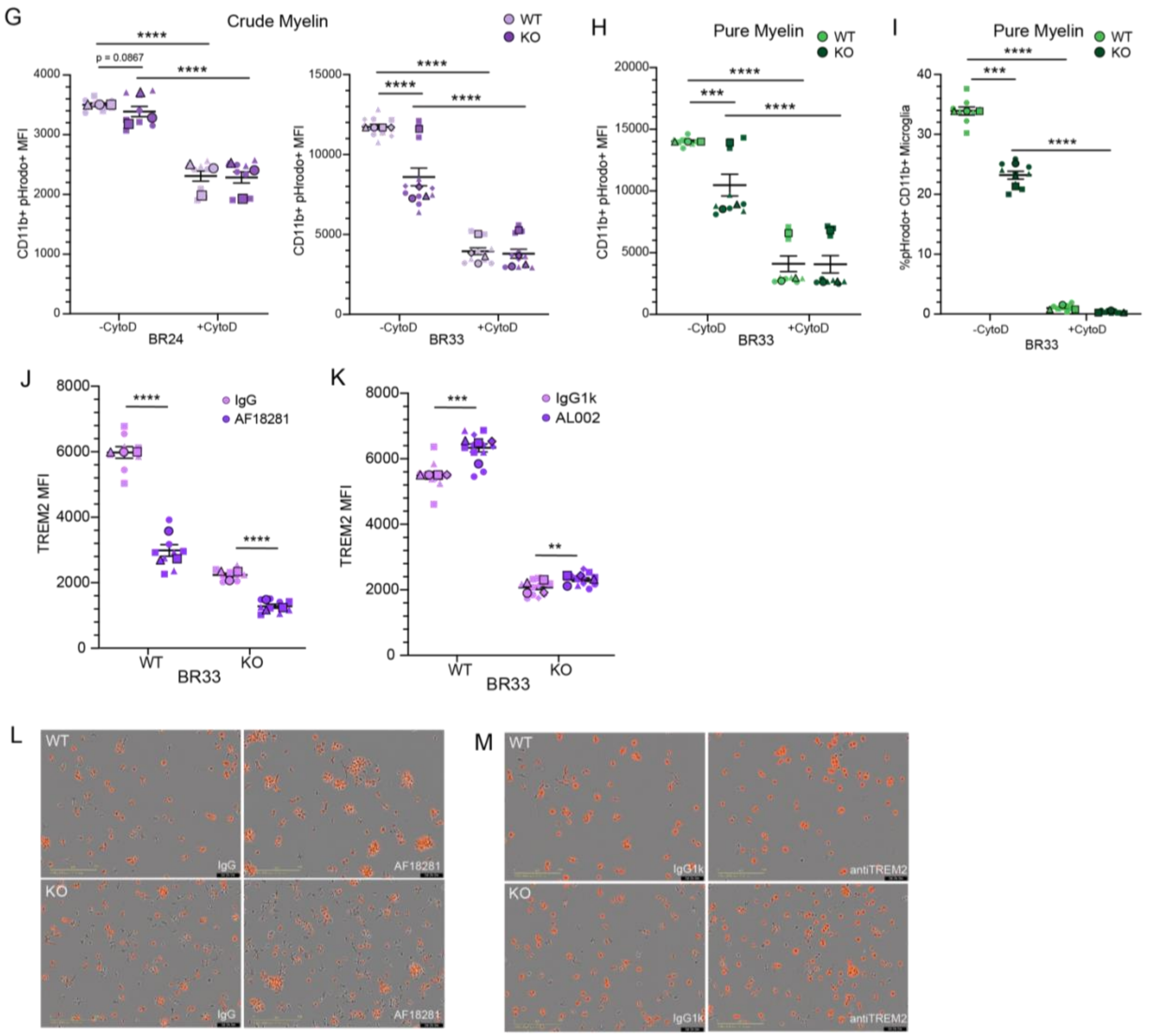


**Figure S4: Supplemental data corresponding to Figure 4**.

(A) Uncropped western blot corresponding to Figure 4C. Red outlined areas mark regions shown in the corresponding figure.

(B) Example gating strategy for flow cytometric analysis of pHrodo-conjugated substrates detailed in Figure 4. Initial gates are set to identify single cells using FSC-A and FSC-H, which is followed by isolation of live cells using Zombie Violet negative staining in the BV421 channel. CD11b+ microglia (PE-Cy7 channel) were isolated, and pHrodo+ cells (FITC channel) were determined using gating for a negative control (untreated cells) as well as confirmation with cells treated with CytoD.

(C-D) Flow cytometric analysis of iMG synaptosome (C) and AN (D) engulfment measured using CD11b+ pHrodo+ mean fluorescence intensity (MFI) for BR24 (left) and BR33 (right) iMGs. Results from each differentiation were normalized the WT average across all differentiations for visualization.

(E) Schematic of preparation and pHrodo-conjugation of crude and pure mouse myelin particles. For crude myelin, mouse brains were isolated and homogenized, then placed in a sucrose gradient followed by ultracentrifugation, with the interphase isolated and undergoing an osmotic shock. For pure myelin the same protocol was performed, with the myelin undergoing an

additional sucrose gradient after osmotic shocks to remove crude particles. Myelin was conjugated with pHrodo and provided to untreated or CytoD pre-treated microglia for 12-18 hours.
(F-G) Flow cytometric analysis of iMG crude myelin engulfment measured using %CD11b+pHrodo+ iMG (F) and CD11b+ pHrodo+ MFI (G) for BR24 (left) and BR33 (right) iMGs. Results from each differentiation were normalized the WT average across all differentiations.
(H-I) Flow cytometric analysis of iMG pure myelin engulfment measured using %CD11b+pHrodo+ iMG (H) and CD11b+ pHrodo+ MFI (I) for BR33 iMGs. Results from each differentiation were normalized the WT average across all differentiations for visualization.
(J-K) Flow cytometric analysis of TREM2 MFI on CD11b+ BR33 ABCA7-WT and -KO iMGs treated with AF18281 (J) or AL002 (K) anti-TREM2 antibodies. Results normalized to WT IgGk1/IgG average across all differentiations for visualization.
(L-M) Representative DQ-BSA images of BR33 ABCA7-WT and -KO iMGs treated with AF18281 (L) or AL002 (M) anti-TREM2 antibodies at the 2-hour post-chase time-point. Scale bar = 200µm.
For all scatter plots, large, outlined points are representative of differentiation average and small points represent well-level replicates within differentiation. Different shapes denote separate differentiation. Statistical analyses display results of mixed-effects ANOVA on raw data (not normalized to WT) to capture within- and between-differentiation variance. Significant effect of genotype was determined: $p<0.05$ = *, $p<0.01$ = **, $p<0.001$ = ***, $p<0.0001$ = ****. Error bars = mean +/- SEM.

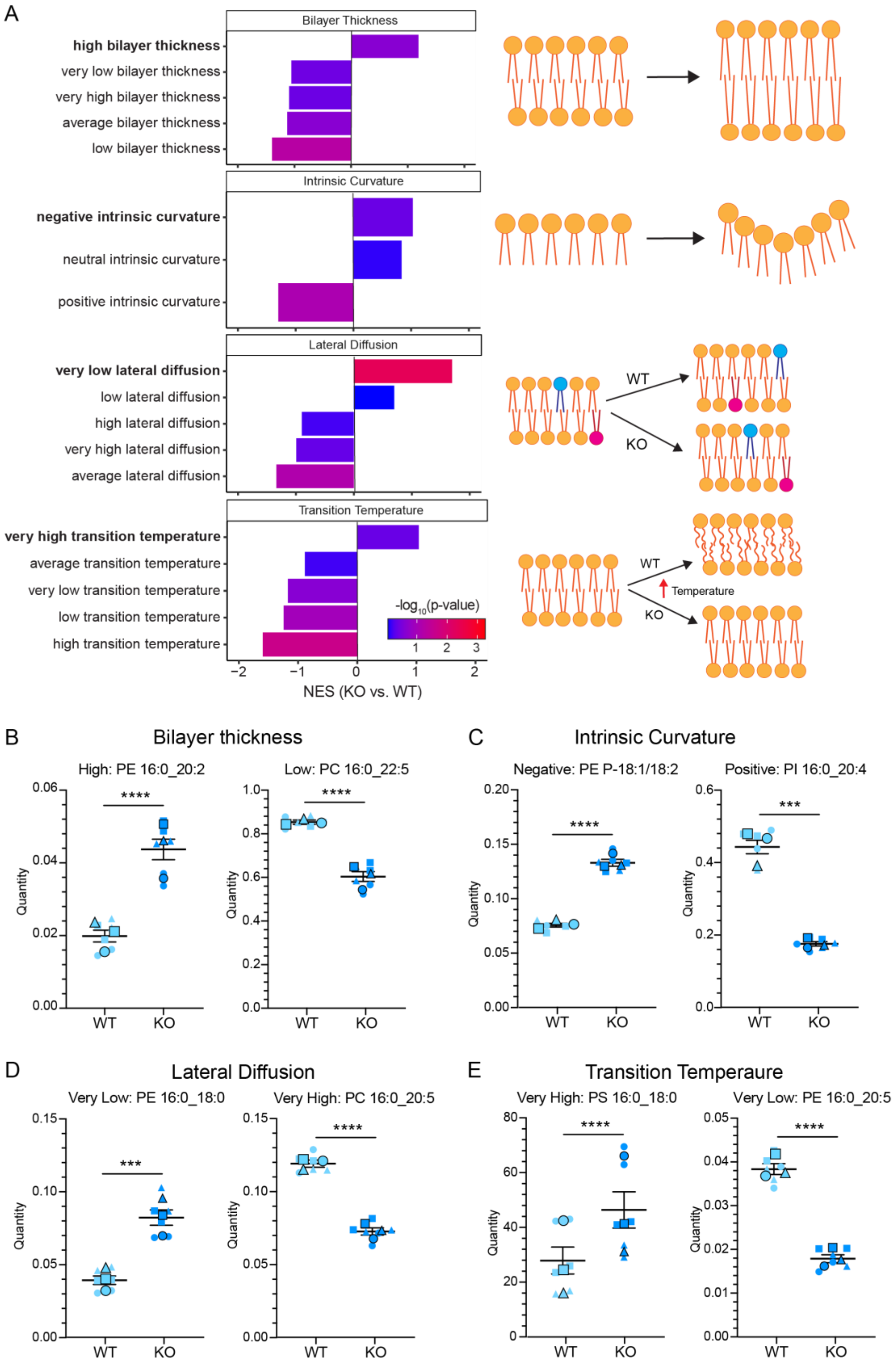

A
Bilayer Thickness
high bilayer thickness
very low bilayer thickness
very high bilayer thickness
average bilayer thickness
low bilayer thickness
Intrinsic Curvature
negative intrinsic curvature
neutral intrinsic curvature
positive intrinsic curvature
Lateral Diffusion
very low lateral diffusion
low lateral diffusion
high lateral diffusion
very high lateral diffusion
average lateral diffusion
WT
KO
Transition Temperature
very high transition temperature
average transition temperature
very low transition temperature
low transition temperature
high transition temperature
-log10(p-value)
Temperature
NES (KO vs. WT)
B
Bilayer thickness
High: PE 16:0_20:2
Low: PC 16:0_22:5
C
Intrinsic Curvature
Negative: PE P-18:1/18:2
Positive: PI 16:0_20:4
D
Lateral Diffusion
Very Low: PE 16:0_18:0
Very High: PC 16:0_20:5
E
Transition Temperaure
Very High: PS 16:0_18:0
Very Low: PE 16:0_20:5
Quantity
WT
KO

F

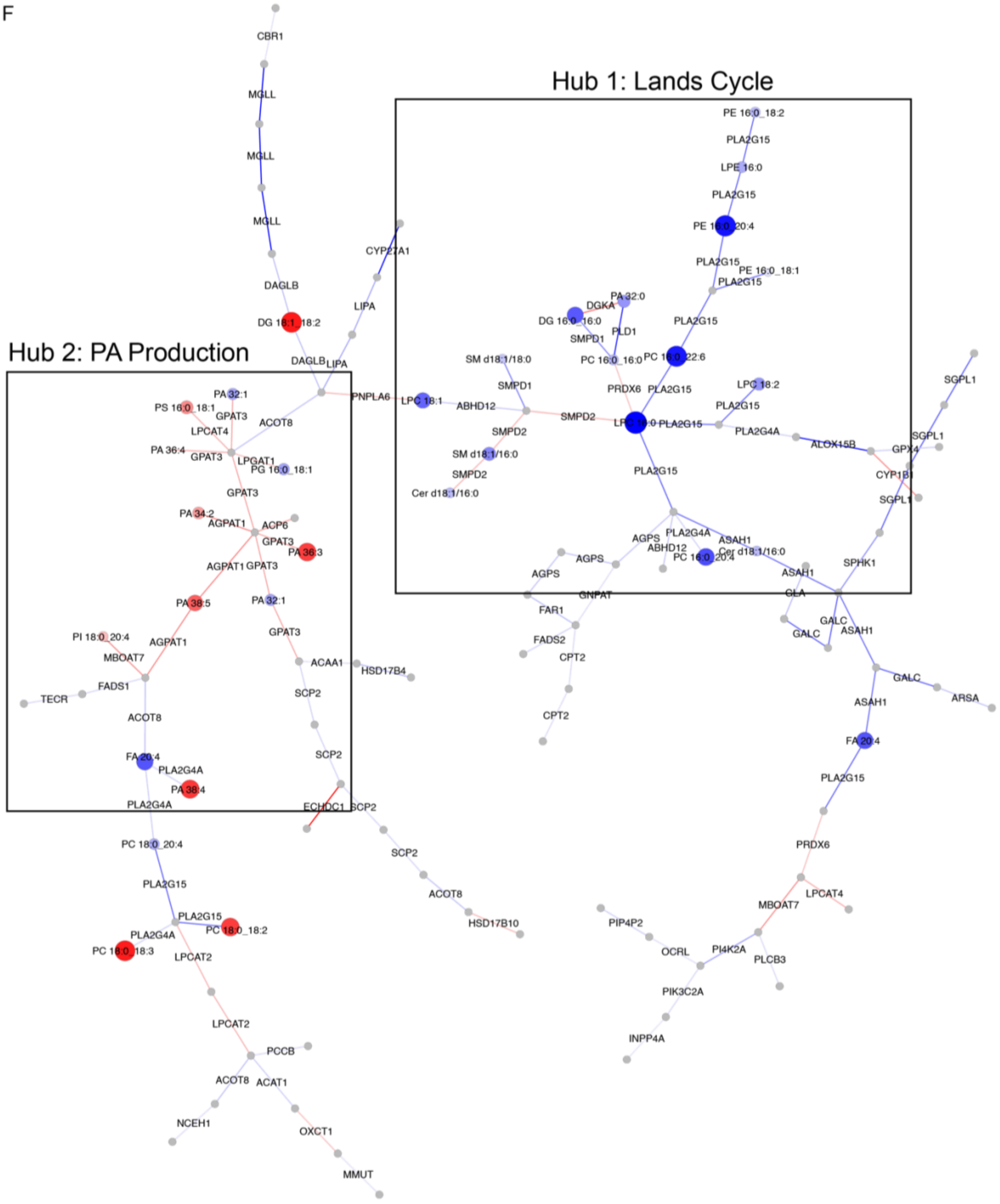
Hub 1: Lands Cycle
Hub 2: PA Production
CBR1
MGLL
MGLL
MGLL
DAGLB
DG 18:1_18:2
DAGLB
LIPA
CYP27A1
LIPA
PNPLA6
PE 16:0_18:2
PLA2G15
LPE 16:0
PLA2G15
PE 16:0_20:4
PLA2G15
PE 16:0_18:1
PLA2G15
PA 32:0
DGKA
DG 16:0_16:0
PLD1
SMPD1
PLA2G15
PC 16:0_16:0
PC 16:0_22:6
SM d18:1/18:0
SMPD1
PRDX6
PLA2G15
LPC 18:2
LPC 18:1
ABHD12
PLA2G15
SMPD2
LPC 16:0
PLA2G15
PLA2G4A
SMPD2
SM d18:1/16:0
SMPD2
Cer d18:1/16:0
ALOX15B
GPX4
SGPL1
SGPL1
CYP1B1
SGPL1
PLA2G15
AGPS
PLA2G4A
ASAH1
ABHD12
Cer d18:1/16:0
PC 16:0_20:4
AGPS
AGPS
ASAH1
SPHK1
GNPAT
GLA
FAR1
GALC
ASAH1
GALC
FADS2
CPT2
GALC
ARSA
ASAH1
CPT2
FA 20:4
PLA2G15
PRDX6
LPCAT4
MBOAT7
PIP4P2
OCRL
PI4K2A
PLCB3
PIK3C2A
INPP4A
PA 32:1
PS 16:0_18:1
GPAT3
ACOT8
LPCAT4
PA 36:4
GPAT3
LPGAT1
PG 16:0_18:1
GPAT3
PA 34:2
AGPAT1
ACP6
GPAT3
PA 36:3
AGPAT1
GPAT3
PA 38:5
PA 32:1
PI 18:0_20:4
AGPAT1
GPAT3
MBOAT7
ACAA1
HSD17B4
FADS1
TECR
SCP2
ACOT8
SCP2
FA 20:4
PLA2G4A
PA 38:4
PLA2G4A
ECHDC1
SCP2
SCP2
PC 18:0_20:4
ACOT8
PLA2G15
HSD17B10
PLA2G15
PC 18:0_18:2
PLA2G4A
PC 18:0_18:3
LPCAT2
LPCAT2
PCCB
ACOT8
ACAT1
NCEH1
OXCT1
MMUT

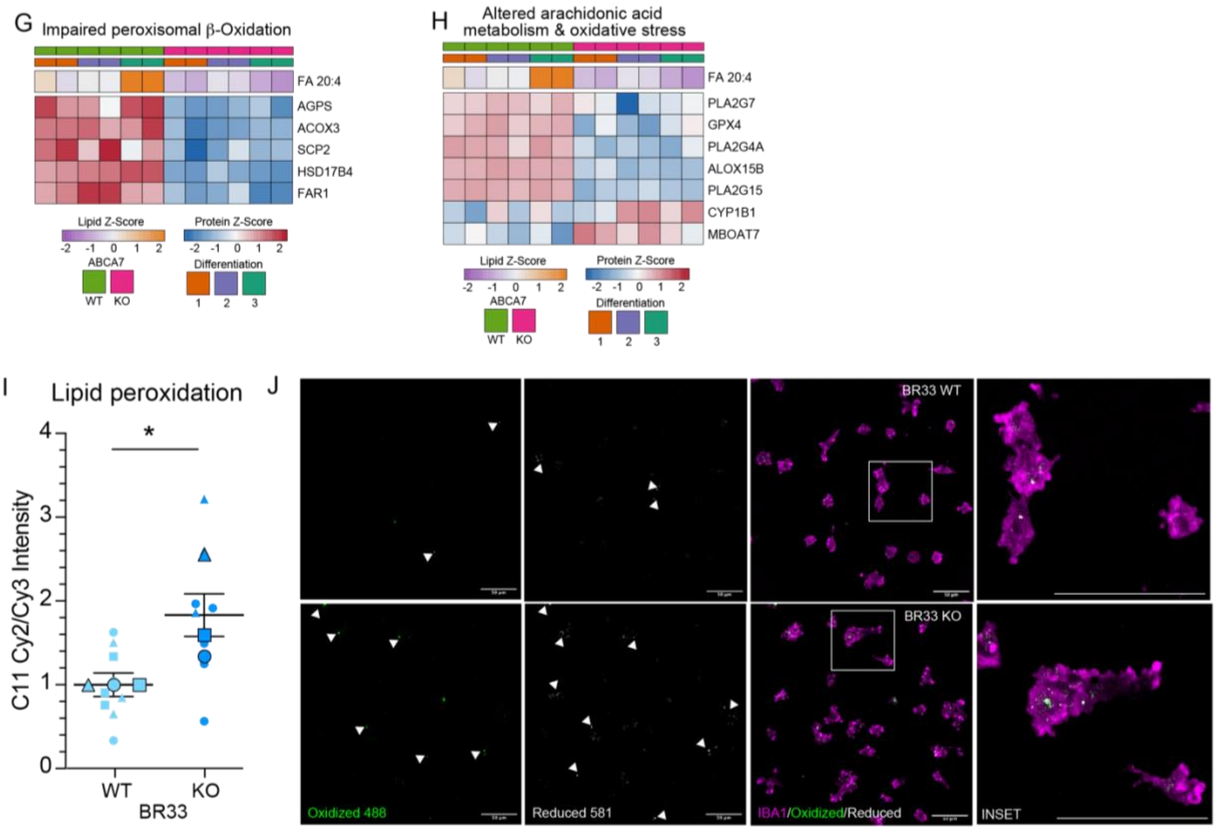


**Figure S5: Supplemental data corresponding to Figure 5**.

(A) Lipid set enrichment analysis (LSEA) of LION-annotated lipid biophysical characteristics to evaluate dynamics of lipids altered by ABCA7-KO in iMGs. Bars are organized by normalized enrichment score (NES) and colored by significance of enrichment (-$\log_{10}$ p-value). Bolded terms are significantly enriched characteristics between ABCA7-WT and -KO iMGs. Images to the right are schematics of predicted alterations to lipid dynamics based on significantly enriched characteristics.

(B-E) Scatter plots of leading-edge lipids used for analysis in S5A to evaluate biophysical characteristics of differentially abundant lipids between ABCA7-WT and -KO iMGs for bilayer thickness (B) intrinsic curvature (C) lateral diffusion (D) and transition temperature (E). Individual points are replicates used in lipidomic analysis, with symbols depicting replicates within each differentiation. Statistical significance is denoted from differential expression of lipids in Table S6 tab 3. All leading-edge lipids can be found in Table S6 tab 4. Values depicted are derived from Table S6 tab 2 for normalized abundance. Statistics depicted are derived from Table S6 tab 3, with significance levels representative of adjusted p-value testing difference of lipid species abundance between ABCA7-WT and -KO iMG.

(F) Enlarged image of the top interconnected metabolite-protein concordant sub-network (as represented in Figure 6A) identified through GATOM network analysis. Lipid nodes are colored and sized by most negative (blue) to positive (red) $\log_2$ fold change between ABCA7-KO and -WT iMGs. Grey nodes represent metabolite intermediaries not measured in the current dataset. Edges represent proteins involved in metabolite or lipid conversion and colored by $\log_2$ fold change between ABCA7-WT and -KO iMGs. Boxed areas are two manually selected central hubs of dysregulated protein-lipid interactions identified in the network.

(G-H) Heatmaps of lipid-protein pathways with known consequences from central GATOM hubs and containing altered lipid/protein abundance, binned into (G) impaired peroxisomal β-oxidation and (H) altered arachidonic acid metabolism/oxidative stress.

(I) Analysis of lipid peroxidation in BR33 ABCA7-WT and -KO iMGs using BODIPY 581/591-C11 fluorescent sensor. Peroxidation status was determined by measuring intensity of oxidized (Cy2, excitation/emission = 488/510) to reduced (Cy3, excitation/emission = 581/591) puncta in IBA1+ microglia. Results normalized to WT = 1 for each differentiation for visualization.
(J) Representative images of BR33 ABCA7-WT and -KO IBA1+ iMGs treated with BODIPY-C11 to measure oxidized (green) and reduced (white) lipids. Scale bar = 50µm.
Large, outlined points are representative of differentiation average and small points represent well-level replicates within differentiation. Different shapes denote separate differentiation. Statistical analyses display results of mixed-effects ANOVA of raw data (not normalized to WT) to capture within- and between-differentiation variance. Significant effect of genotype was determined: $p<0.05$ = *, $p<0.01$ = **, $p<0.001$ = ***, $p<0.0001$ = ****. Error bars = mean +/- SEM.

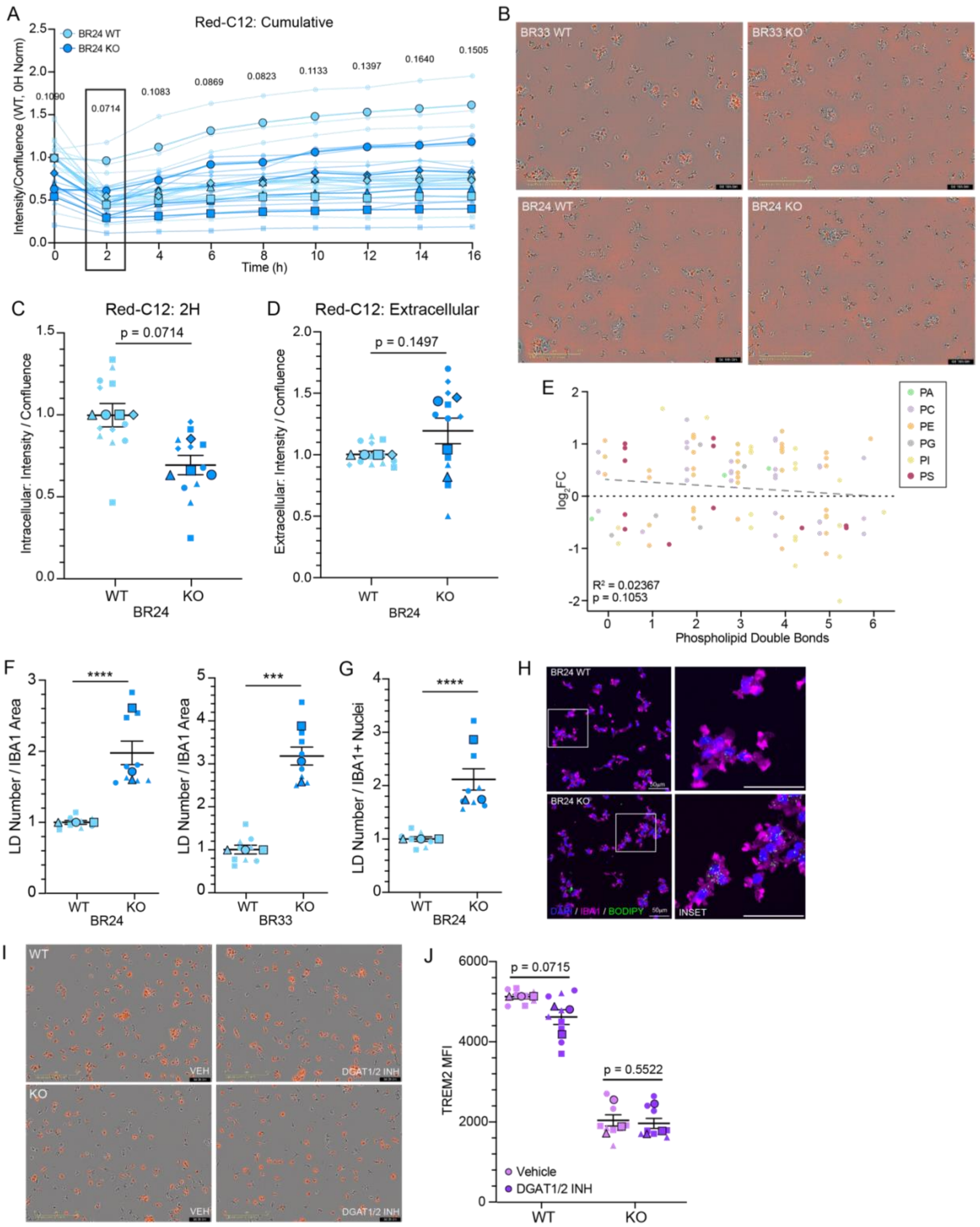


**Figure S6: Supplemental data corresponding to Figure 6.**

(A) Measurement of longitudinal intracellular signal of fatty acids labeled for 16 hours with BODIPY Red-C12 in BR24 ABCA7-WT and -KO iMGs. Solid, thick lines correspond to

differentiation averages, with thin, faint lines corresponding to individual replicates within each experiment. Results were normalized to WT = 1 for 0H time-point for visualization.
(B) Representative Incucyte image of intracellular lipids labeled with Red-C12 lipid sensor at exact time-point media was replaced with HBSS (16H) for extracellular chase to measure efflux. Scale bar = 200µm.
(C) Analysis of intracellular fatty acids 2 hours post-treatment with Red-C12 lipid sensor in BR24 ABCA7-WT and -KO iMGs. Results normalized to WT = 1 for each differentiation for visualization.
(D) Analysis of extracellular Red-C12 fatty acids 4 hours post-media replacement in BR24 ABCA7-WT and -KO iMGs. Results normalized to WT = 1 for each differentiation for visualization.
(E) Scatter plot of phospholipids classified by degree of unsaturation (number of double bonds) and $log_2FC$ abundance difference between ABCA7-WT and -KO iMGs. Horizontal line and error bars are mean $log_2FC$ and SD for lipids with each level of unsaturation. Points are colored by phospholipid class assigned in Figure 5C. Grey line denotes results from linear regression of double bond number to mean $log_2FC$ of TG abundance. $R^2$ and p-value of linear regression are denoted in upper left-hand corner of graph.
(F) Quantification of BODIPY+ lipid droplet number normalized to IBA1+ area in BR24 (left) and BR33 (right) ABCA7-WT and -KO iMGs. Results were normalized to the WT within each differentiation for visualization.
(G) Quantification of BODIPY+ lipid droplet number normalized to IBA1+ nuclei in BR24 ABCA7-WT and -KO iMGs. Results were normalized to the WT within each differentiation for visualization.
(H) Representative ICC images of DAPI, BODIPY and IBA1 in BR24 ABCA7-WT and -KO iMGs. Scale bar = 50µm.
(I) Representative DQ-BSA images of BR33 ABCA7-WT and -KO iMGs treated with DGAT1/2 inhibitor (DGAT1/2 INH) or DMSO vehicle (VEH) at the 2-hour post-chase time-point. Scale bar = 200µm.
(J) Flow cytometric analysis of TREM2 MFI on CD11b+ BR33 ABCA7-WT and -KO iMGs treated with DGAT1/2 inhibitor (DGAT1/2 INH) or DMSO vehicle (DMSO). Results normalized to WT VEH average across all differentiations for visualization.
Large, outlined points are representative of differentiation average and small points represent well-level replicates within differentiation. Different shapes denote separate differentiation. Statistical analyses display results of mixed-effects ANOVA of raw data (not normalized to WT) to capture within- and between-differentiation variance. Significant effect of genotype was determined: $p<0.05$ = *, $p<0.01$ = **, $p<0.001$ = ***, $p<0.0001$ = ****. Error bars = mean +/- SEM.

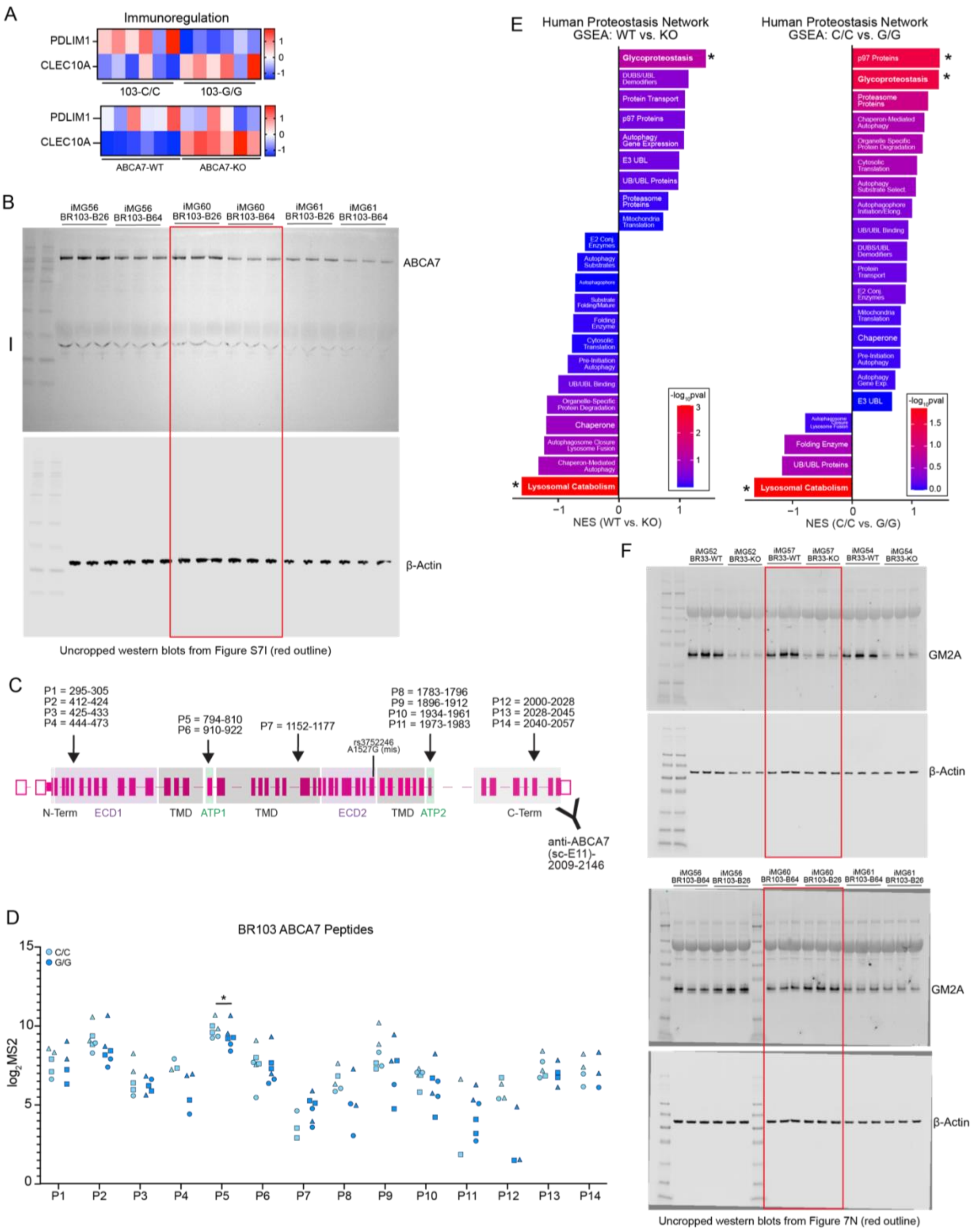

A
Immunoregulation
PDLIM1
CLEC10A
103-C/C
103-G/G
ABCA7-WT
ABCA7-KO
B
iMG56 BR103-B26
iMG56 BR103-B64
iMG60 BR103-B26
iMG60 BR103-B64
iMG61 BR103-B26
iMG61 BR103-B64
ABCA7
β-Actin
Uncropped western blots from Figure S7I (red outline)
C
P1 = 295-305
P2 = 412-424
P3 = 425-433
P4 = 444-473
P5 = 794-810
P6 = 910-922
P7 = 1152-1177
rs3752246
A1527G (mis)
P8 = 1783-1796
P9 = 1896-1912
P10 = 1934-1961
P11 = 1973-1983
P12 = 2000-2028
P13 = 2028-2045
P14 = 2040-2057
N-Term
ECD1
TMD
ATP1
TMD
ECD2
TMD
ATP2
C-Term
anti-ABCA7 (sc-E11)-2009-2146
D
BR103 ABCA7 Peptides
C/C
G/G
log2MS2
P1
P2
P3
P4
P5
P6
P7
P8
P9
P10
P11
P12
P13
P14
E
Human Proteostasis Network GSEA: WT vs. KO
Glycoproteostasis
DUBS/UBL Demodifiers
Protein Transport
p97 Proteins
Autophagy Gene Expression
E3 UBL
UB/UBL Proteins
Proteasome Proteins
Mitochondria Translation
E2 Conj. Enzymes
Autophagy Substrates
Autophagophore
Substrate Folding/Mature
Folding Enzyme
Cytosolic Translation
Pre-Initiation Autophagy
UB/UBL Binding
Organelle-Specific Protein Degradation
Chaperone
Autophagosome Closure Lysosome Fusion
Chaperon-Mediated Autophagy
Lysosomal Catabolism
-log10pval
NES (WT vs. KO)
Human Proteostasis Network GSEA: C/C vs. G/G
p97 Proteins
Glycoproteostasis
Proteasome Proteins
Chaperon-Mediated Autophagy
Organelle Specific Protein Degradation
Cytosolic Translation
Autophagy Substrate Select.
Autophagophore Initiation/Elong.
UB/UBL Binding
DUBS/UBL Demodifiers
Protein Transport
E2 Conj. Enzymes
Mitochondria Translation
Chaperone
Pre-Initiation Autophagy
Autophagy Gene Exp.
E3 UBL
Folding Enzyme
UB/UBL Proteins
Lysosomal Catabolism
NES (C/C vs. G/G)
F
iMG52 BR33-WT
iMG52 BR33-KO
iMG57 BR33-WT
iMG57 BR33-KO
iMG54 BR33-WT
iMG54 BR33-KO
GM2A
β-Actin
iMG56 BR103-B64
iMG56 BR103-B26
iMG60 BR103-B64
iMG60 BR103-B26
iMG61 BR103-B64
iMG61 BR103-B26
GM2A
β-Actin
Uncropped western blots from Figure 7N (red outline)

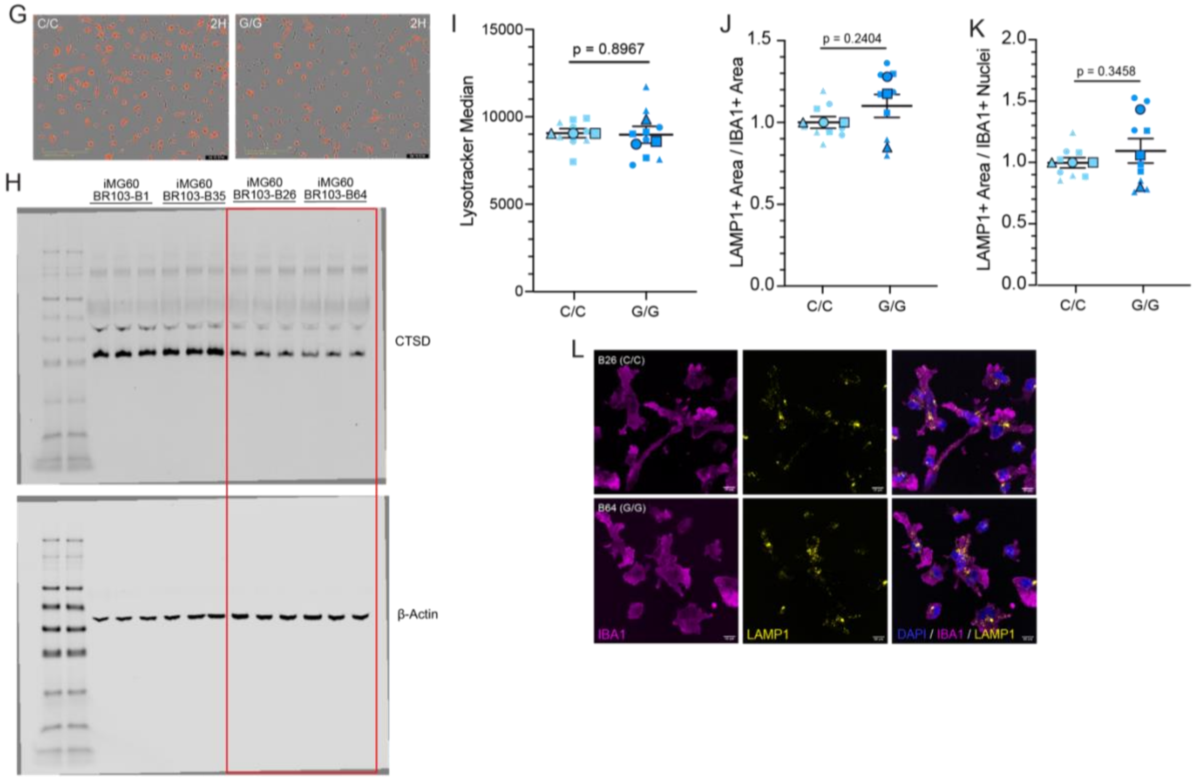


Uncropped western blots from Figure 7P (red outline)

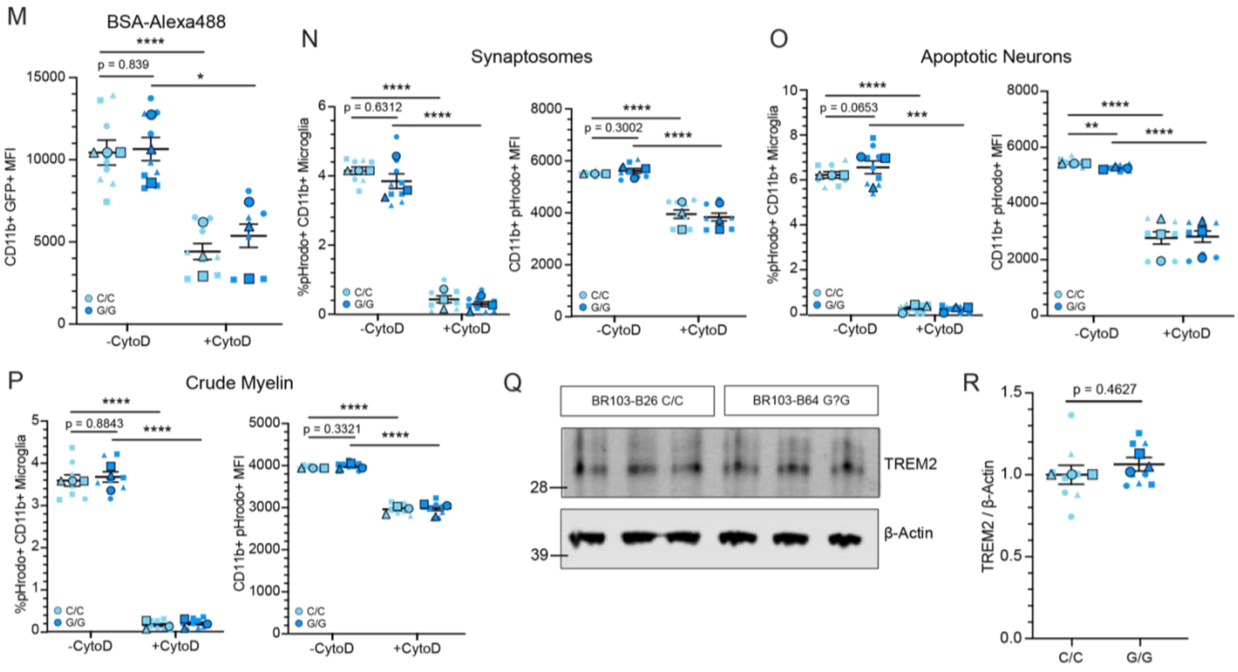

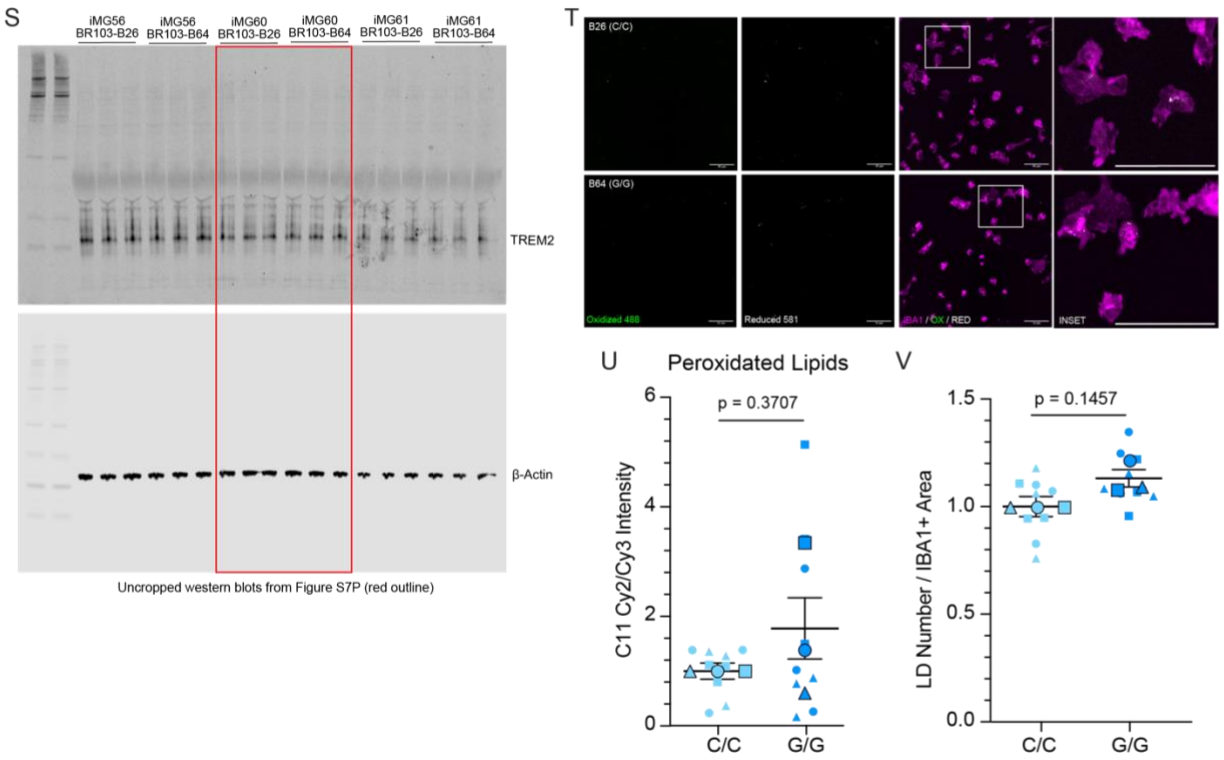


**Figure S7: Supplemental data corresponding to Figure 7**.

(A) Heatmap of scaled protein abundance values of shared immunoregulatory proteins that are differentially expressed in BR33 ABCA7-WT versus -KO (top) and BR103-C/C versus BR103-G/G (bottom) iMGs.

(B) Uncropped western blot corresponding to Figure 7I. Red outlined areas denote regions depicted in the corresponding figure.

(C) *ABCA7* gene structure outlining exons corresponding to proteomic regions. Arrows indicate peptides captured by DIA-MS (P1-P14), and their corresponding ABCA7 amino acids. Y symbol denotes the C-terminal binding region of ABCA7 antibody (Santa Cruz, sc-E11) used for western blot analyses.

(D) Abundance of ABCA7 peptides denoted in S7C in BR103 iMGs. Symbol shapes denote differentiation. ND = peptide was not detected in all samples. Asterisk denotes nominal significance of paired t-test between BR103-C/C and BR103-G/G iMGs.

(E) Results of GSEA preranked analysis using Human Proteostasis Network gene sets in ABCA7-KO versus -WT iMGs (left) and BR103-C/C versus BR103-G/G iMGs (right). Asterisks denote gene set with significant enrichment in dataset. Bolded terms denote gene sets with shared enrichment between both comparisons.

(F) Uncropped western blot corresponding to Figure 7N. Red outlined areas mark regions shown in the corresponding figure.

(G) Representative DQ-BSA images of BR103-C/C (left) and BR103-G/G (right) iMGs at the 2-hour post-chase time-point. Scale bar = 200μm.

(H) Uncropped western blot corresponding to Figure 7P. Red outlined areas mark regions shown in the corresponding figure.

(I) Flow cytometric measurement of Lysotracker median fluorescence intensity BR103 iMGs. Results were normalized to the BR103-C/C average across all differentiations for visualization.

(J-K) Analysis of LAMP1+ area normalized to IBA1+ area (J) and nuclei (K) in BR103 iMGs. Results were normalized to BR103-C/C within each differentiation for visualization.

(L) Representative ICC images of LAMP1 in IBA1+ microglia in BR103 iMGs. Scale bar = 10μm.

(M) Quantification of BSA-488 MFI in CD11b+ BR103 iMGs. Results were normalized to BR103-C/C average across all experiments for visualization.
(N-P) Flow cytometric analysis of pHrodo-conjugated synaptosome (N), apoptotic neuron (O) and crude myelin (P) engulfment for BR103 iMGs. Results from each differentiation were normalized the BR103-C/C average across all differentiations for visualization.
(Q) Example western blot of TREM2 and β-actin in BR103 iMGs. Left numbers denote relative molecular weights of protein standard (kDa).
(R) Western blot analysis of TREM2 expression in BR103 iMGs. Results normalized to BR103-C/C for each differentiation for visualization.
(S) Uncropped western blot corresponding to Figure S7Q. Red outlined areas mark regions shown in the corresponding figure.
(T) Representative images of BR103 IBA1+ iMGs treated with BODIPY-C11 to measure oxidized (green) and reduced (white) lipids. Scale bar = 50µm.
(U) Analysis of lipid peroxidation in BR103 iMGs using BODIPY 581/591-C11 fluorescent sensor. Peroxidation status was determined by measuring intensity of oxidized (Cy2, excitation/emission = 488/510) to reduced (Cy3, excitation/emission = 581/591) puncta in IBA1+ microglia. Results normalized to BR103-C/C = 1 for each differentiation for visualization.
(V) Quantification of BODIPY+ lipid droplet number normalized to IBA1+ area in BR103 iMGs. Results were normalized to BR103-C/C = 1 within each differentiation for visualization.
For all scatter plots, large, outlined points are representative of differentiation average and small points represent well-level replicates within differentiation. Different shapes denote separate differentiation. Statistical analyses display results of mixed-effects ANOVA of raw data to capture within- and between-differentiation variance. Significant effect of genotype was determined: $p<0.05$ = *, $p<0.01$ = **, $p<0.001$ = ***, $p<0.0001$ = ****. Error bars = mean +/- SEM.

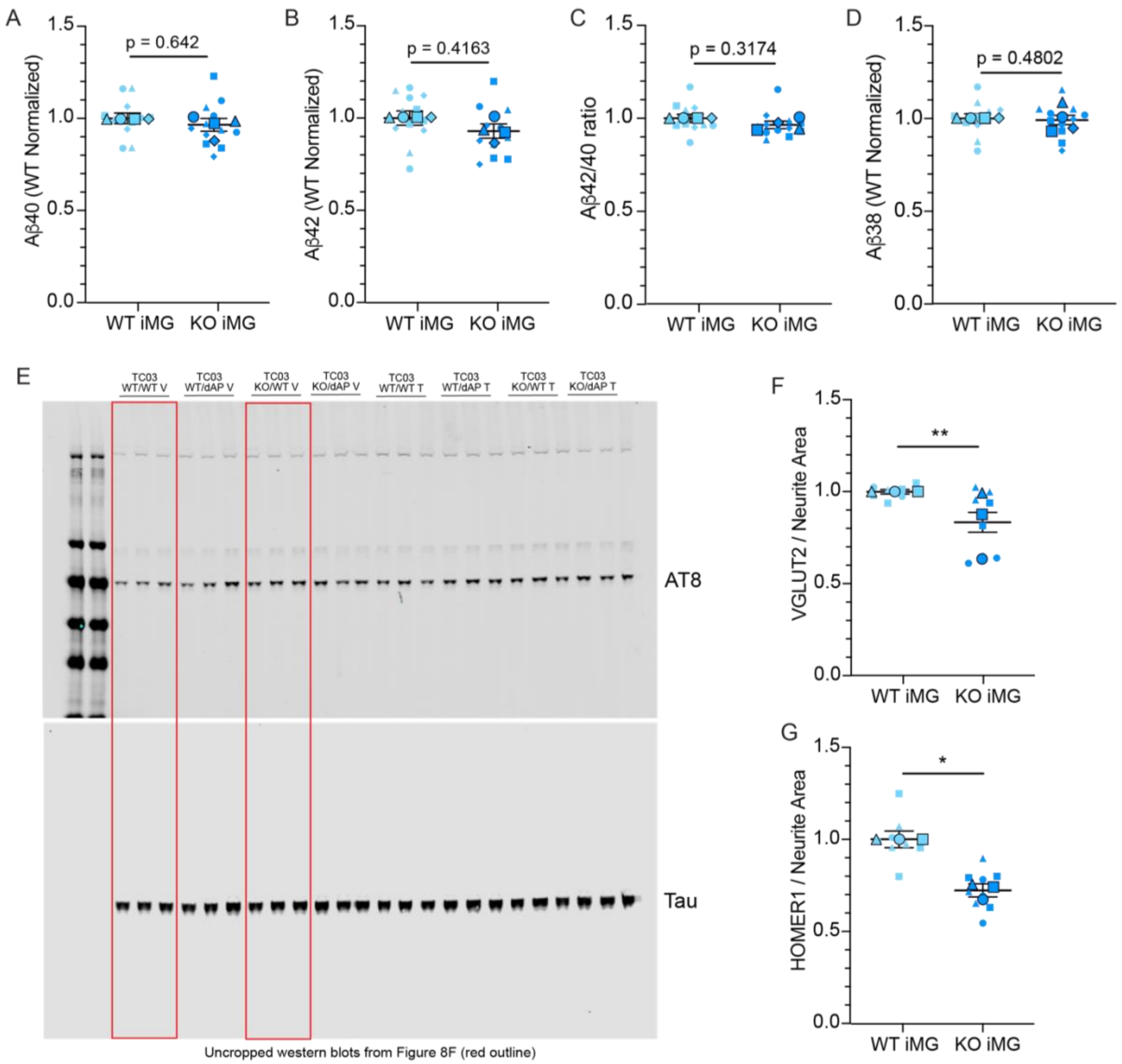


**Figure S8: Supplemental data corresponding to Figure 8**.
(A-D) Secreted Aβ40 (A), Aβ42 (B), ratio of secreted Aβ42/40 (C) and secreted Aβ38 (D) protein levels measured in conditioned media of tri-cultures with ABCA7-WT and -KO iMG. Results were normalized to ABCA7-WT iMG tri-cultures within each differentiation for visualization.
(E) Uncropped western blot corresponding to Figure 8D. Red outlined areas mark regions shown in the corresponding figure.
(F-G) VGLUT2+ (F) and HOMER1+ (G) area normalized to TUJ1+ neurite area analyzed using ICC analysis in tri-cultures with ABCA7-WT or -KO iMGs. Results were normalized to ABCA7-WT iMG tri-cultures within each differentiation for visualization.
For all scatter plots, large, outlined points are representative of differentiation average and small points represent well-level replicates within differentiation. Different shapes denote separate differentiation. Statistical analyses display results of mixed-effects ANOVA of raw data to capture within- and between-differentiation variance. Significant effect of genotype was determined: $p<0.05$ = *, $p<0.01$ = **, $p<0.001$ = ***, $p<0.0001$ = ****. Error bars = mean +/- SEM.

**SUPPLEMENTAL DATA**

**Table S1:** DIA-MS proteomics of ABCA7-WT and -KO iAs: Tab1 contains Raw MS2 values; Tab2 contains batch-corrected MS2 values.

**Table S2:** DIA-MS proteomics of ABCA7-WT and -KO iNs: Tab1 contains Raw MS2 values; Tab2 contains batch-corrected MS2 values.

**Table S3:** DIA-MS proteomics of ABCA7-WT and -KO iMGs: Tab1 contains Raw MS2 values; Tab2 contains batch-corrected MS2 values.

**Table S4:** ABCA7-KO vs -WT DEPs identified through LIMMA analysis of DIA-MS proteomics. Tab1 contains iA DEPs; Tab2 contains iN DEPs; Tab3 contains iMG DEPs.

**Table S5**: All significantly enriched pathways identified in GSEA preranked analysis of ABCA7-KO versus -WT iMG proteomics. Tab1 contains input ranked gene list; Tab2 contains positively enriched pathways; Tab3 contains negatively enriched pathways.

**Table S6**: Lipidomics analysis of BR33 ABCA7-WT and -KO iMGs. Raw and processed data are included in “Tab 1” and “Tab 2”. Remaining tabs include statistics, visualized values, and source data used to generate all plots including lipidomic analyses.

**Table S7**: DIA-MS proteomics of BR103 rs3752246-G/G and -C/C iMGs. Tab 1 contains raw MS2 values; Tab2 contains batch-corrected MS2 values; Tab3 contains peptide-level ABCA7 MS2 values.

**Table S8**: BR103 rs3752246-C/C versus -G/G DEPs identified through LIMMA analysis of DIA-MS proteomics and ranked GSEA gene list. Tab1 contains DEP gene list and logFC values identified through LIMMA analysis; Tab2 contains GSEA input ranked gene list.

**Table S9**: Results of GSEA preranked analysis of Human Proteostasis Network (HPN) gene sets. Tab1 contains GSEA of HPN gene sets on BR103 iMG DEPs; Tab2 contains GSEA of HPN gene sets on ABCA7-KO vs. -WT iMG DEPs.
